# Reversal of pulmonary veno-occlusive disease phenotypes by inhibition of the integrated stress response

**DOI:** 10.1101/2023.11.27.568924

**Authors:** Amit Prabhakar, Rahul Kumar, Meetu Wadhwa, Prajakta Ghatpande, Jingkun Zhang, Ziwen Zhao, Carlos O. Lizama, Bhushan N. Kharbikar, Stefan Gräf, Carmen M. Treacy, Nicholas W. Morrell, Brian B. Graham, Giorgio Lagna, Akiko Hata

## Abstract

Pulmonary veno-occlusive disease (PVOD) is a rare form of pulmonary hypertension arising from EIF2AK4 gene mutations or mitomycin C (MMC) administration. The lack of effective PVOD therapies is compounded by a limited understanding of the mechanisms driving the vascular remodeling in PVOD. We show that the administration of MMC in rats mediates the activation of protein kinase R (PKR) and the integrated stress response (ISR), which lead to the release of the endothelial adhesion molecule VE-Cadherin in the complex with Rad51 to the circulation, disruption of endothelial barrier, and vascular remodeling. Pharmacological inhibition of PKR or ISR attenuates the depletion of VE-Cadherin, elevation of vascular permeability, and vascular remodeling instigated by MMC, suggesting potential clinical intervention for PVOD. Finally, the severity of PVOD phenotypes was increased by a heterozygous *BMPR2* mutation that truncates the carboxyl tail of BMPR2, underscoring the role of deregulated BMP signal in the development of PVOD.

## Introduction

Pulmonary veno-occlusive disease (PVOD)/pulmonary capillary hemangiomatosis is a rare condition that represents a subgroup of patients with pulmonary hypertension (PH)^1^. PVOD is a progressive disease characterized by the involvement of pulmonary vein (PV) and/or pulmonary capillary (PC) in addition to pulmonary artery (PA)^1, 2^. The vascular remodeling in PVOD includes structural remodeling of post-capillary small veins in the lung, resulting in intimal thickening, luminal occlusion and thrombosis, fibrous intimal proliferation of septal veins and preseptal venules associated with dilatation and proliferation of PC^1^. PVOD is caused by biallelic loss-of-function/expression (LOF/E) mutations in the *EIF2AK4* gene, which encodes general control nonderepressible 2 (GCN2), a critical regulator of the eukaryotic translation initiation factor 2 (eIF2)^3, 4^. Hence, PVOD appeared clearly distinct from PAH, which had been linked to monoallelic LOF/E mutations in genes involved in bone morphogenetic protein (BMP) signaling, such as *GDF2, BMPR2, ACVRL1, ENDOGLIN, SMAD1,* and *SMAD9* ^5, 6^. However, a recent study has found that 8.5% of pulmonary arterial hypertension (PAH) patients carry biallelic LOF/E mutations in *EIF2AK4*, similarly to PVOD patients^7^. Conversely, some PVOD patients carry *BMPR2* mutations, similar to those found in PAH patients^8, 9^. The genetic overlap between PVOD and PAH indicates that the genetic analysis is not sufficient for a definitive diagnosis of the disease. This is a significant clinical roadblock because PVOD patients are at a high risk of developing pulmonary edema, respiratory failure, and even death from PAH-specific therapies^1^. On the other hand, PVOD might share a common etiology with PAH, and it might be possible to identify a therapeutic approach beneficial to both diseases. The mechanism(s) underlying the pathogenesis of PVOD, however, especially in regard to the effect of BMPR2 mutations, are not fully understood.

Earlier research indicate that cancer patients treated with chemotherapy drugs, such as mitomycin C (MMC), bleomycin, and cisplatin, are at a heightened risk of developing PVOD^10–12^. These drugs, known as alkylating agents, cause DNA damage and are cytotoxic^13^. When MMC was administered to rats, they developed a range of cardiovascular phenotypes similar to those seen in PVOD patients, such as right ventricular (RV) hypertrophy and remodeling in the pulmonary vasculature^11, 14–16^. Thus, studying the effects of MMC in rats could help us better understand the pathogenesis of PVOD and discover potential treatments. The involvement of DNA damage in the pathogenesis of PAH has been suggested on the basis of the higher level of DNA damage detected in the lungs and remodeled arteries of patients and in animal models of PAH^17^. Furthermore, risk factors associated with PAH, such as oxidative stress and inflammation, also induce DNA damage^13, 17^. It is shown previously that the DNA repair enzyme Rad51 rapidly decreased after MMC treatment or depletion of BMPR2, which resulted in the accumulation of DNA damage in human pulmonary microvascular endothelial cells (PMVECs) and in the vascular endothelium of PAH patients and mouse models^18^, suggesting that the loss of Rad51 triggered pulmonary vascular remodeling through a hitherto unknown mechanism.

The integrated stress response (ISR) is an evolutionary conserved adaptive intracellular mechanism involved in the maintenance of homeostasis upon changes in the cellular environment and pathological stimuli^19^. Maladaptive ISR activation, however, is linked to various diseases and ageing^19^. Together with GCN2, three other eIF2 kinases sense distinct types of stress and become active^20^: heme-regulated inhibitor kinase (HRI), double-stranded RNA-activated protein kinase (PKR), and PKR-like endoplasmic reticulum kinase (PERK); upon activation, these kinases phosphorylate serine-51 (Ser^51^) of the alpha subunit of eukaryotic initiation factor 2 (eIF2α)^21^. GCN2 is activated by amino acid deprivation or UV irradiation, while PKR is activated by viral infection^20^. When of eIF2α is phosphorylated at Ser^51^ (p-eIF2α), it associates with eIF2B, a guanine nucleotide exchange factor for eIF2. This association inhibits eIF2B activity and attenuates cap-dependent translation of most mRNAs^19^. Simultaneously, it promotes translation of specific transcripts, such as cyclic AMP-dependent transcription factor 4 (ATF4), a master transcriptional regulator for activating the ISR pathway^21^. Because PVOD patients carry biallelic LOF/E mutations in the *EIF2AK4* gene encoding GCN2, it is plausible to speculate that the pathogenesis of PVOD is mediated by deregulation of eIF2 and global protein synthesis; however, it has not been explored previously and the etiology of PVOD remains largely unknown^3, 4^.

In this study, we report that Rad51 interacts with vascular endothelial-cadherin (VE-Cad, also known as Cdh5), an endothelial-specific adhesion molecule^22, 23^, thereby protecting it from degradation. Upon MMC treatment in rat, Rad51 and VE-Cad are depleted from the pulmonary vascular endothelium, leading to increased permeability and loss of barrier function. Exposure to MMC triggers the induction and activation of PKR, resulting in increased phosphorylation of eIF2α and ISR activation. However, when we administered inhibitors of ISR or PKR, the secretion of the <u>V</u>E-Cad/<u>R</u>ad51 <u>c</u>omplex (hereafter referred to as VRC) is inhibited and the barrier function of the vascular endothelium as well as the pulmonary vascular phenotypes are restored. Thus, the evidence we present suggests that inhibition of the PKR-ISR axis might be effective in preventing or ameliorating the pathogenesis of PVOD.

## Results

### Fbh1 mediates Rad51 degradation after MMC treatment

A study has shown that upon MMC treatment, Rad51, an enzyme essential for DNA repair, rapidly decreases in PVMECs in a proteasome-dependent manner, which leads to DNA damage^18^. To identify an E3 ubiquitin ligase that conjugates ubiquitin to Rad51 and mediates its degradation, small inhibitory RNAs (siRNAs) targeting candidate E3 ubiquitin ligases—including Fbh1 (also known as Fbxo18), Nedd4-1 (also known as Nedd4), Nedd4-2 (also known as Nedd4L), and Smurf1—were transfected into PMVECs, followed by an immunoblot analysis of Rad51 (**Fig. 1a**). Non-specific control siRNA (siCtrl) or siRNA for Rad51 (siRad51) was transfected as controls (**Fig. 1a**). Seventy % reduction of Fbh1 by siRNA (siFbh1) resulted in a 4.5-fold increase in Rad51 compared to siCtrl cells, while the depletion of other E3 ubiquitin ligases did not significantly alter Rad51 (**Fig. 1a**). When increasing amounts of Fbh1 were expressed in PMVECs, Rad51 protein was dose-dependently decreased, supporting that Fbh1 leads to Rad51 degradation (**Fig. 1b**). Conversely, when Fbh1 was depleted with siRNA (siFbh1), Rad51 was no longer diminished after MMC treatment (**Fig. 1c**). A time-dependent change in the amount of Fbh1 and Rad51 after MMC treatment showed that Rad51 decreased as early as 2 h and kept declining for up to 14 h, while Fbh1 gradually increased (**Fig. 1d**), further confirming an inverse correlation between Fbh1 and Rad51 protein amount. We also noted the reduction of endothelial adhesion protein VE-Cad along with Rad51 (**Fig. 1d**). Both Fbh1 and Rad51 resided in the cytoplasm and the nucleus (**Fig. 1e, left**). After MMC treatment, the quantity of Fbh1 in the cytoplasm and nucleus increased 3.1-fold and 3.6-fold, respectively (**Fig. 1e, right**). This resulted in a decrease of Rad51 levels to 38% in the cytoplasm and 23% in the nucleus (**Fig. 1e, right**). The fraction of damaged nuclei, which were identified as an elongated comet-like shape by an alkaline comet assay, increased from 3-5% to 25% after MMC treatment (**Fig. 1f, siCtrl, white arrows**). This is consistent with the result of an induction of phosphorylated histone H2AX (γH2AX), a marker of DNA damage, after MMC treatment (**Fig. 1c, siCtrl**). When 70% of Fbh1 was depleted by siFbh1 (**Fig. 1a**), PMVECs were protected from DNA damage after MMC treatment (**Fig. 1f**). When Rad51 was depleted by siRad51 (**Fig. 1a**), the comet assay (**Fig. 1f, siRad51**) and γH2AX (**Fig. 1c, siRad51**) showed that the level of DNA damage was elevated even without MMC treatment. Exogenous expression of Rad51 in PMVECs prevented DNA damage (γH2AX) mediated by MMC (**Supplementary Fig. S1**). These findings demonstrate that MMC mediates DNA damage through Fbh1-dependent degradation of Rad51.

**Fig. 1.**
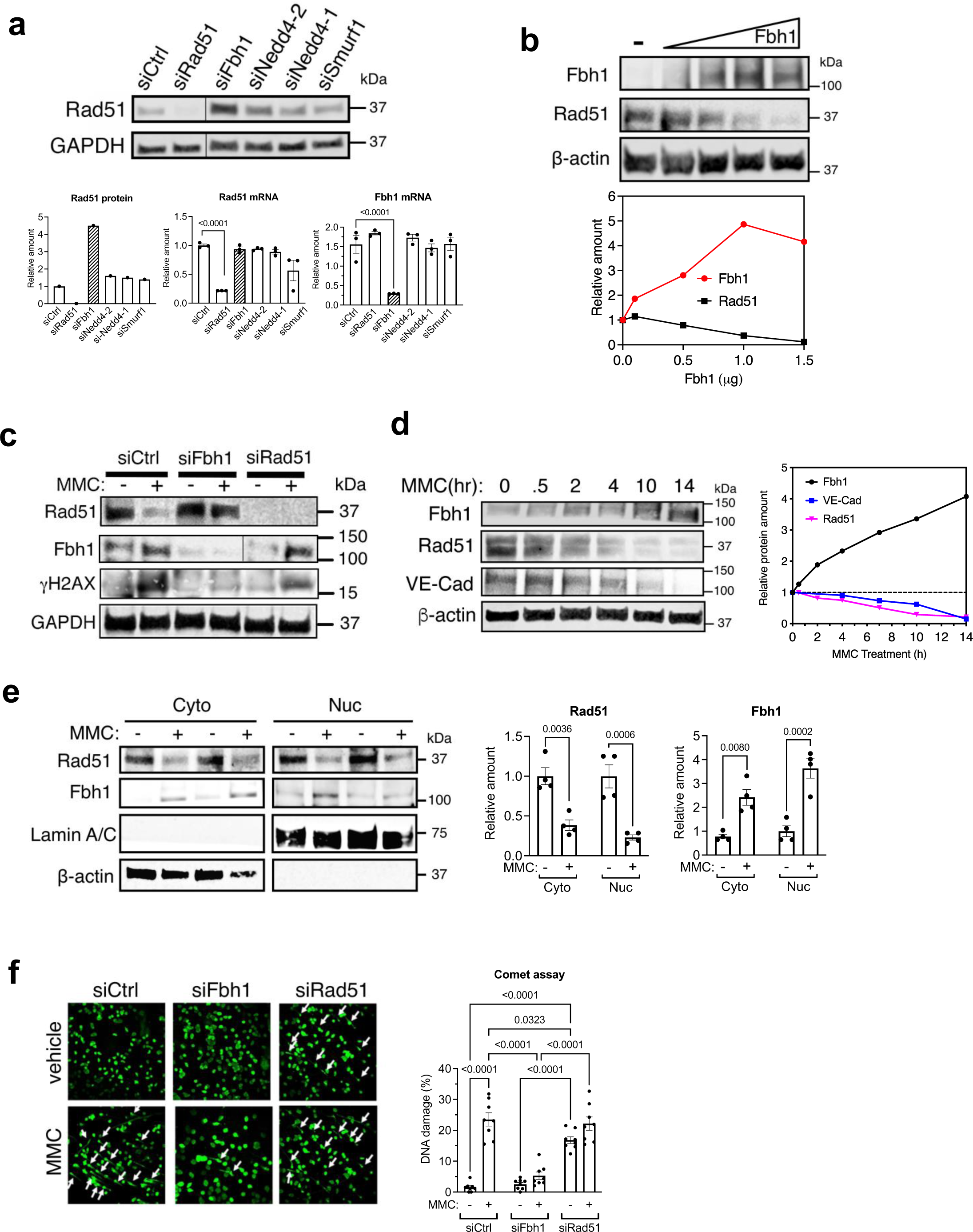
E3 ubiquitin ligase Fbh1 mediates Rad51 degradation upon MMC treatment. **a.** PMVECs were transfected with siRNA targeting Rad51, Fbh1, Nedd4-1, Nedd4-2, Smurf1 or non-specific control siRNA (siCtrl) and the total cell lysates and total RNAs were subjected to immunoblot analysis of Rad51 and GAPDH (loading control) (top). Rad51 protein amount normalized to GAPDH and the levels of Rad51 and Fbh1 mRNAs normalized to GAPDH are shown (bottom). The levels of mRNAs are shown as mean±SEM. n=3 independent RNA samples (1×10^6^ cells per sample). **b.** PMVECs were transfected with empty vector or increasing amounts of Fbh1 expression plasmid (0.1, 0.5, 1.0, or 1.5 μg) and the total cell lysates from 1×10^6^ cells were subjected to immunoblot analysis of Fbh1, Rad51, and β-actin (loading control) (top). The amount of Fbh1 and Rad51 normalized to β-actin are shown (bottom). **c.** Total cell lysates from 1×10^6^ cells were prepared from vehicle or MMC treated PMVECs after the transfection of siCtrl, siFbh1, or siRad51 and subjected to immunoblot of γH2AX, Rad51, Fbh1, GAPDH (loading control). **d.** Time-course changes in the amounts of Rad51, Fbh1 and VE-Cad following MMC treatment were examined by immunoblot using 1×10^6^ cells (left). The amounts of proteins, normalized to β-actin, are shown (right). **e.** The cytoplasmic (Cyto) and nuclear (Nuc) fraction of PMVECs treated with vehicle (saline) or MMC for 14 h were subjected to immunoblot analysis of Rad51, Fbh1, Lamin A/C (loading control for nuclear fraction) and β-actin (loading control for cytoplasmic fraction) (left). The relative amount of Rad51 and Fbh1 in Cyto and Nuc compartments was quantitated and shown as mean±SEM (right). n=4 independent samples (1×10^6^ cells per sample). **f.** Alkaline comet assay was performed in PMVECs treated with vehicle (mock) or MMC for 14 h-after the transfection of siCtrl, siFbh1, or siRad51. Representative confocal images are shown (left). White arrows indicate DNA with double strand breaks. The bar graph indicates the levels of DNA damage as a percentage of nuclei with damaged DNA out of total nuclei as mean±SEM (right). Approximately one hundred nuclei per condition were examined. n=3 independent samples (5×10^3^ cells per sample).

### Role of the MMC-Fbh1-Rad51-VE-Cad axis in the regulation of endothelial permeability

Next, the role of the MMC-Fbh1-Rad51 axis on the adherence junction (AJ) protein VE-Cad, which is integral to the junctional structure and endothelial barrier function ^22^, was examined by immunofluorescence (IF) staining. VE-Cad staining reduced to 24% after MMC treatment (**Fig. 2a, left, siCtrl**). However, when Fbh1 was depleted, VE-Cad was not diminished by MMC (**Fig. 2a, left, siFbh1**). When Rad51 was depleted, VE-Cad was reduced to 12 % of that in siCtrl cells (**Fig. 2a, left, siRad51**). The level of VE-Cad in cells transfected with both siFbh1 and siRad51 was as low as that in siRad51 cells (**Fig. 2a, left, siFbh1+siRad51**), confirming that Rad51 is the target of Fbh1 that plays a role in the control of VE-Cad. Similarly, the staining of another endothelial adhesion protein ZO-1 (zona occludens-1, also known as TJP1) decreased to 11.1% upon MMC treatment (**Fig. 2a, right, siFbh1**), but when Rad51 was stabilized by siFbh1, ZO-1 did not diminish following MMC treatment (**Fig. 2a, right, siFbh1**). Furthermore, the exogenous expression of Rad51 prevented the depletion of VE-Cad by MMC (**Supplementary Fig. S1**). These results suggest a critical role of the Fbh1-mediated degradation of Rad51 in the depletion of endothelial junctional proteins in PMVECs when treated with MMC.

**Fig. 2.**
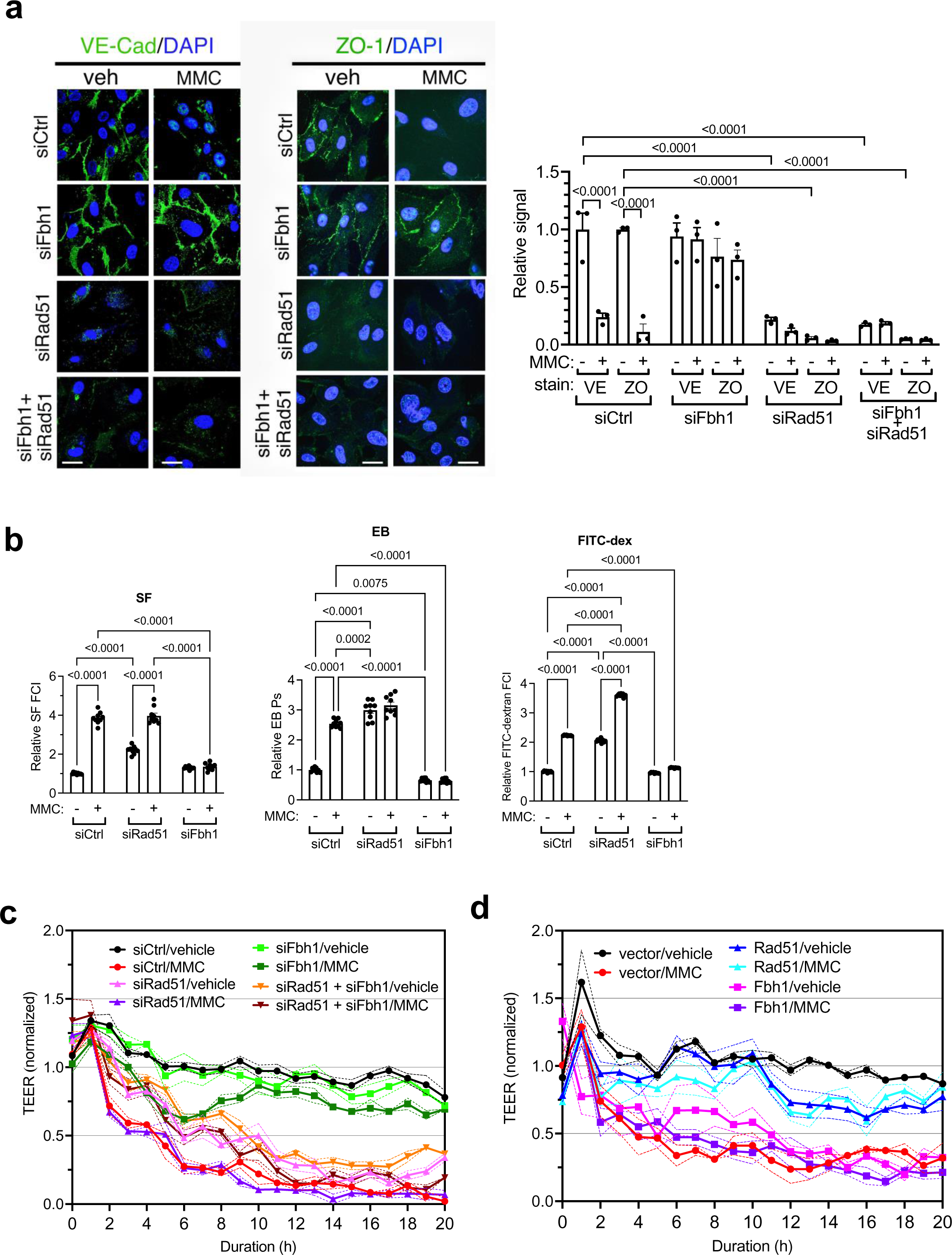
MMC treatment increases the permeability and impairs the endothelial barrier. **a.** PMVECs transfected with siCtrl, siFbh1, siRad51, or siFbh1+siRad51 were treated with vehicle (none) or MMC for 14 h, followed by IF staining of VE-Cad and ZO-1 (green) (left). Cell nuclei were stained with DAPI (blue). Scale bar= 25 μm. The intensity of IF signal of VE-Cad (VE) and ZO-1 (ZO) relative to vehicle-treated, siCtrl transfected cells is shown as mean±SEM (right). n=3 independent samples (5×10^3^ cells per sample). **b.** PMVECs transfected with siCtrl, siFbh1, or siRad51 were treated with vehicle or MMC for 14 h and subjected to TVP assay using molecular tracers: sodium fluorescein (SF), Evans blue dye (EB), or FITC-dextran (FITC-dex). The concentration of the tracer was presented as fluorescence intensity (FCI) or permeability coefficient (Ps) (mean±SEM). n=3 independent samples (5×10^3^ cells per sample). **c.** PMVECs transfected with siCtrl, siFbh1, siRad51, or siFbh1+siRad51 were treated with vehicle or MMC and subjected to the measurement of TEER (Ω cm^2^) every hour. The results are shown as mean±SEM. n=3 independent samples (5×10^3^ cells per sample). **d.** PMVECs were transfected with empty vector, Rad51 or Fbh1 expression plasmid. After the treatment with vehicle or MMC, TEER was measured every hour and the relative TEER values are shown as mean±SEM. n=3 independent samples (5×10^3^ cells per sample).

To examine the effect of the Fbh1-Rad51-VE-Cad axis on vascular permeability, PMVECs were transfected with siCtrl, siRad51, or siFbh1, then treated with or without MMC for 14 h. They were subjected to a transwell vascular permeability (TVP) assay with three molecular tracers with different molecular weights (MW): sodium fluorescein dye (SF; MW ∼0.4 kDa), Evans Blue dye (EB; MW ∼1 kDa), and FITC-dextran (FITC-dex; MW ∼40 kDa) (**Fig. 2b**) ^24^. We found an increase in the permeability of PMVECs treated with MMC or siRad51, based on the data on all three dyes (**Fig. 2b**). Again, Fbh1 depletion by siRNA, which prevents the decrease of Rad51 (**Fig. 1a**, **siFbh1**) and VE-Cad (**Fig. 2a, siFbh1**), averted the increase in permeability after MMC treatment (**Fig. 2b**), thus, confirming that the Fbh1-Rad51-VE-Cad axis plays a role in the maintenance of the barrier function in PMVECs. Finally, PMVECs were transfected with siCtrl, siRad51, siFbh1, or siFbh1+siRad51, then cultured in the presence or absence of MMC. The barrier function was examined by measuring the trans-endothelial electrical resistance (TEER) (**Fig. 2c**). A rapid decline in TEER was observed in siCtrl cells after MMC treatment (**Fig. 2c, red**), indicative of an impaired endothelial barrier. In contrast, untreated siCtrl cells retained a high TEER (**Fig. 2c, black**), indicative of an intact EC barrier. siRad51 cells that were not treated with MMC showed a TEER decline similar to that of MMC-treated siCtrl cells (**Fig. 2c, pink**), which was further reduced upon MMC treatment (**Fig. 2c, purple**). However, when Fbh1 was depleted by siFbh1, the TEER remained high upon exposure to MMC (**Fig. 2c, dark green**). This suggests that the endothelial barrier remained intact even after MMC treatment because Rad51 cannot be degraded by Fbh1. Co-transfection of siFbh1 and siRad51 abolished the protection against MMC-induced barrier damage, indicating the critical role of Fbh1-dependent Rad51 degradation (**Fig. 2c, brown**). Exogenous expression of Fbh1 was sufficient to damage the endothelial barrier without MMC treatment (**Fig. 2d, pink**). Like cells depleted in Fbh1 (**Fig. 2c, dark green**), the TEER remained high even after MMC treatment in PMVECs overexpressing Rad51 (**Fig. 2d, light blue**), suggesting that the presence of Rad51 provides protection against MMC-induced barrier damage. The results suggest that the MMC-Fbh1-Rad51 axis not only impairs genome integrity, but also causes damage to EC-EC junctions and EC barrier. These findings shed light on the crucial role of the MMC-Fbh1-Rad51 axis in endothelial injury.

### Formation of a Rad51–VE-Cad complex in vascular endothelial cells

We performed a proximity ligation assay (PLA) to detect the putative colocalization of endogenous Rad51 and VE-Cad in PMVECs *in situ* ^25^. A robust PLA signal (red dots) appeared in untreated PMVECs (**Fig. 3a, left, top panels**) but not detected in the absence of primary antibodies (**Fig. 3a, left, bottom panels**), suggesting that Rad51 and VE-Cad colocalize in PMVECs. MMC treatment reduced the number of PLA signal to 17% (**Fig. 3a**), reflecting the MMC-induced depletion of Rad51 (**Fig. 1d**) and VE-Cad (**Fig. 2a**). Immunoprecipitation (IP) with an anti-Rad51 antibody followed by immunoblot with an anti-VE-Cad antibody also detected a VRC (VE-Cad/Rad51 complex) in untreated PMVECs, which was reduced to 16% by MMC treatment (**Fig. 3b**). In addition, colocalization of VE-Cad and Rad51 at the endothelial junctions was confirmed by IF staining (**Fig. 3c**). A mapping of the cytoplasmic domain (Cyto) of VE-Cad [amino acid (aa) 621-789], which associates with Rad51 using deletion mutants (Prox, Mid, Dist, and Δ5) revealed that Rad51 interacted with aa 715-730 of VE-Cad which overlapped between the Mid and Dist mutants (**Fig. 3d, red arrow**). The Rad51-interacting region (aa 715-730) does not overlap with those regions known to associate with other VE-Cad associating proteins, such as VE-PTP, p120-catenin, and β-catenin,^26^ but it contains one of three microtubule-associated protein light chain 3 (LC3) interacting motifs (LC3 motif) ^27, 28^ (**Fig. 3d, green box**). Cadherin-6 (Cdh6), another member of the cadherin-superfamily of proteins, contains four LC3 motifs that associate with the autophagic apparatus GABARAP and lead to the lysosomal degradation of Cdh6 ^27, 28^. We speculate that the three LC3 motifs additively increase the association of VE-Cad with Rad51, as the Cyt, Dist, and Δ5 mutants, which contain three LC3 motifs, associated with a higher amount of Rad51 than the Mid mutant, which contains only one LC3 motif (**Fig. 3d)**. Phosphorylation of eIF2 and activation of ISR are implicated in the induction of autophagy^29^. A time-dependent conversion of a cytosolic form of the LC3 protein (LC3-I) to the phosphatidylethanolamine-conjugated form (LC3-II) was observed as early as 2 h after MMC treatment (**Fig. 3e**), indicating activation of autophagy upon MMC treatment. The association of Rad51 with the LC3 motifs of VE-Cad might have a role in protecting VE-Cad from autophagosome-mediated proteolysis.

**Fig. 3.**
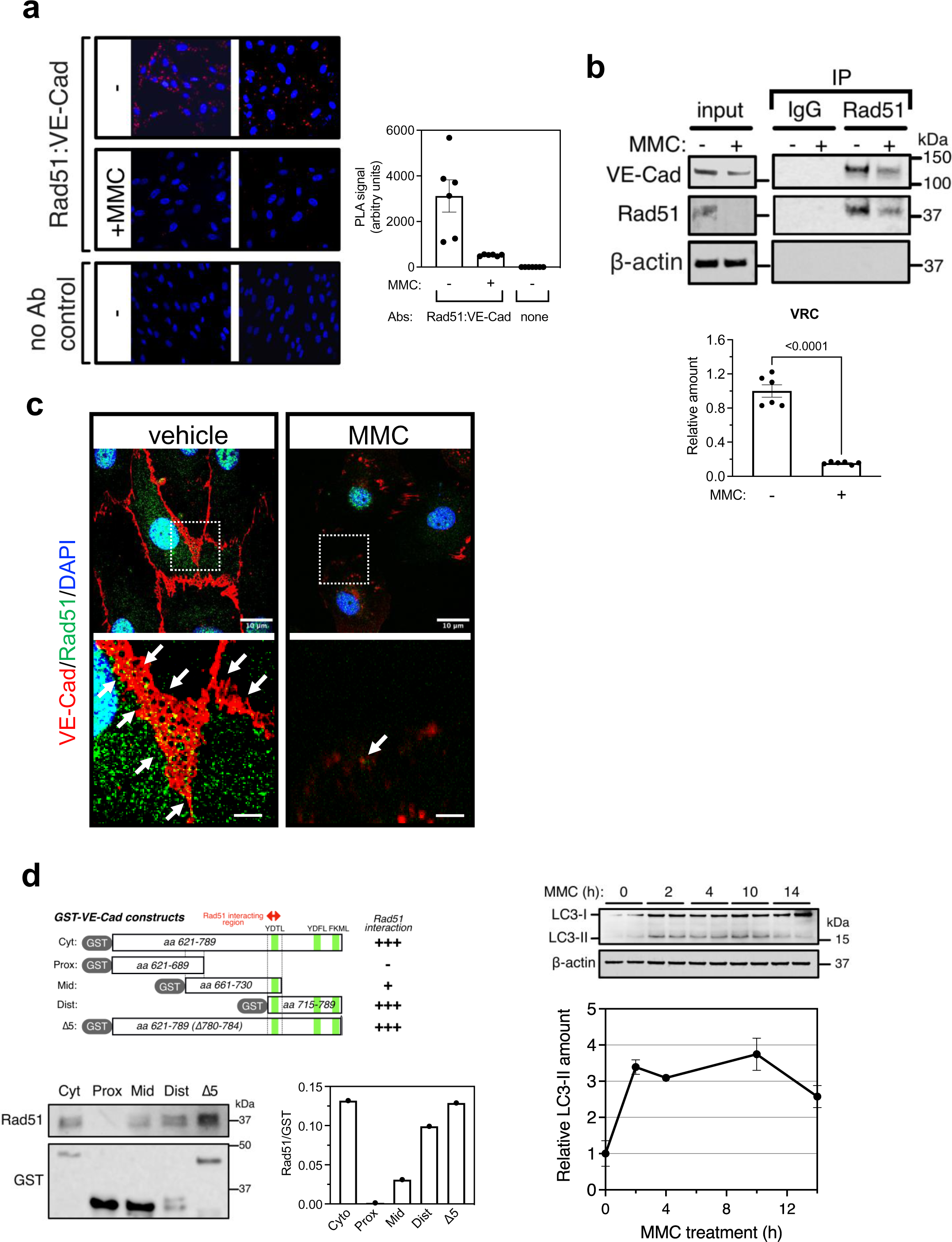
Interaction of Rad51 and VE-Cad in PMVECs. **a.** The images of in situ PLA using an anti-Rad51 and anti-VE-Cad antibodies in PMVECs treated with vehicle or MMC for 14 h (left). The images of samples without primary antibodies (PLA probes only) are shown as control (No Abs). PLA signal (red dots) represent the colocalization of VE-Cad and Rad51. DAPI was used to stain the nuclei. The PLA signal was quantitated and shown as mean±SEM (right). n=3 independent samples (5×10^3^ cells per sample). **b.** PMVECs treated with vehicle or MMC for 14 h were subjected to immunoprecipitation (IP) by non-specific IgG or an anti-Rad51 antibody, followed by immunoblot analysis of VE-Cad, Rad51, and β-actin (loading control) (top, IP). The total cell lysates without IP were also subjected to immunoblot (top, input). The relative amount of VRC is shown as mean±SEM (bottom). n=6 independent samples (1×10^6^ cells per sample). **c.** PMVECs treated with vehicle or MMC for 14 h were subjected to IF staining of VE-Cad (red) and Rad51(green). Cell nuclei were stained with DAPI (blue). The area indicated by the white box (top) is magnified and displayed at the bottom. Arrows indicate endothelial junctions where VE-Cad and Rad51 colocalize (yellow). Light blue signal is due to the nuclear localization of Rad51. Scale bars= 10 μm (top) and 40 μm (bottom). **d.** A schematic representation of VE-Cad cytoplasmic domain (Cyt; aa 621-789) and four deletion mutants (Prox, Mid, Dist, and Δ5) that are as GST-fusion proteins (top). The amount of Rad51 pulled down with GST-fusion proteins was analyzed by immunoblot (bottom left). The ratio of Rad51 bound by GST-fusion protein and the amount of GST-fusion protein (Rad51/GST) is shown as a bar graph (bottom right) and summarized as +/-(top). Rad51 interaction region and LC3 motifs are indicated in red arrow and green boxes, respectively. **e.** Time course conversion of LC3-I form of LC3 protein to LC3-II form and β-actin (control) after MMC treatment was examined by immunoblot (top). The relative amount of LC3-II normalized to β-actin is shown as mean±SEM (bottom). n=2 independent samples (1×10^6^ cells per sample).

### Release of a VRC to the extracellular compartment

The detection of a VRC at a basal state in PMVECs led us to investigate its behavior upon MMC treatment. We hypothesized that upon MMC treatment, a VRC might be released from PMVECs into the extracellular space, leading to their depletion in PMVECs. We collected the conditioned media (CM) from PMVECs treated with either vehicle or MMC for 14 h, followed by immunoblot. We detected a 7-fold and 2.7-fold increase in VE-Cad and Rad51 in the CM after MMC treatment, respectively (**Fig. 4a**). Enzyme-linked immunoassay (ELISA) demonstrated the rapid detection of both proteins in the CM 2 h after MMC treatment, with their levels gradually increasing in a time-dependent manner (**Fig. 4b**). To determine whether these proteins were released as a complex, we immunoprecipitated Rad51 from the CM of PMVECs treated with or without MMC, followed by immunoblot with an anti-VE-Cad antibody. VRC was detected in the CM from untreated cells, and the amount of VRC in the CM was increased 1.7-fold after MMC treatment (**Fig. 4c**), complementing the depletion of VRC inside the cells upon MMC treatment (**Fig. 3a and 3b**). Notably, the molecular size of the VE-Cad in the CM was 130 kDa and was indistinguishable from the intracellular VE-Cad, eliminating the possibility that VE-Cad detected the CM upon MMC treatment is a product of proteolytic cleavage, as previously reported in endothelial cells undergoing apoptosis^30^.

**Fig. 4.**
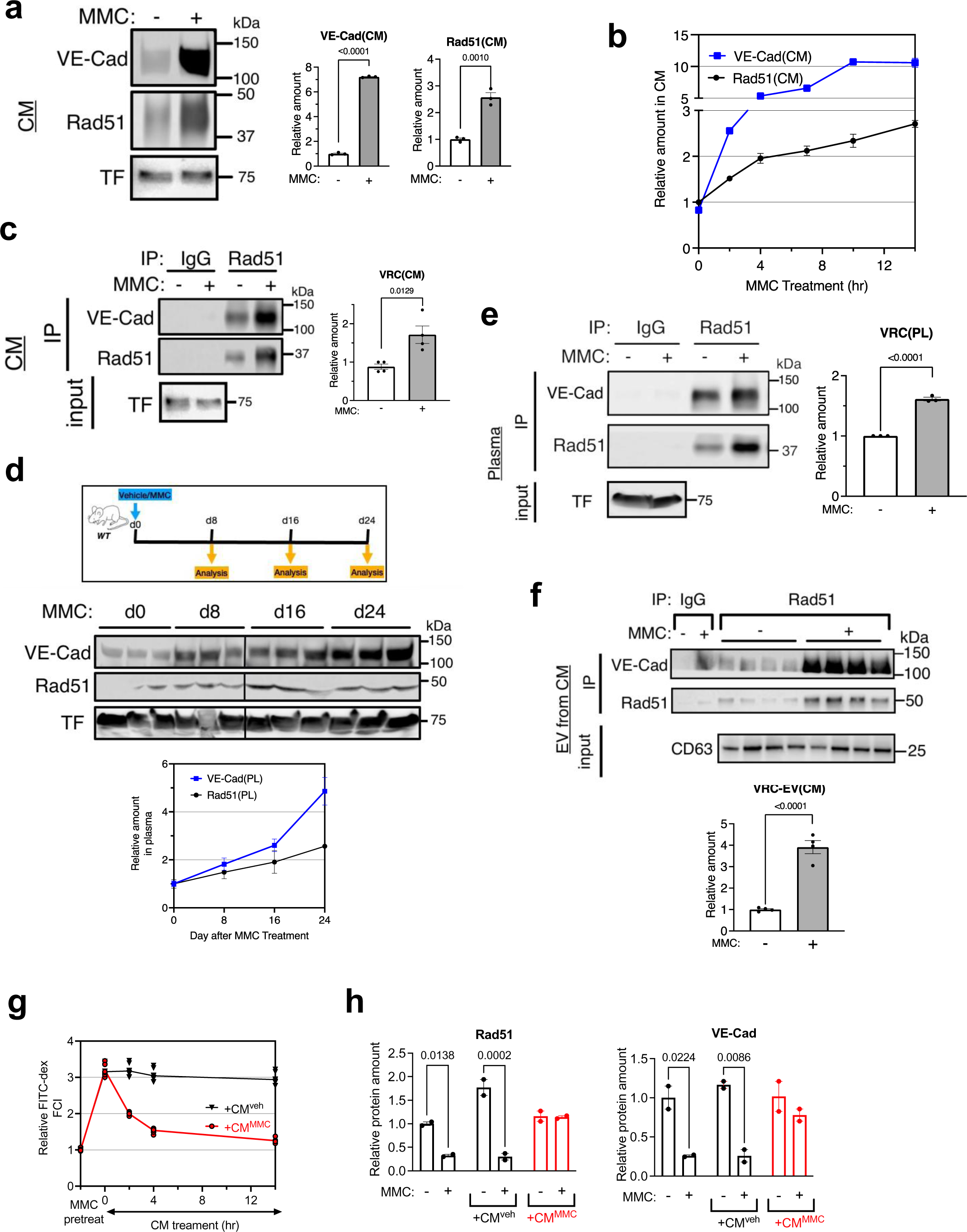
Secretion of VE-Cad/Rad51 complex from the vascular endothelial cells. **a.** Conditioned media (CM) were collected from PMVECs treated with vehicle or MMC for 14 h and subjected to immunoblot with an anti-VE-Cad, anti-Rad51, and anti-Transferrin (TF) antibody (loading control) (left) and its quantitation (right). n=3 independent samples (1×10^6^ cells per sample). **b.** A time-course change in the amount of VE-Cad and Rad51 in the CM of PMVECs after MMC treatment (0-14 h) were measured by ELISA and plotted as mean±SEM. n=3 independent samples (1×10^6^ cells per sample). **c.** CM from PMVECs treated with vehicle or MMC for 14 h were subjected to IP with non-specific IgG (control) or an anti-Rad51 antibody, followed by an anti-VE-Cad (for VRC) and anti-Rad51 antibody. Immunoblot with an anti-TF antibody is shown as loading control. n=4 independent samples (1×10^6^ cells per sample). **d.** A scheme of MMC treatment in rats (top). The plasma samples of WT rats injected once with vehicle or MMC were collected 8, 16, and 24 days after the injection and subjected to immunoblot of VE-Cad and Rad51 (middle). Immunoblot with an anti-TF antibody is shown as loading control. The relative amount of Rad51 and VE-Cad in the plasma of vehicle or MMC (d8,d16, and d24) treated rats were quantitated and plotted as mean±SEM (bottom) n=3 independent samples per group. **e.** The plasma of vehicle or MMC treated rat (d24) were subjected to an IP with an anti-Rad51 antibody, followed by immunoblot analysis of an anti-VE-Cad (for VRC) and anti-Rad51 antibody to detect the interaction between these proteins. Immunoblot with an anti-TF antibody (TF) is shown as loading control. n=3 independent samples. **f.** Extracellular vesicles (EV) were purified from the CM of PMVECs treated with vehicle or MMC for 14 h and subjected to IP with with non-specific IgG (control) or an anti-Rad51 antibody, followed by immunoblot with an anti-VE-Cad and anti-Rad51 antibodies to detect the interaction between these proteins (top). The amount of VE-Cad associating with Rad51 (VRC) was quantitated and shown as mean±SEM (bottom). Immunoblot with an anti-CD63 antibody, the EV marker, is shown as loading control. n=4 independent samples per condition (1×10^6^ cells per sample). **g.** The CM from PMVECs treated with vehicle (CM^veh^) and MMC for 14 h (CM^MMC^) were supplemented to the culture media of PMVECs pretreated with MMC for 2 h (5×10^3^ cells). A time-course change (0-14 h) in the permeability was measured by a TPV assay in triplicate with FITC-dex as a tracer. The result is shown as relative FCI and presented as mean±SEM. **h.** The amount of Rad51 (left) and VE-Cad (right) protein in PMVECs after the incubation in the presence of with CM^veh^ or CM^MMC^ for 14 h were examined by immunoblot and quantitated. The result is shown as mean±SEM. n=2 independent samples (1×10^6^ cells per sample).

To observe whether VRC secretion from vascular endothelial cells also occurs in vivo, we examined the time-dependent change in the amount of VE-Cad and Rad51 in the plasma of Sprague-Dawley rats administered once with 3 mg/kg MMC, which is sufficient to induce PVOD phenotypes in rats within 24 days (**Fig. 4d, top**). Immunoblot analysis revealed a time-dependent increase in both proteins in the plasma of MMC-treated rats (**Fig. 4d, middle and bottom**). Twenty-four days (d24) after MMC administration, the amount of VRC in the plasma was 1.6-fold higher in MMC-treated vs. vehicle-treated rats (**Fig. 4e**). This suggests that VRC might also be released from the vascular endothelium into circulation upon MMC treatment in vivo. Different cells, including ECs, release extracellular vesicles and particles (hereafter referred to as EV) that encapsulate proteins, lipids, and nucleic acids^31^. The nanoparticle tracking analysis (NTA) showed that PMVECs stimulated with MMC secreted 20-fold higher number of EV than that of vehicle-treated cells (**Supplementary Fig. S2a**). This result validates that MMC treatment induces the secretion of EV and the release of the VRC from PMVECs. Furthermore, the EV released from the MMC-stimulated PMVECs ranged from 100 to 290 nm in diameter and was larger than those released from control cells (**Supplementary Fig. S2b**). The EV isolated from vehicle- or MMC-treated PMVECs, which was positive for the EV marker CD63^32^, were subjected to IP with an anti-Rad51 antibody, followed by immunoblot with an anti-VE-Cad antibody (**Fig. 4f**). We detected a 4-fold higher amount of VRC in the EV derived from MMC-treated PMVECs compared to mock-treated PMVECs (**Fig. 4f, top**). Therefore, it seems that MMC promotes the release of VRC via EV, resulting the depletion of VRC and impairment of junctional structure in the endothelium.

EV not only play a role in releasing cargo into the extracellular milieu, but also in transferring it to other cells^31^. We tested whether MMC-treated ‘recipient’ ECs could uptake the VRC encapsulated in EV, which were released by ‘donor’ ECs, and reverse the damaged junctional structure and increased permeability. The CM from PMVECs treated with MMC for 14 h (CM^MMC^) was collected and replaced with the media of PMVECs pre-treated with MMC for 2 h, followed by a TVP assay using FITC-dextran. We used CM from a vehicle treated PMVECs (CM^veh^) as a control. After exposure to MMC, the permeability of the recipient cells increased ∼3-fold. However, after adding CM^MMC^, but not CM^veh^, endothelial permeability gradually returned to basal levels (and not CM^veh^) in a time-dependent manner, indicating significant repair of the endothelial barrier (**Fig. 4g, red**). Consistently, the levels of cellular Rad51 and VE-Cad, which were depleted after the pre-treatment with MMC (**Fig. 4h and Supplementary Fig. S3a**), were rescued by the addition of CM^MMC^ (**Fig. 4h, +CM^MMC^**), but not by CM^veh^ (**Fig. 4h, +CM^veh^**). IF staining of VE-Cad further confirmed the rescue of VE-Cad in MMC-treated PMVECs when supplemented with CM^MMC^ (**Supplementary Fig. S3b, +CM^MMC^**). These results support a model in which a VRC, secreted via EV, is capable of restoring the endothelial barrier and reducing permeability when it is taken up by the damaged ECs.

### Activation of the integrated stress response pathway by MMC

To dissect the genetic pathway that mediates the degradation of Rad51 via Fbh1 in response to DNA damage mediated by MMC, we performed a transcriptome analysis in PMVECs treated with MMC for 0, 4, and 14 h. An analysis of differentially expressed genes (vehicle vs MMC) detected that 2,385 ATF4 target genes (**Fig. 5a and Supplementary Table 1)**—including *ATF3, ATF5, GADD34*, *GADD45, CHOP,* and *TRIB3*—were induced by MMC treatment (**Supplementary Fig. S4a**). We also noted the induction of the p53 pathway as a result of DNA damage upon MMC treatment ^13^ (**Supplementary Fig. S4b**). As ATF4 is the main effector of the integrated stress response (ISR)^33^, this result demonstrated the activation of the ISR pathway in PMVECs exposed to MMC. qRT-PCR of lung lysates of WT rats treated with MMC verified increased levels of transcripts of ATF4 and ATF4 target gene, such as *ATF3* and *GADD34* (**Fig. 5b**), as in PMVECs (**Supplementary Fig. S4a**). We noted PKR mRNA encoded by the EIF2AK2 gene was also increased in the lung of rats exposed to MMC (**Fig. 5b**). Three consensus ATF4 binding sites (**Fig. 5c, ATF4 bs**), conserved among all vertebrate *EIF2AK2* orthologs, within the first intron of the EIF2AK2 gene. To investigate whether ATF4 regulates the EIF2AK2 gene, we performed chromatin immunoprecipitation (ChIP) assay. In unstimulated cells, we detected a basal level of association of ATF4 with *EIF2AK2.* However, MMC treatment triggered a 37-fold increase in ATF4 binding to *EIF2AK2* (**Fig. 5c, bottom**). This suggests that MMC promotes the recruitment of ATF4 to *EIF2AK2*, thereby increasing the amount of PKR mRNA (**Fig. 5b**). This result suggests that the MMC-dependent induction of PKR via ATF4 leads to the phosphorylation of eIF2α, activating the ISR cascade and generating a feedforward mechanism for persistent ISR activation.

**Fig. 5.**
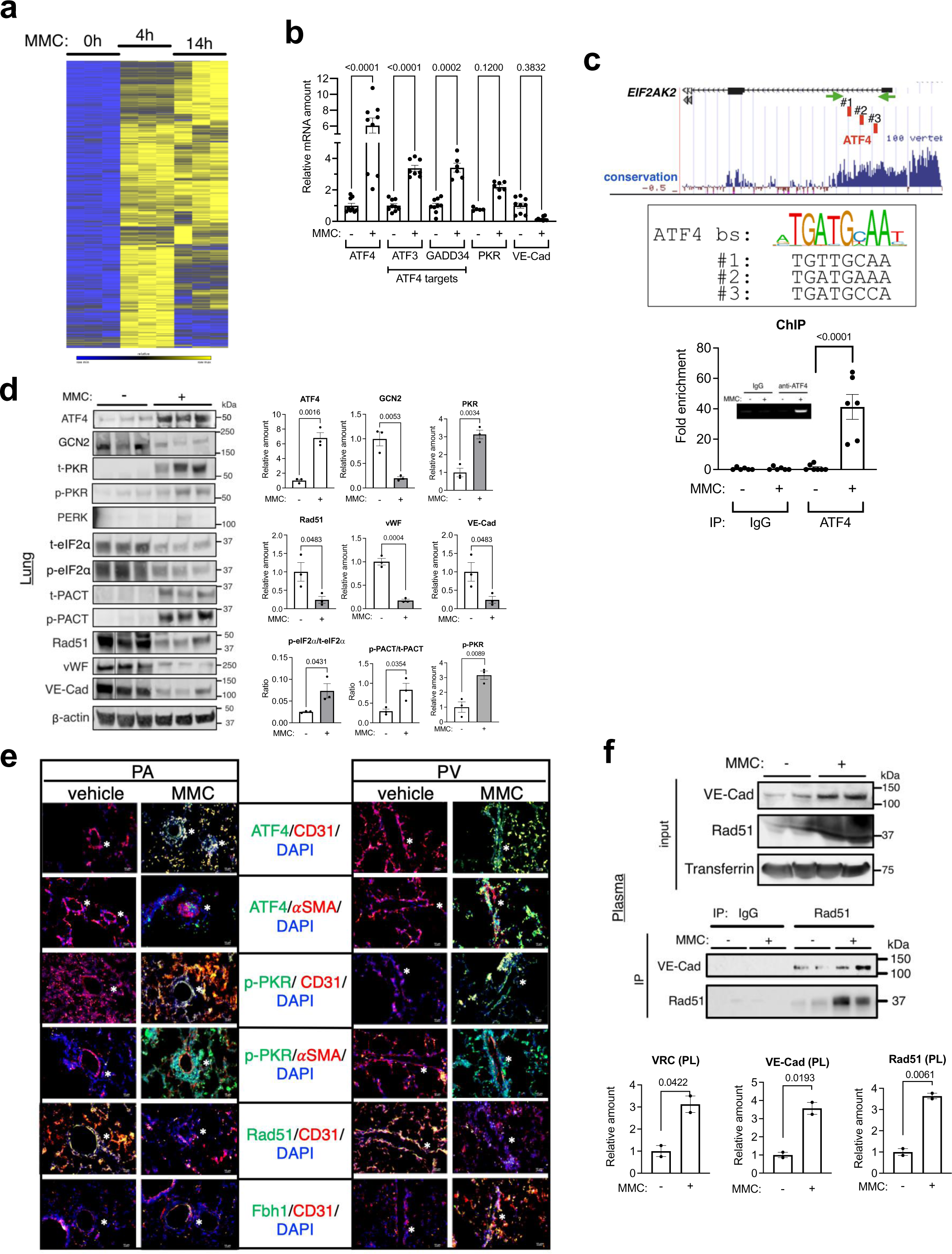
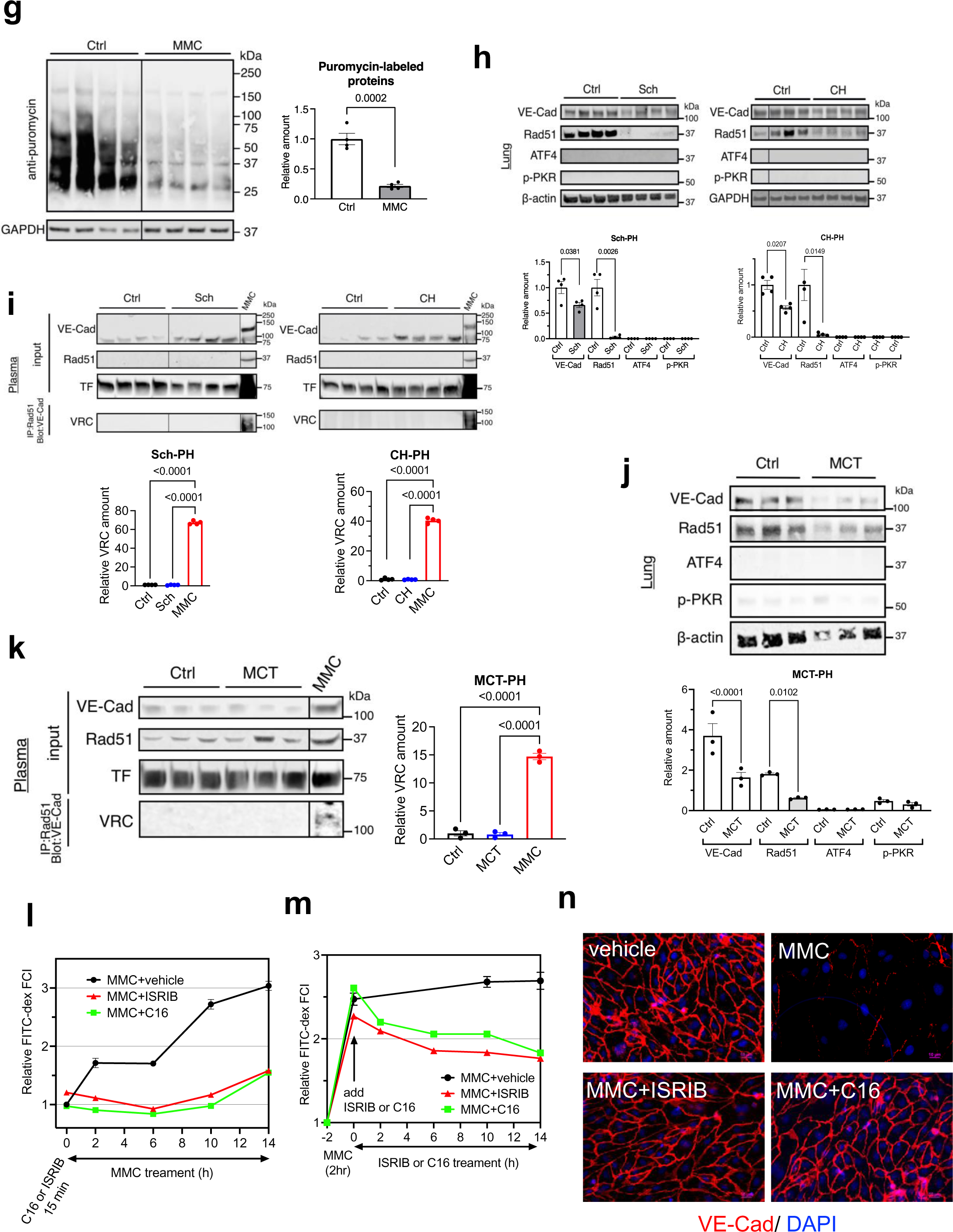
MMC activates ISR pathway via PKR induction. **a.** ATF4 target genes differentially expressed in PMVECs (1×10^6^ cells) treated with MMC for 0, 4, or 14 h are shown in a heat map. n=3 independent RNA samples per condition. **b.** Total RNAs isolated from the lungs of WT rats injected with MMC (d24) were subjected to qRT-PCR of ATF4 mRNAs, ATF4 target mRNAs (ATF3 and GADD34), PKR mRNA, and VE-Cad mRNA (control of MMC treatment) in triplicates. The results are normalized by GAPDH mRNAs and presented as mean±SEM. **c.** ChIP assay in PMVECs (1×10^6^ cells) treated with vehicle or MMC for 4 h. ChIP was performed with non-specific IgG (control) or an anti-ATF4 antibody, followed by PCR amplification using primers (green arrows) franking the genomic region of the EIF2AK2 gene containing ATF4 binding motifs (red boxes) (top). The PCR result is shown as fold-enrichment over the input sample as mean±SEM (bottom) and the PCR product image is shown as an inset. n=3 independent experiments. **d.** Lung lysates of WT rats 24 days after vehicle or MMC administration were subjected to immunoblot of the indicated proteins (left). The amount of indicated proteins normalized to β-actin, the ratio of p-eIF2α: t-eIF2α and p-PACT: t-PACT are shown as mean±SEM (right). n=3 independent samples per condition. **e.** The lungs from WT rats administered with vehicle or MMC were harvested on day 24 and subjected to IF staining with anti-ATF4 (green), anti-p-PKR (green), anti-αSMA (red), anti-CD31 (red), anti-Rad51 (green) and anti-Fbh1 (green) antibody and the images of PAs and PVs are taken, and merged images are shown. Cell nuclei were stained with DAPI (blue). Asterisk indicates the location of PA or PV. Scale bar=10 μm. **f.** Rad51 and VE-Cad in the plasma of WT rats 24 days after vehicle or MMC administration was detected by immunoblot (top). The plasma samples were also subjected to immunoprecipitation with non-specific IgG (control) or an anti-Rad51 antibody, followed by immunoblot with an anti-VE-Cad (for VRC), anti-Rad51 (middle) and anti-Transferrin (TF) antibody (loading control). The relative amount of VRC, VE-Cad, and Rad51 normalized to TF is shown as mean±SEM (bottom). n=2 per condition**. g.** In vivo puromycin incorporation assay was performed using lung lysates of WT rats (n=4 per condition) 24 days after vehicle or MMC administration. After normalized by the total protein amount, puromycin-labeled proteins were visualized by immunoblot with an anti-puromycin antibody and presented in quadruple (left). The amount of puromycin-labeled proteins normalized to GAPDH (loading control) is shown as mean±SEM (right). **h.** The amount of indicated proteins in the lung was compared between control mice and *Schistosomia* (Sch)*-*induced PH mice (top left) and chronic hypoxia (CH)-induced PH mice (top right). The amount of indicated proteins normalized to β-actin (loading control) is shown as mean±SEM (bottom). n=4 per condition. **i.** The plasma of Sch-PH (left), CH-PH mice (right), and their controls (Ctrl) were subjected to immunoblot with an anti-VE-Cad and anti-Rad51 antibody. Immunoblot with an anti-TF antibody blot is shown as loading control. The plasma samples were also subjected to immunoprecipitation with anti-Rad51 antibody, followed by immunoblot with an anti-VE-Cad antibody for the detection of VRC (top). The plasma of MMC-treated rats (MMC) was included as control. The relative amount of VRC is shown as mean±SEM (bottom). n=4 independent samples per condition**. j.** The amount of indicated proteins in the lung was compared between vehicle-(Ctrl) or MCT-administered (MCT) WT rats. The amount of indicated proteins normalized to β-actin (loading control) is shown as mean±SEM (bottom). n=3 independent samples per condition. **k.** The plasma of vehicle-(Ctrl) or MCT-administered WT rats (MCT) were subjected to immunoblot with an anti-VE-Cad and anti-Rad51 antibody (top). Immunoblot with an anti-TF antibody blot is shown as loading control (left). The plasma samples were also subjected to immunoprecipitation with anti-Rad51 antibody, followed by immunoblot with an anti-VE-Cad antibody for the detection of VRC (left). The plasma of MMC-treated rats (MMC) was included as control. The relative amount of VRC is shown as mean±SEM (right). n=3 independent samples per condition**. l.** PMVECs (5×10^3^ cells) were pre-treated with PKR inhibitor C16 or ISR inhibitor ISRIB for 15 min, followed by MMC exposure for 1-14 h. Time-course changes in the permeability was examined by a TPV assay using FITC-dex as a tracer. The results are presented as relative FCI; mean±SEM. n=4 independent samples. **m.** PMVECs (5×10^3^ cells) were pre-treated with MMC for 2h, followed by the treatment with PKR inhibitor C16 or ISR inhibitor ISRIB for 0-14 h. Time-course changes in the permeability was examined by a TPV assay using FITC-dex as a tracer. The results are presented as relative FCI; mean±SEM. n=4 independent samples. **n.** PMVECs treated with vehicle or MMC alone, or MMC and ISRIB or C16 for 14 h were subjected to IF staining with anti-VE-Cad antibody (red) and DAPI stain (blue) for nuclei. Scale bar=10 μm

To explore the role of MMC-mediated ISR activation and EC damage in vivo, we performed immunoblot analysis of the endothelial proteins [VE-Cad and von Willebrand factor (vWF)] in lung lysates from rats exposed to either vehicle or MMC. We found a depletion of these proteins, indicating obliteration of vascular endothelial cells following MMC treatment (**Fig. 5d**). Rad51 and VE-Cad were also diminished (**Fig. 5d**), as seen in PMVECs treated with MMC (**Fig. 1d**). Furthermore, ATF4 protein level was increased ∼7-fold, a hallmark of ISR activation, in MMC-treated rats (**Fig. 5d**). As ISR can be initiated by the phosphorylation of eIF2α by eIF2α kinases such as GCN2, PKR, PERK, or HRI, we also investigated their protein levels and found that PKR increased by 3.1-fold in MMC-treated rats (**Fig. 5d**). Unlike PKR, GCN2 was significantly diminished after MMC treatment, mimicking PVOD patients with LOF/E mutations in *EIF2AK4* encoding GCN2 (**Fig. 5d**). A time-course analysis showed that the depletion of GCN2 occurred as early as 8 days (d8) after MMC treatment and remained low up to 24 days (d24) (**Supplementary Fig S5**). PERK was expressed in low amounts both with and without MMC (**Fig. 5d**), and HRI is mainly expressed in erythroid cells^34^, leaving PKR as the likely mediator of eIF2α phosphorylation and ISR activation triggered by MMC. When phosphorylated at serine-246 (S^246^), the PKR activator protein (PACT) associates with PKR and activates its kinase activity^35^. We found that the level of S^2^^46^-phosphorylated PACT (p-PACT) relative to total PACT (p-PACT/t-PACT) was 2.8-fold higher in MMC-treated rats compared to vehicle-treated (control) rats (**Fig. 5d**), suggesting PKR activation by PACT in rats exposed to MMC. Furthermore, there was a 3-fold increase in the level of PKR autophosphorylation on threonine-446 (p-PKR), indicating full activation of the PKR kinase activity upon MMC treatment ^36^ (**Fig. 5d**). Consistently, we detected a 2.9-fold increase in p-eIF2α relative to total eIF2α (p-eIF2α/t-eIF2α) (**Fig. 5d**). IF staining of ATF4 or p-PKR showed the induction of p-PKR and ATF4 in the CD31-positive endothelium of PAs and PVs in MMC-treated rats (**Fig. 5e**). Decreased Rad51 staining and increased Fbh1 staining were also evident in the endothelium of PAs and PVs in MMC-treated rats (**Fig. 5e**). In contrast, the levels of VRC, VE-Cad, and Rad51 in the plasma increased 3.1-fold, 3.6-fold, and 3.6-fold, respectively, after the MMC treatment (**Fig. 5f**). The NTA revealed that the size of the EV in the plasma of MMC-treated rats ranged from 100 to 290 nm in diameter (**Supplementary Fig. S2d**), which overlapped with the EV released by MMC-stimulated PMVECs (**Supplementary Fig. S2a**). Furthermore, the quantity of EV in the plasma of MMC-treated rats was 2.5-fold higher than that that of vehicle-treated rats (**Supplementary Fig. S2c**), corroborating that MMC induces the release of VRC through EV.

Annexin V staining revealed apoptotic cells within the pulmonary vascular endothelium, indicated by positive CD31 staining, on day 8 (d8) following MMC, but not vehicle, administration (**Supplementary Fig. S6**). However, by day 16 (d16) and 24 (d24), the number of apoptotic (Annexin V-positive) cells had decreased (**Supplementary Fig. S6**). These results imply that the activation of ISR and the depletion of VRC in the pulmonary vascular endothelial cells trigger apoptosis and the obliteration of the vascular endothelium as early as the 8^th^ day after MMC treatment.

To examine whether eIF2 phosphorylation by PKR upon MMC treatment results in the inhibition of global protein synthesis, rats injected with vehicle (Ctrl) or MMC were treated with puromycin, and lung lysates were subjected to immunoblot with an anti-puromycin antibody to detect puromycin-incorporated nascent proteins. The total levels of puromycin-labeled proteins were 78% lower in MMC-injected rats than control rats, confirming global translational inhibition in the lungs of MMC-treated rats (**Fig. 5g**). Altogether, these results suggest that MMC activates PKR, phosphorylates eIF2α, and inhibits translation, leading to preferential translation of *ATF4*, transcriptional activation ATF4 target genes, and execution of the ISR.

To examine whether the activation of the PKR-ISR pathway is detectable in other classes of PH, we analyzed lung lysates from mice with Schistosomia-induced PH (Sch-PH) and chronic hypoxia-induced PH (CH-PH) ^37^. Both Sch-PH and CH-PH mice exhibited PH phenotypes, such as increased RVSP and RV/LV+S ratio (**Supplementary Fig. S7a**). Similar to MMC-treated rats, VE-Cad and Rad51 were reduced in the lungs of these mice, however, neither PKR activation (p-PKR induction) nor the induction of ATF4 was detected (**Fig. 5h**), indicating no activation of the PKR-ISR pathway in Sch-PH and CH-PH mice. Unlike the plasma of MMC-treated rats (**Fig. 5f**), neither full-length (130 kDa) VE-Cad, Rad51, nor VRC were detected in the plasma of Sch-PH or CH-PH mice (**Fig. 5i**). The extracellular domain of VE-Cad (90kDa), resulting from the proteolytic cleavage, was detected in both mouse models of PH (**Fig. 5j**). We also examined the lung and plasma samples from Monocrotaline (MCT) induced rat PH model (MCT-PH). MCT-PH rats exhibited an increase in RVSP and RV/LV+S ratio, hallmarks of PH (**Supplementary Fig. S7b**). The lungs lysates of MCT-PH rats demonstrated no induction of p-PKR or ATF4 compared to control rats (**Fig. 5j**), indicating no activation of the PKR-ISR pathway in MCT-PH rats compared. Like the plasma of Sch-PH and CH-PH mice, VRC was not detected in the plasma of MCT-PH rats unlike MMC-mediated PVOD rats (**Fig. 5k**). These results suggest that ISR activation and the release of VE-Cad and Rad51 into circulation are unique to PVOD model animals.

To investigate whether the effects of MMC on endothelial dysfunction could be mitigated by pharmacological intervention, we used the PKR antagonist C16 [6,8-Dihydro-8-(1*H*-imidazol-5-ylmethylene)-7*H*-pyrrolo[2,3-*g*]benzothiazol-7-one; IC_50_=210 nM] ^38^ or the ISR inhibitor ISRIB [trans-N,N’-(Cyclohexane-1,4-diyl)bis(2-(4-chlorophenoxy)acetamide; IC_50_=5 nM] ^39, 40^ to precondition PMVECs for 15 min, followed by MMC treatment and a TVP assay (**Fig. 5l**). As observed previously, MMC treatment resulted in 3-fold increase in permeability by 14 h (**Fig. 5l, black**), but this effect was reduced to ∼50% when cells were pre-treated with C16 or ISRIB (**Fig. 5l, green and red**). In contrast, when PMVECs were first treated with MMC for 2 h to impair the endothelial barrier, and then exposed to C16 and ISRIB, cell permeability decreased by ∼35% compared to vehicle-treated cells in a time-dependent manner (**Fig. 5m, green or red vs black**). Furthermore, IF staining of VE-Cad indicated that cotreatment with MMC and ISRIB, or C16 prevented VE-Cad depletion and preserved the integrity of endothelial junctions (**Fig. 5n**). These findings suggest that inhibiting the PKR-mediated activation of ISR prevents the endothelial barrier damage caused by MMC. This highlights the importance of examining the effects of these inhibitors on the pulmonary vascular remodeling induced by MMC in rats.

### ISRIB prevents vascular remodeling mediated by MMC in rats

MMC treatment in humans and rats causes PVOD^14^. Because the ISR pathway is activated in MMC-treated rats, we hypothesized that the administration of the ISR inhibitor ISRIB might be able to prevent the MMC-induced PVOD phenotype in rats^41, 42^. Since *BMPR2* mutations have been found in some PVOD patients^8, 9^, we also tested the effect of MMC and ISRIB in *BMPR2* mutant (Mut) rats, in which monoallelic deletions of 2 nucleotides (nt) (Δ2; E503fs) or 26 nt (Δ26; Q495fs) were introduced in the BMPR2 gene by TALEN-mediated genome editing (**Supplementary Fig. S8a**); both frameshift alleles express truncated proteins lacking the carboxyl terminal tail domain (CTD) of BMPR2 (ΔCTD-BMPR2) and resembling *BMPR2* mutations found in PAH patients^43–45^. Immunoblot analysis showed both full-length BMPR2 (FL-BMPR2) (**Supplementary Fig. S8b**) and ΔCTD-BMPR2 in Mut rats (**Supplementary Fig. S8c**). Administration of MMC did not alter the amount of BMPR2-FL (**Supplementary Fig. S8c**). Since homozygous *BMPR2* mutation (both Q495fs and E503fs) are embryonic lethal, all the Mut *BMPR2* rats used in this study were heterozygotes. When pulmonary artery smooth muscle cells (PASMC) derived from WT and Mut (E503fs) rats were stimulated with increasing concentrations of BMP4, the phosphorylation of Smad1/5/8 (p-Smad1/5/8) was indistinguishable between WT and Mut PASMC (**Supplementary Fig. S9a**). Upon BMP4 stimulation, changes over time in p-Smad1/5/8 (**Supplementary Fig. S9b, top**) and in the mRNAs of Smad1/5/8 target genes—such as *Id3* and *Smad6*—were similar between WT and Mut PASMC (**Supplementary Fig. S9b, bottom**). Similar results were obtained with another BMP ligand BMP7 (**Supplementary Fig. S9c**), indicating that the BMP-BMPR-Smad signaling pathway is intact in Mut rats. These results underscore the essential role of the Smad-independent signaling pathway(s) linked to the BMPR2 CTD during vascular remodeling in PAH and embryogenesis ^43–46^.

Eight to nine weeks old WT and Mut (both male and female) rats were subjected to a single injection of vehicle or MMC (3 mg/kg) on day 0 (d0) with ISRIB (0.25 mg/kg) or vehicle, followed by ISRIB treatment three times a week until d24 (total 9 injections), and an analysis of cardiopulmonary phenotypes on d24. When lung extracts from WT or Mut rats treated with MMC/ISRIB were subjected to immunoblot analysis, we found that the ISRIB treatment prevented the induction of ATF4 and PKR autophosphorylation (p-PKR) by MMC in both WT and Mut rats, indicating an attenuation of the PKR-ISR pathway (**Fig. 6a).** Depletion of Rad51 and VE-Cad by MMC was also prevented by ISRIB treatment (**Fig. 6a**). Furthermore, ISRIB treatment prevented the depletion of GCN2 induced by MMC, indicating that the ISR activation by MMC mediates the depletion of GCN2 (**Supplementary Fig. S5**). The level of eIF2α phosphorylation, as indicated by the p-eIF2α /t-eIF2α ratio, was not altered by ISRIB treatment in response to MMC (**Supplementary Fig. S10a and S10b**). This is consistent with previous reports, which demonstrate that ISRIB attenuates the ISR without inhibiting eIF2α phosphorylation, but rather by affecting eIF2B conformation ^39, 47^.

**Fig. 6.**
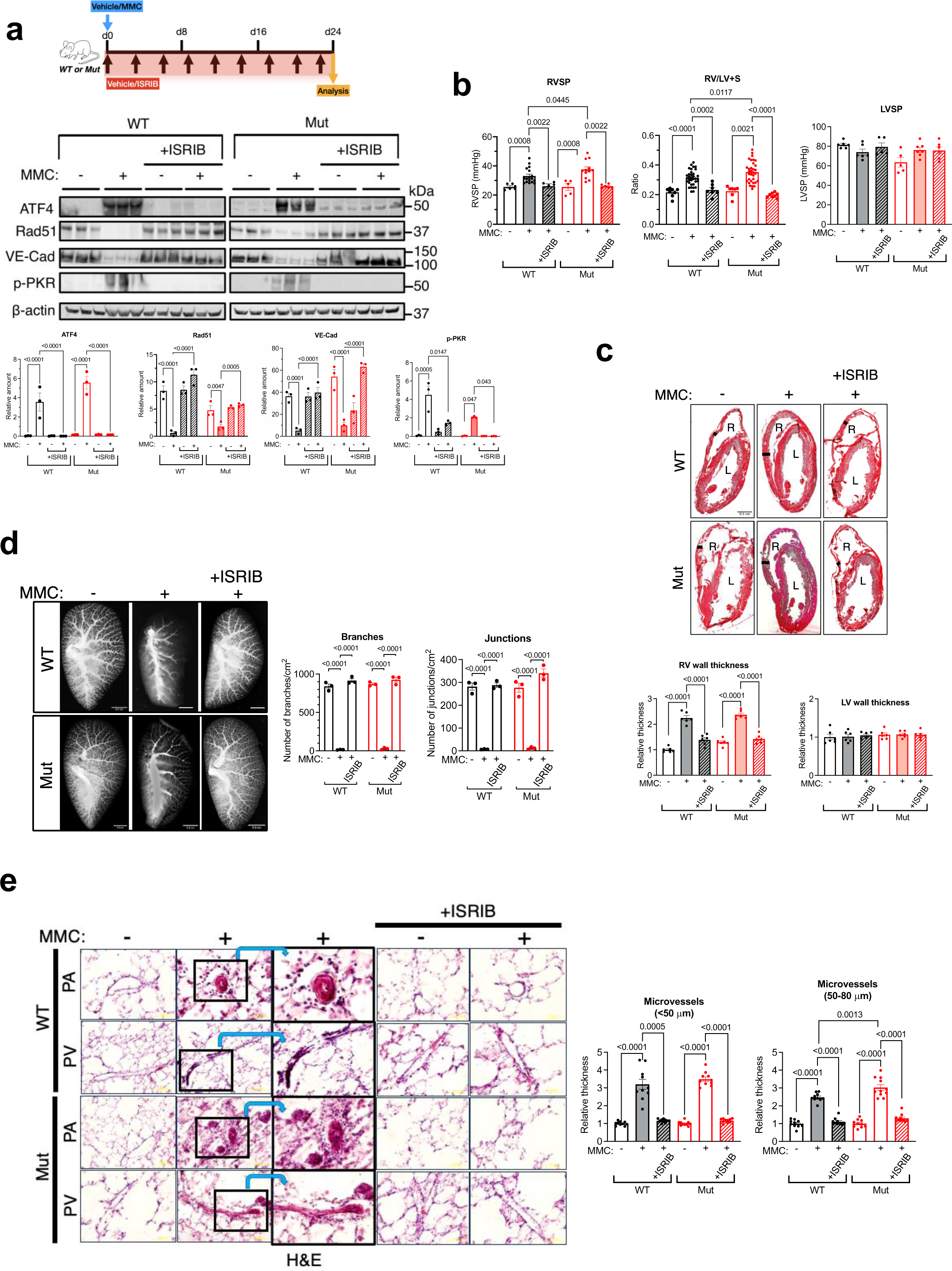
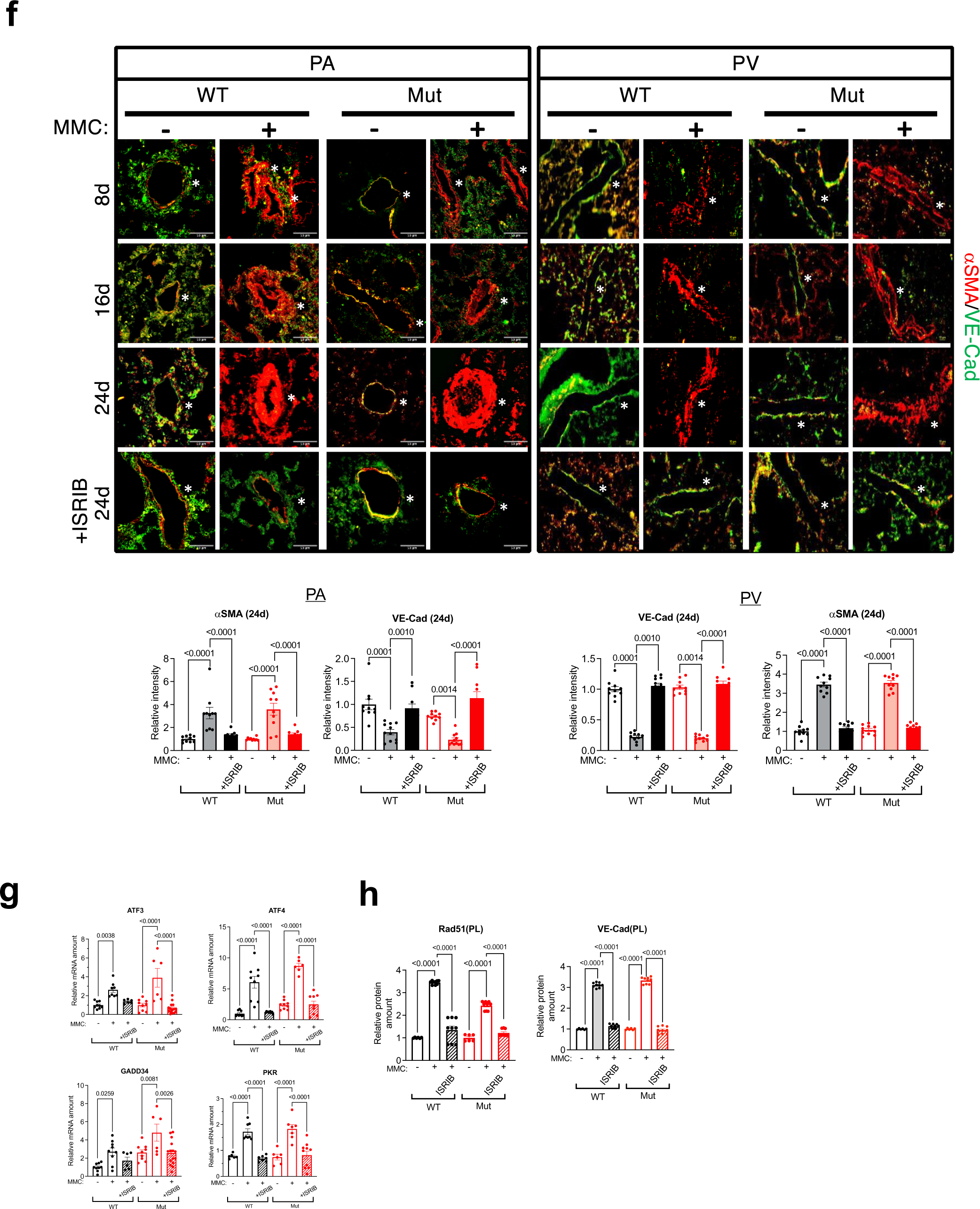
Inhibition of the ISR prevents MMC induced PVOD in WT and Mut rats. **a.** A scheme of ISRIB treatment in MMC rats (top). Immunoblot analysis of indicated proteins in total lung lysates from vehicle (-), MMC (+) with or without ISRIB in WT and Mut rats (middle). The amount of indicated proteins relative to β-actin are shown as mean+SEM (bottom). n=3 independent samples. **b.** RVSP (mmHg), RV/LV+S ratio, and LVSP (mmHg) in control and MMC-exposed rats (MMC). n=8∼34 independent samples. **c.** H&E staining of heart from vehicle-, MMC-, or MMC+ISRIB-treated WT and Mut rats (top). R and L indicate right and left ventricle, respectively. A black bar is used to indicate RV wall thickness. RV and LV wall thickness was measured by ImageJ at three different locations per sample and shown as mean±SEM (bottom). n=6 independent samples. **d.** Pulmonary vasculature of WT and Mut rats treated with vehicle or MMC with or without ISRIB was casted by Microfil on day 24 (d24), and holistic images of a lobe are shown as black and white images (left). Scale bar= 0.5 cm. A number of branches and junctions (per cm^2^) of distal pulmonary vessels was counted and shown as mean±SEM (right). n=3 independent samples per group. **e.** H&E staining of pulmonary vasculature (PA and PV) in WT and Mut rats treated with vehicle or MMC with or without ISRIB was performed on d24 and shown (left). The third column is a magnified image of the black rectangle area in the second column (left). The medial thickness of microvessels (<50μm diameter) and medium sized vessels (50-80μm diameter) was quantitated, converted to the value relative to the medial thickness of vehicle-treated WT rats, and shown as mean+SEM (right). Images are acquired at 20X magnification. n=10 independent samples. **f.** PA (left) and PV (right) from WT and Mut rats 8, 16, or 24 days after vehicle or MMC administration were stained with an anti-αSMA (red) and anti-VE-Cad (green) antibody for smooth muscle cells and endothelial cells, respectively (top). The fourth row represents 24 days MMC+ISRIB treated condition. Asterisk indicates the location of PA or PV. Scale bar=50 μm. The IF signals of d24 images were quantitated, converted to the value relative to the signal intensity of vehicle-treated WT rats, and shown as mean+SEM (bottom). n=10-11 independent samples. **g.** The level of mRNAs of ATF4 target genes in the lung of from WT and Mut rats administered with vehicle, MMC, or MMC+ISRIB was analyzed by qRT-PCR and shown as mean±SEM. n=3 independent samples. **h.** The amount of Rad51 and VE-Cad in the plasma of rats treated with MMC with or without ISRIB was quantitated by ELISA and is shown as mean±SEM. n=3 independent samples.

We next analyzed several pathological traits observed in PAH and/or POVD patients, as well as in their corresponding animal models, to monitor the effects of the *BMPR2* mutation and the inhibition of the ISR. The PA pressure [right ventricular systolic pressure (RVSP)] and RV hypertrophy, measured by right ventricle RV/LV + septum (S) weight ratio (RV/LV+S) of WT and Mut rats at the basal state was indistinguishable [25.6 mmHg in WT and 25.5 mmHg in Mut (for RVSP) and 0.216 in WT and 0.224 in Mut (for RV/LV+S)], indicating no sign of spontaneous PH development in Mut rats as reported on the rat carrying a heterozygous loss-of-expression *BMPR2* mutation ^48^ (**Fig. 6b**). When WT and Mut rats were administered with MMC, RVSP was elevated by 46% (to 33.1 mmHg) in WT rats and by 55 % (to 37.6 mmHg) in Mut rats and RV/LV+S ratio was elevated by 41% (to 0.313) in WT rats and by 52% (to 0.352) in Mut rats (**Fig. 6b**). These results indicate Mut rats develop more severe PH phenotypes compared to WT rats upon MMC treatment. No change in the left ventricular systolic pressure (LVSP) was detected after MMC treatment (**Fig. 6b**). With the co-treatment with ISRIB and MMC, the RVSP in both WT and Mut rats decreased to the levels (26.0 mmHg in WT and 26.1 mmHg in Mut) similar to those in vehicle-treated WT and Mut rats (**Fig. 6b**). Similarly, the RV/LV+S ratio was decreased to 0.232 and 0.196 with the cotreatment with ISRIB and MMC in WT and Mut rats, respectively, which are similar to vehicle-treated WT and Mut rats (**Fig. 6b**). RV hypertrophy was determined by 2.2-fold and 2.4-fold increase in the wall thickness of the RV in WT and Mut rats treated with MMC, respectively, which was prevented by ISRIB (**Fig. 6c**). There was no change in LV wall thickness in WT and Mut rats treated with MMC (**Fig. 6c**). Together with the result of LVSP (**Fig. 6b**), these results confirm the administration of MMC does not induce LV hypertrophy. Both male and female rats developed a similar severity of pulmonary hypertension phenotypes (RVSP and RV/LV+S) after MMC treatment (**Supplementary Fig. S11**). The lung weight (**Supplementary Fig. S12a**) and the lung weight/ body weight ratio (**Supplementary Fig. S12b**) were also elevated after MMC treatment by 43% and 66% (lung weight) and 63% and 137% (lung/body) in WT and Mut rats, respectively and ISRIB treatment prevented these effects. The lung weight in Mut rats became higher than that in WT rats after MMC treatment (**Supplementary Fig. S12a**), similar to the results for RVSP and RV/LV+S (**Fig. 6b**). We noted that ISRIB treatment partially rescued body weight loss after the administration of MMC (**Supplementary Fig. S12c**). No change in the weight of the liver, heart, and kidney was detected upon MMC or ISRIB treatment (**Supplementary Fig. S12d-f**). Unlike in the lung, the vascular morphology in the heart and liver remained unchanged (**Supplementary Fig. S13a**). Furthermore, ATF4 induction (**Supplementary Fig. S13b**) and protein synthesis inhibition (**Supplementary Fig. S13c**) were not detected in the heart and liver after MMC treatment, demonstrating no ISR activation in these organs. Thus, at least in this model, the *BMPR2* mutation appeared to exacerbate PVOD phenotypes (**Fig. 6a and 6b**), and ISRIB prevented the development of PVOD in Mut rats similar to WT rats.

The images of the Microfil-casted pulmonary vessels indicated a decrease in the density of the distal pulmonary vessels in MMC-treated WT and Mut rats (**Fig. 6d, left**). Counting the number of branches and junctions in the distal pulmonary vessels revealed a significant reduction in both after MMC treatment and this reduction was prevented by ISRIB treatment (**Fig. 6d, right**). When pulmonary vasculature was perfused with EB dye in MMC-treated WT and Mut rats, the extravasation of EB dye stained the entire lung blue (**Supplementary Fig. S14a**) and the intensity of EB stain increased ∼1.7-fold compared to that in vehicle-treated WT and Mut rats (**Supplementary Fig. S14b**). This is due to the impaired endothelial barrier and increased vascular permeability as a result of MMC treatment. However, when WT or Mut rats were co-treated with ISRIB, the extravasation of EB was prevented, and the appearance of the lung resembled that in the control WT and Mut rats (**Supplementary Fig. S14a**). Moreover, the intensity of the EB stain in the ISRIB-treated rats was indistinguishable from that in the control rats (**Supplementary Fig. S14b**), demonstrating that the endothelial barrier and vascular permeability remained intact when WT or Mut rats were treated with ISRIB. Furthermore, MMC-treated rats developed medial thickening in PAs and PVs, as reported previously ^11, 14^ (**Fig. 6e, left**). Both microvessels (<50 μm in diameter) and medium-sized vessels (50-80 μm in diameter) underwent remodeling in WT and Mut rats following MMC treatment (**Fig. 6e, right**). However, the medial thickness of the medium-sized vessels (50-80 μm) increased 3-fold after MMC treatment in Mut rats, while in WT rats, it only increased 2.5-fold (**Fig. 6e, right**). This suggests that more severe vascular remodeling is induced in Mut rats with *BMPR2* mutation upon the administration of MMC. Consistent with the physiological parameters (**Fig. 6b**), vascular remodeling in Mut rats was more severe than in WT rats, but ISRIB attenuated the vascular phenotype in Mut and WT rats equally (**Fig. 6e**). IF staining of PAs (**Fig. 6f, left**) and PVs (**Fig. 6f, right**) with VE-Cad, a vascular endothelial cell marker, and α-smooth muscle actin (αSMA), a smooth muscle marker, showed that the depletion of endothelial cells and the growth of smooth muscle cells advanced in a time-dependent manner after MMC treatment in WT and Mut rats (**Fig. 6f**). By 24 days post-MMC treatment, VE-Cad signals had dropped to 30-40%, while αSMA signals had increased to 300% in comparison to vehicle-treated WT and Mut rats (**Fig. 6f**). However, these MMC-mediated vascular changes were prevented by ISRIB (**Fig. 6f**). After MMC administration, the levels of Rad51 and VE-Cad in the plasma of Mut rats were higher than in WT rats (**Supplementary Fig. S15)**, which is consistent with a more severe vascular phenotype in Mut rats. The vascular phenotypes of two Mut lines (Q495fs and E503fs) treated with MMC were indistinguishable (**Supplementary Fig. S16**). Altogether, these results demonstrate that ISRIB effectively blocks both the vascular remodeling and the impairment of endothelial barriers mediated by MMC in rats, even on a *BMPR2-*Mut background.

The protective role of ISRIB against pulmonary vasculopathy in rats seem to be through its inhibition of ISR pathway, as the induction of PKR mRNA and several mRNAs encoding other ISR pathway genes, such as *ATF3*, *ATF4*, and *GADD34,* was abolished by ISRIB in MMC-treated WT and Mut rats (**Fig. 6g**). The result of a ChIP assay further supported this, showing that the recruitment of ATF4 to the PKR gene locus in response to MMC was attenuated by ISRIB treatment (**Supplementary Fig. S17**). This confirms that ATF4 acts as a critical regulator of PKR mRNA upon ISR activation. ELISA, which showed that, after ISRIB treatment, the amount of VE-Cad and Rad51 in the plasma of MMC-treated WT and Mut rats was reduced to the level of vehicle-treated rats (**Fig. 6h**). Therefore, in rats, an ISR inhibitor prevents both a PVOD-like phenotype and the release of the VRC into the plasma upon DNA damage stress. This further establishes a relationship among DNA damage, ISR, plasma VRC, and PVOD.

To confirm the activation of ISR by PKR in the pathogenesis of PVOD, we examined the effect of the PKR inhibitor C16 on MMC-mediated PVOD. WT rats were administered either a vehicle or MMC (3 mg/kg) on d0, and a vehicle or C16 (33.5 μg/kg) on d0, d8, and d16 (three injections), followed by the analysis of physiological parameters on d24 (**Fig. 7a).** The administration of C16 prevented the MMC-induced increase in RVSP, RV/LV+S, and lung/body weight ratio (**Fig. 7a**). Vascular remodeling, which includes medial hypertrophy (**Fig. 7b, H&E**) and fibrous growth of the adventitia (**Fig. 7b, blue stain in Trichrome**) as well as the reduction of the density of distal pulmonary vessels (**Supplementary Fig. S18**) mediated by MMC were attenuated by the administration of C16. Perfusion of the pulmonary vasculature with EB dye revealed that, like ISRIB treatment, C16 treatment prevented the increase in vascular permeability observed in MMC-treated rats (**Supplementary Fig. S14a**). As a result, the EB stain of the lung was similar to that seen in vehicle-treated rats (**Supplementary Fig. S14b**). Immunoblot analysis indicated that the MMC-mediated phosphorylation of eIF2α (p-eIF2α/t-eIF2α ratio) and the induction of PKR were both prevented by C16 (**Fig. 7c**). This validates the successful inhibition of the PKR-ATF4 axis and confirms that PKR— rather than GCN2, PERK, or HRI—is the eIF2α kinase responsible for the activation of ISR by MMC. The MMC-mediated ATF4 induction was also abolished by C16, confirming attenuation of the ISR (**Fig. 7c**). The decrease of Rad51 and VE-Cad in cells (**Fig. 7c**) and their increase in plasma (**Fig. 7d**) in response to MMC were both prevented by C16. This evidence supports a model in which PKR activates the ISR upon MMC treatment, leading to a decrease in cellular Rad51 and VE-Cad, disruption of the junctional structure in ECs, increased permeability, and vascular remodeling.

**Fig. 7.**
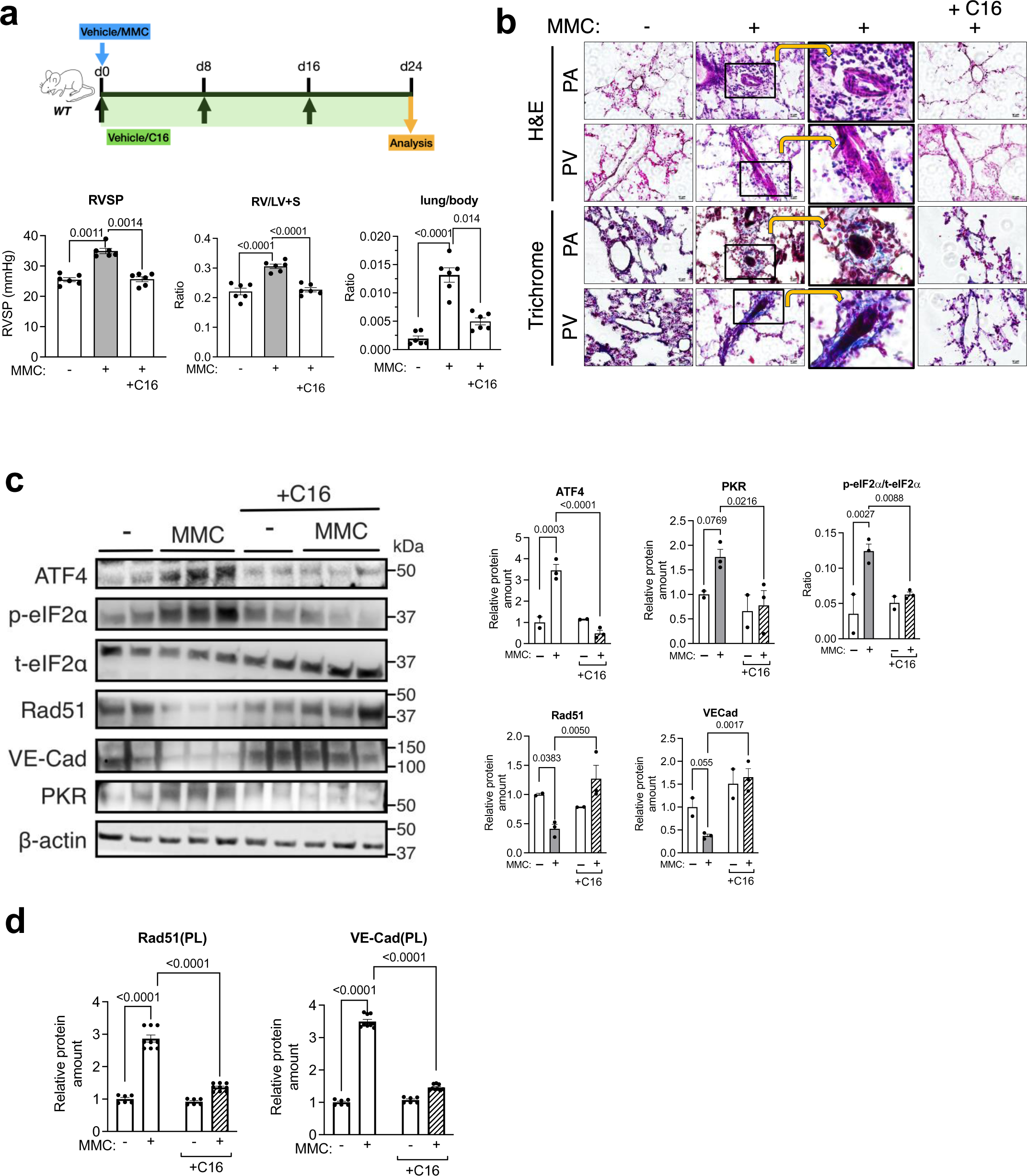
PKR antagonist C16 prevents PVOD phenotypes. **a.** A scheme of C16 treatment in WT rats. RVSP, RV/LV+S ratio, and lung/body weight ratio in vehicle- or MMC-exposed WT rats with or without C16 are shown. n=6 independent samples. **b.** H&E and Gomori’s trichrome staining of pulmonary vasculature (PA and PV) in WT rats treated with vehicle or MMC with or without C16. The third column is a magnified image of the black rectangle area in the second column. Images are acquired at 20X magnification. **c.** Lung lysates from WT rats injected with vehicle or MMC with or without C16 were subjected to immunoblot analysis of indicated proteins (left). The relative amount of proteins normalized to β-actin (loading control) and p-eIF2a/t-eIF2a ratio is shown as mean±SEM (right). n=2-3 independent samples per condition. **d.** The amount of Rad51 and VE-Cad in the plasma of rats treated with MMC with or without C16 was quantitated by ELISA and is shown as mean±SEM. n=3 independent samples.

### Delayed ISRIB treatment reverses MMC-mediated PVOD

Next, we examined whether delayed administration of ISRIB could reverse MMC-induced vascular lesions after they had already developed. Because the elevation of RVSP (24% increase over control) and RV/LV+S ratio (52% increase over control) (**Supplementary Fig. S19a)** as well as medial hypertrophy in PAs and PVs (**Fig. 6f and Supplementary Fig. S19b**) were evident as early as 8 days (d8) after MMC administration, we tested the effect of ISRIB for either 16 days or 8 days, starting on d8 (**Fig. 8a, top**). We injected MMC in WT rats on d0, followed by administering ISRIB (0.25 mg/kg) six times between d8 and d24; the phenotype assessed on d24 (**Fig. 8a, ISRIB 16d**). Another group of WT rats was treated with ISRIB three times between d8 and d16, and their phenotypes were assessed on d24 (**Fig. 8a, ISRIB 8d**). In both 16-day and 8-day treatments, ISRIB reversed PVOD phenotypes, reducing RVSP and RV/LV+S (**Fig. 8a**). LVSP was not affected by either MMC or ISRIB treatment (**Fig. 8a**). Similarly, both 16-day and 8-day ISRIB treatments reduced vascular remodeling in PAs and PVs (**Fig. 8b, H&E**), vascular fibrosis in PAs (**Fig. 8b, blue stain, Trichrome**), and lung/body weight ratio (**Supplementary Fig. S20a)**. IF staining showed the rescue of VE-Cad and Rad51 in the vascular endothelium following delayed ISRIB treatment (**Fig. 8c**). Delayed ISRIB treatment also reduced the mRNAs of ISR-induced genes—such as *ATF3, ATF4, GADD34,* and *PKR*—in the lung tissues of MMC-treated rats (**Fig. 8d**), This result indicates effective inhibition of ISR. The reduction of Rad51 and VE-Cad mRNAs by MMC was not only reversed but also significantly induced above the basal level (**Fig. 8d**). Similarly, the delayed ISRIB treatment in MMC-treated rat lungs reversed the induction of ATF4 and PKR proteins, the increased p-PACT/t-PACT ratio, and the reduction of Rad51 and VE-Cad proteins (**Fig. 8e and Supplementary Fig. S20b**). As expected, no significant reduction in p-eIF2α was detected (**Supplementary Fig. S20c**). EdU staining revealed a robust increase in EdU-positive (proliferating) cells within the vascular smooth muscle and adventitia layers in the MMC-treated rats, which was consistent with the overgrowth of these vascular layers in these rats. But there were no proliferating cells in the CD31-positive vascular endothelium (**Supplementary Fig. S21**). When rats underwent the delayed ISRIB treatment after MMC treatment, EdU-positive cells became evident in the endothelial layer, but not in the smooth muscle or adventitia layers, indicating that ISRIB treatment promotes the proliferation of endothelial cells to restore the endothelial layer that was impaired by MMC (**Supplementary Fig. S21**). Furthermore, the elevated plasma concentrations of Rad51 and VE-Cad in MMC-treated rats were reduced to levels similar to those in vehicle-treated rats by the delayed ISRIB treatment (**Fig. 8f**). Thus, ISRIB ameliorates MMC-mediated vascular remodeling mediated in the rat model, appearing to act by inhibiting ISR and restoring VE-Cad and Rad51 in the endothelium, even eight days after the pathogenesis has been initiated.

**Fig. 8.**
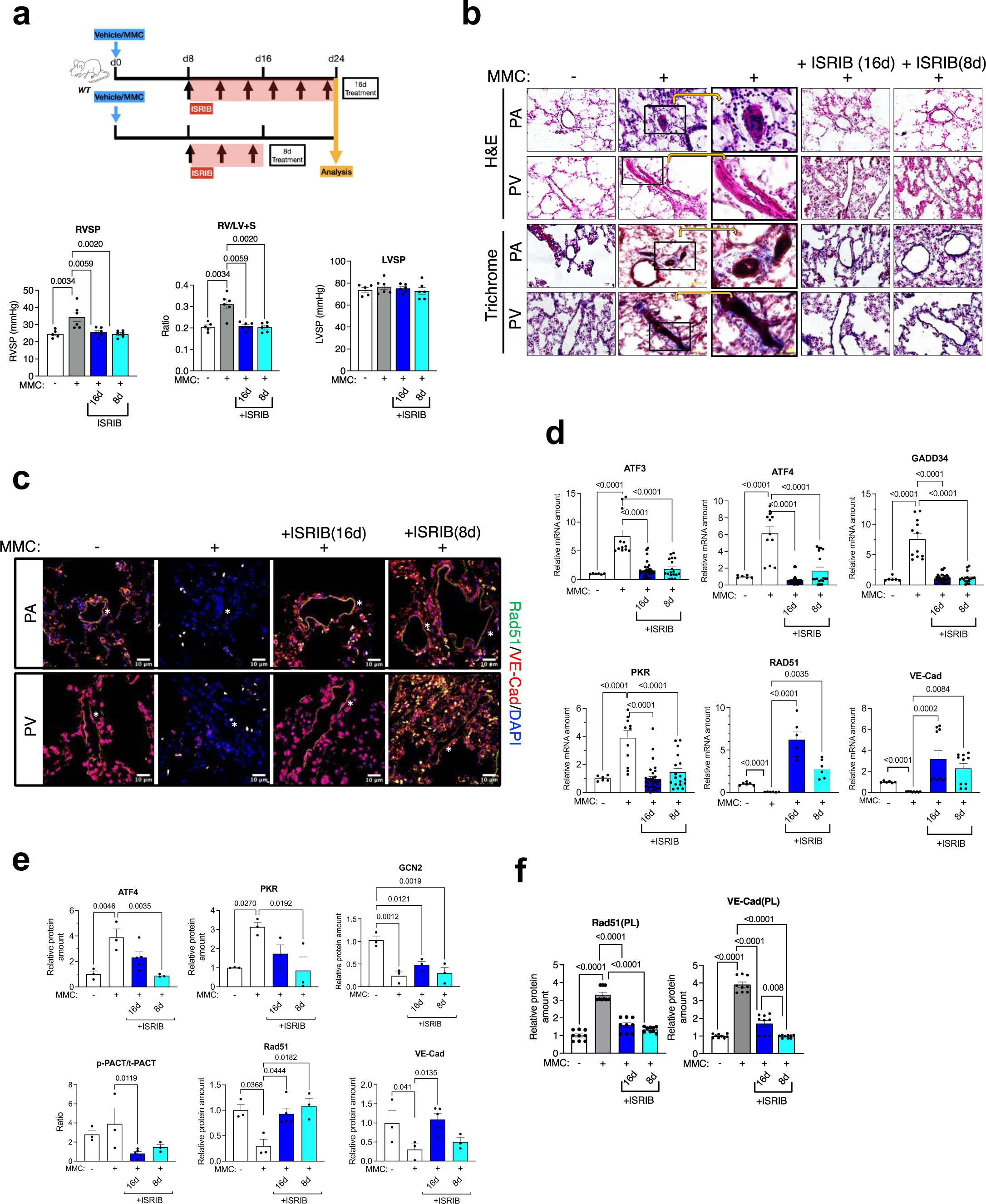
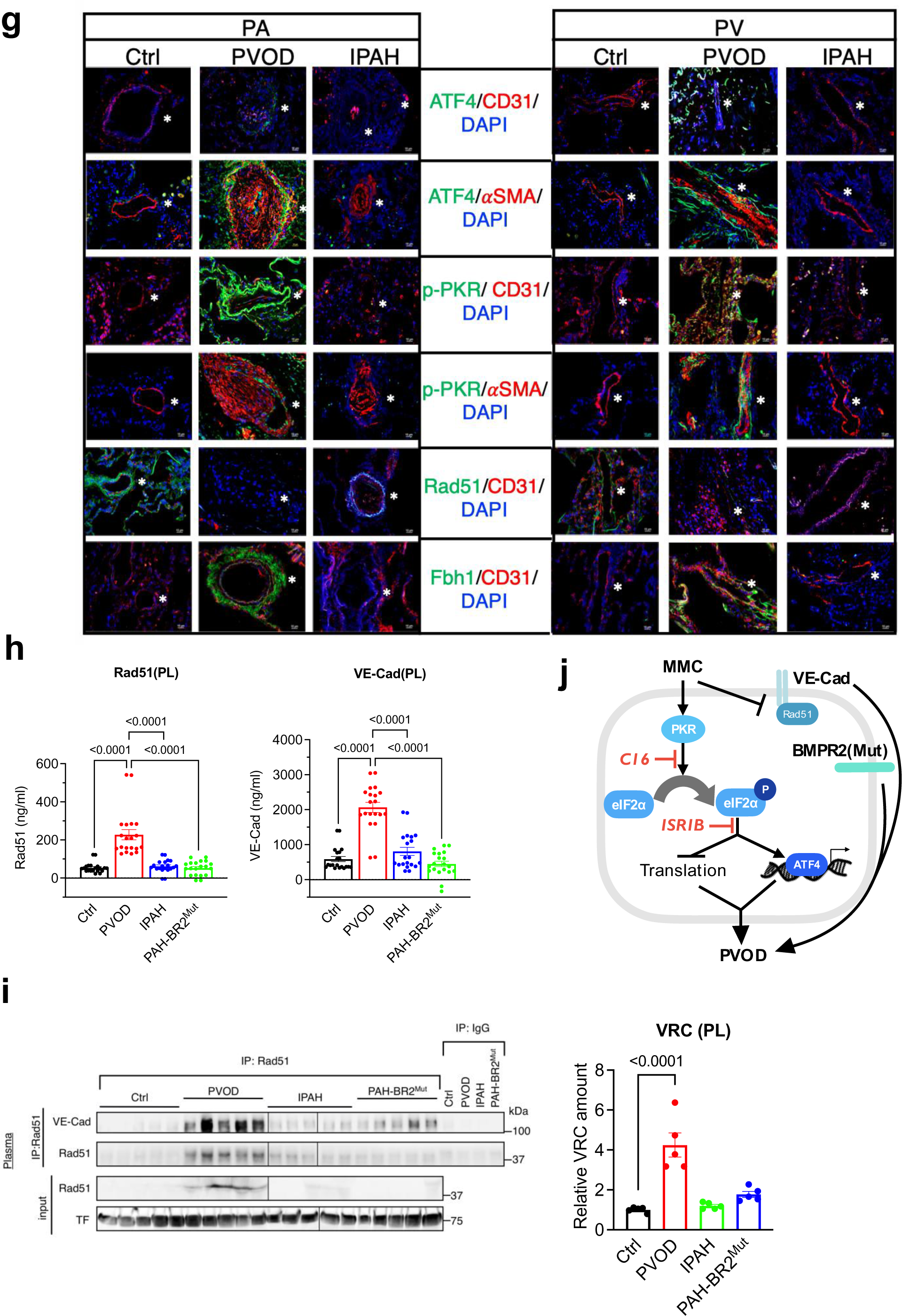
Delayed ISRIB treatment reverses PVOD phenotypes in rats. **a.** Scheme of delayed ISRIB treatment (16d and 8d treatment) (top). RVSP, RV/LV+S ratio, and LVSP in vehicle- or MMC-exposed WT rats with or without delayed ISRIB treatment (16d and 8d) are shown (bottom). n=6 independent samples. **b.** H&E and Gomori’s trichrome staining of pulmonary vasculature (PA and PV) in WT rats treated with vehicle or MMC with or without delayed ISRIB treatment (16 and 8d). The third column is a magnified image of the black rectangle area in the second column. Images are acquired at 20X magnification. **c.** IF staining of pulmonary vasculature (PA and PV) in WT rats treated with vehicle or MMC with or without delayed ISRIB treatment (16 and 8d) with anti-VE-Cad (red) antibody, anti-Rad51 (green) antibody, and DAPI (for nuclei). Scale bar=10 μm. Asterisk indicates the location of PA or PV. **d.** The amount of indicated mRNAs relative to GAPDH mRNA in the lungs of rats treated with MMC with or without delayed ISRIB treatment is analyzed by qRT-PCR and shown as mean±SEM. n= at least 6 independent samples. **e.** The amount of indicated proteins relative to GAPDH protein in the lungs of rats treated with MMC with or without delayed ISRIB treatment is analyzed by immunoblot analysis and shown as mean±SEM. n=3 independent samples per condition **f.** The amount of Rad51 and VE-Cad in the plasma of rats treated with MMC with or without delayed ISRIB treatment was quantitated by ELISA and is shown as mean±SEM. n=3 independent samples. **g.** The human lung samples from control individuals (Ctrl), PVOD, and IPAH patients were stained with anti-ATF4 (green), anti-p-PKR (green), anti-CD31(red), anti-αSMA (red), and anti-Rad51 (green), or anti-Fbh1 (green) antibodies and the images of PAs (left) and PVs (right) are taken, and merged images are shown. Cell nuclei were stained with DAPI (blue). Asterisk indicates the location of PA or PV. Scale bar=10 μm. **h.** The concentrations (ng/ml) of Rad51 and VE-Cad in the plasma of control (non-PAH) individuals, PVOD patients, IPAH patients, and PAH patients with *BMPR2* mutations (PAH-BR2^Mut^), and PVOD patients were measured in duplicate by ELISA and plotted as mean±SEM. n=10 independent samples per group. **i.** The plasma from control individuals (Ctrl) and patients with PVOD, IPAH and PAH-BR2^Mut^ were subjected to IP with an anti-Rad51 antibody, followed by immunoblot analysis of VE-Cad (for VRC) and Rad51 (left). The plasma were also subjected to immunoblot analysis of Rad51 and transferrin (TF) (loading control) (left). The relative amount of VRC is shown as mean+SEM (right). n=5 independent samples per group. **j.** A schematic diagram of MMC-induced aberrant activation of the PKR-ISR pathway and the depletion of VE-Cad and Rad51 promote the pathogenesis of PVOD, which is facilitated by the deregulation of BMPR2 signaling pathway. C16 and ISRIB are equally effective in attenuating the MMC-induced PVOD phenotypes.

Next we examined the levels of ATF4 and p-PKR in the lung samples from patients with PVOD, idiopathic PAH (IPAH), and control (non-PAH) individuals (Ctrl) by IF staining. Both ATF4 and p-PKR signals were detected in the endothelial and the smooth muscle cells of PAs and PVs from PVOD patients, but they were detected in neither vascular cells in control subjects or IPAH patients (**Fig. 8g**). This indicates that the activation of the PKR-ISR axis is specific to the pulmonary vasculature of PVOD patients. This is consistent with observations from the Sch-PH, CH-PH, and MCT-PH animal models, in which no activation of the PKR-ISR axis was detected (**Fig. 5h-k**). We also measured the plasma concentrations of Rad51 and VE-Cad in PVOD patients as well as in controls (Ctrl), IPAH patients without *BMPR2* mutations, and PAH patients with *BMPR2* mutations (PAH-BR2^Mut^) using ELISA (**Fig. 8h**) and immunoblot (**Supplementary Fig. S22**). These demonstrated ∼3-5-fold higher concentrations of Rad51 and VE-Cad in the plasma of PVOD patients (Rad51: 227.5 ng/ml and VE-Cad: 2066.0 ng/ml) compared to controls (Rad51: 54.1 ng/ml and VE-Cad: 587.9 ng/ml), IPAH patients (Rad51: 62.1 ng/ml and VE-Cad: 807.6 ng/ml), or PAH-BR2^Mut^ patients (Rad51: 52.7 ng/ml and VE-Cad: 449.4 ng/ml) (**Fig. 8h and Supplementary Fig. S22**). These results are consistent with the findings from the rat PVOD model. There was no significant increase in the plasma Rad51 or VE-Cad was observed in IPAH and PAH-BR2^Mut^ patients (**Fig. 8h and Supplementary Fig. S22**), which is consistent with the results in Sch-PH and CH-PH mice (**Fig. 5h**). The amount of VRC in the plasma of PVOD patients was 4.2-fold higher than in control individuals or patients with IPAH or PAH-BR2^Mut^ (**Fig. 8i**). The results in human not only validate the findings of the rat PVOD model in human PVOD patients, but also suggest that VRC might serve as a circulating diagnostic biomarker for PVOD. In summary, this study provides evidence that the activation of the PKR-ISR axis plays a critical role in the development of PVOD phenotypes and PKR or ISR antagonists effectively attenuate pulmonary vascular remodeling in rats (**Fig. 8j**).

## Discussion

In this study, we show that the ISR activation by PKR promotes pulmonary vascular remodeling in an animal model of PVOD. Rats carrying the BMPR2 ΔCTD mutation exhibit more severe PVOD phenotypes than their WT littermates, indicating that impaired BMPR2 and its downstream signaling pathway exacerbate vascular remodeling in the PA, PV, and capillaries. MMC treatment causes the depletion of VE-Cad and Rad51 from endothelial junctions, which then leads to the disruption of pulmonary vascular integrity. We also show that small molecule inhibitors of PKR or ISR, which restore VE-Cad and Rad51 in the endothelium and reverse PVOD phenotypes in both wild type and *BMPR2* mutant rats, and therefore, are potential therapeutics for patients with PVOD, regardless of the BMPR2 gene status.

PKR has been shown to exhibit multiple effects on the vascular endothelium, including neovascularization by promoting vascular endothelial growth factor (VEGF) production^49^ and TNF-α-induced inflammation through modulation of endothelial adhesion molecules^50^. The inhibition of PKR by C16 blocks both angiogenesis and tumor cell growth in vivo^51^. Thus, it is plausible that the inhibition of MMC-mediated vascular remodeling by C16 may be a combined effect of attenuating PKR-dependent ISR activation and inhibiting the pro-angiogenic and pro-inflammatory activities of PKR.

We found evidence of the colocalization and coregulation between endothelial junctional protein VE-Cad and the DNA repair enzyme Rad51. Endothelial cell-cell junctions contain both TJs and AJs, each with their respective set of adhesion molecules^22, 26^. AJs are formed by the cadherin superfamily of proteins and VE-Cad plays a central role in their stabilization and control of permeability through the association with catenins and the actin cytoskeleton^52^. Therefore, the amount, stability, localization, and adhesions of VE-Cad are regulated by multiple mechanisms^22, 26^. It has been reported that VE-Cad undergoes proteolytic cleavage and the extracellular domain of VE-Cad (∼90 kDa) is shed into the extracellular space experiencing apoptosis ^30, 53, 54^. Importantly, the molecular weight of VE-Cad secreted upon MMC stimulation is ∼130kDa, which is indistinguishable from VE-Cad located at EC junctions. Furthermore, when EV from MMC-treated EC were added to the media of recipient EC impaired by MMC, it restores the endothelial barrier, suggesting that VRC-containing EV could potentially be used therapeutically to repair endothelial barrier damage. The mechanisms of EV biosynthesis and their release, and the identity and biological activity of EV cargo other than the VRC are still unclear and necessitate further investigation. The increased presence of VRC in the plasma of PVOD patients, but not in IPAH patients or control individuals, highlights the potential of this complex as a diagnostic biomarker for PVOD, a finding that carries significant clinical implications. In addition to its barrier function, it is reported that ZO-1 (TJP-1) plays a role in modulating the potency of the BMP-Smad signaling pathway in pluripotent stem cells^55^. Thus, it is plausible that MMC-mediated depletion of ZO-1 in the vascular endothelium could alter the BMP-Smad signaling activity, which then contributes to vascular remodeling.

The functions of nuclear Rad51 in DNA double strand break repair, DNA replication, and meiosis are well established^56^; however, the role of cytoplasmic Rad51 is less known, despite a significant amount of Rad51 being localized in the cytoplasm^57, 58^. Rad51 interaction site was mapped at the cytoplasmic domain of VE-Cad (aa 715-730) that overlaps with one of LC3 motifs, which are shared among 18 members of the cadherin superfamily ^27^. Furthermore, LC3 motifs are 100% conserved among VE-Cad orthologs in vertebrate species ^27^, thus we speculate that the Rad51–VE-Cad association in EC and its role in the preservation of junctional integrity are evolutionarily conserved. It also raises the possibility that Rad51 interacts with other cadherin proteins in other cell types and protects cadherins from autophagosome-mediated degradation. Activation of autophagy has been implicated in the pathogenesis of IPAH based on the result of the increased amount of LC3B, one of the LC3 isoforms, in the lung of IPAH patients^59^ as well as in the pulmonary vascular endothelium of Sugen5416/hypoxia PAH model rats^60^. Chloroquine, which blocks the binding of autophagosomes to lysosome^61^, partially attenuates the pathogenesis in monocrotaline-induced rat model of PAH^62^. On the contrary, a decreased level of LC3B has been reported in the intima of PAs in a rat model of chronic thromboembolic PH ^63^. It is important for future studies to determine whether the induction of autophagy is detectable in the pulmonary vascular endothelium of PVOD patients.

While mutations in the *BMPR2* gene are accountable for >70% of PAH^7^, a small number of PVOD patients with *BMPR2* mutation have been reported^9, 64, 65^, and *EIF2AK4* mutations remain a leading genetic cause of PVOD^1, 3^. However, a recent study reported that 8.5% of PAH patients with no known gene mutations carry *EIF2AK4* mutations, which would identify *EIF2AK4* as the second most mutated gene in PAH^7^. The BMPR2 ΔCTD mutant, which mimics one of the BMPR2 gene mutations found in PAH^66^, exacerbates MMC-induced PVOD phenotypes. Previously, it was found that the depletion of BMPR2 is sufficient to diminish Rad51 in PMVECs, suggesting that the carriers of the BMPR2 ΔCTD mutant possess a reduced basal amount of Rad51 in the endothelium^18^. A small amount of VRC has been detected in the plasma of PAH patients with *BMPR2* mutations, but not in IPAH patients or control individuals. Furthermore, the administration of an ISR inhibitor reverses PVOD phenotypes in MMC-treated *BMPR2* mutant rats. These findings indicate possible crosstalk between the deregulated BMPR2 signaling pathway and an increase in the secretion of VRC, and/or the activation of the ISR pathway. Genetic ablation of PKR has been found to alleviate PH in both monocrotaline and Sugen5416/hypoxia models of PAH in rats^67^. Furthermore, PAH is a well-recognized complication of human immunodeficiency virus (HIV) infection^68^. Increased expression of human endogenous retrovirus-K (HERV-K) has been observed in the lungs of PAH patients^69^. Since PKR is activated upon viral infection by binding to double-stranded RNA^70^, it is plausible that the activation of the PKR-ISR pathway might be linked to the pathogenesis of PAH associated with HIV infection. Consequently, attenuation of this pathway by C16 or ISRIB might prove effective in treating other types of PAH in addition to PVOD.

The ISR pathway is essential for organisms to adapt to a changing environment and maintain homeostasis^19, 33^. However, maladaptive activation of the ISR is linked to cognitive and neurogenerative disorders, cancer, and inflammation^19, 33^. ISR activity has been reported to increase with age and is linked to the aging process^71^. Increased levels of PKR and p-eIF2α have been detected in various organs of aged animals, including the kidney, liver, brain, lung, and heart^72^. The administration of ISRIB has been found to restore memory impairments after traumatic brain injury^73, 74^, alleviate cognitive impairments in Down syndrome^75^ and mitigate the effects of aging^76^ and noise-induced hearing loss^77^, and inhibit cancer growth^78^. Our study uncovers a new therapeutic application for ISRIB: the amelioration of pulmonary vascular disease. The effect of ISRIB on improving cognitive impairment due to injury and aging has been attributed to its direct effect on neuronal structure and function, such as dendritic spine formation, in the hippocampus^73, 74, 76^. ISRIB exhibits a half-life of approximately 8 h in mice^73^ and can penetrate the blood-brain barrier without overt toxicity in mice or rats^79^. The binding cavity of ISRIB in eIF2B is evolutionarily conserved from yeast to human^42^. This makes ISRIB effective in preclinical studies across a variety of species, as well as in human patients. Our findings indicate that ISRIB can restore vascular endothelial homeostasis by ISRIB, which raise the possibility that neurocognitive improvements associated with ISRIB might be partially mediated by enhanced cerebrovascular function and/or by the crosstalk between the nervous system and the vascular system in the brain.

Interestingly, the loss of PERK and GCN2 in both humans and mice result in distinct phenotypes despite both kinases performing the identical function of eIF2α phosphorylation and inhibition of global protein synthesis ^33^. PVOD is mediated by biallelic LOF mutations in *EIF2AK4* (encoding GCN2)^3^, whereas Wolcott-Rallison syndrome (WRS, OMIM 226980)—a disease characterized by growth retardation, early-onset diabetes, and epiphyseal dysplasia—is associated with biallelic LOF mutations in *EIF2AK3* (encoding PERK)^80, 81^. Homozygous *EIF2AK4* (GCN2) knockout mice are viable, fertile, and exhibit no phenotypic abnormalities under normal conditions, except failing to adapt to amino acid deprivation^82^. Homozygous *EIF2AK3* (PERK) knockout in mouse recapitulates WRS phenotypes and exhibits growth retardation, neonatal diabetes, and skeletal malformation^83, 84^. PERK is localized in the endoplasmic reticulum (ER) and activated upon unfolded proteins in the ER, which subsequently activating ISR, while GCN2 is localized to the cytosol and activated upon amino acid deprivation^33^. Given GCN2 and PKR are activated in a spatially and temporally specific manner under distinct cellular stress, we speculate that they transmit distinct signals to mediate different biological outcomes^19^. It is intriguing to speculate that despite the genetic inactivation of PERK, eIF2α kinase(s) other than PERK, could mediate maladaptive ISR activation and contribute to the pathogenesis of WRS like PVOD.

## Materials and Methods

Reagents, kits, antibodies, PCR primers, siRNAs, instruments, and software used in the study are listed in the Supplementary Information.

### Animal care and use

All animal experiments were conducted in accordance with the guidelines of the Institutional Animal Care and Use Committee (IACUC) of University of California, San Francisco. The protocol number for the relevant animals and procedures approved by IACUC is AN185765-02E: Title: Role of Growth Factor Signaling in Vascular Physiology” (Approval Date: December 09, 2022).

### Generation of Bmpr2 mutant rats

*Bmpr2* mutant Sprague-Dawley rats were constructed using TALEN technology at Taconic Biosciences with the left arm: 5’-GCACAGTGTGCTGAGGAGA-3’ and the right arm: 5’-TGATATGGGAGAGAA-3’. Two lines of rats with monoallelic deletion of 2 bp (Δ2) in exon 13 *(Bmpr2^Δ2/+^*) or deletion of 26 bp (Δ26) in exon 13 *(Bmpr2^Δ26/+^*) resulting in frameshift mutation of *Bmpr2* E503fs and Q495fs, respectively, were developed in parallel. Both strains and both sexes were studied and compared to confirm the effects of *Bmpr2* mutations in the development of PVOD on the analyzed parameters. WT (+/+) littermates were used as controls.

### MMC-mediated PVOD rat model and administration of C16 and ISRIB

Animals were housed in the vivarium of the cardiovascular research building at UCSF (San Francisco, USA). Both male and female Sprague Dawley rats (8-9 weeks old) were subjected to following protocols to examine the effect of MMC and/or small molecule inhibitors of ISR (ISRIB; Sigma-Aldrich, #19785) or PKR (C16; Sigma-Aldrich, SML0843). MMC was made by dissolving 2 mg MMC in 1 ml saline and was delivered to rats at 3 mg/kg dosage through i.p. injections. Saline was used as vehicle solution for MMC treatment. ISRIB solution was made by dissolving 5 mg ISRIB in 1 ml of dimethyl sulfoxide (DMSO) (Sigma, D2650), followed by dilution to 1 mg/ml and was delivered to rats at 0.25 mg/kg dosage through i.p. injections. The vehicle solution consisted of 1 ml DMSO and 4 ml saline. C16 solution was made by dissolving 10 mg C16 in 1 ml DMSO, followed by dilution to final concentration of 100 μg/ml and was delivered to rats at 33.5 μg/kg dosage through i.p. injections. The vehicle solution consisted of 100 μl DMSO and 10 ml saline.

#### Protocol #1 MMC treatment

Rats were randomly divided into MMC (3 mg/kg) or saline (vehicle)-exposed groups. MMC or saline was administered once by intraperitoneal injection (i.p.) on day 0 (d0). Rats were euthanized at 24 days after the MMC/vehicle injection (d24). Hemodynamic measurements were made, RV hypertrophy was assessed, and tissue samples were collected.

#### Protocol #2 Simultaneous MMC and ISRIB treatment for 24 day

ISRIB (0.25 mg/kg) or vehicle (DMSO) was given by i.p. at 30 min before MMC (3 mg/kg) treatment on d0. Rats were given ISRIB or vehicle by i.p. 3 times a week until d24 (total nine injections) and euthanized on d24.

Hemodynamic measurements were made, right ventricle hypertrophy was assessed, and tissue samples were collected.

#### Protocol #3 Simultaneous MMC and C16 treatment for 24 day

C16 (33.5 μg/kg) or vehicle (DMSO) was given i.p. 30 minutes before MMC (3 mg/kg) treatment on d0, followed by C16 (33.5 μg/kg) or vehicle treatment on d8 and d16 (total 3 injections) and euthanized on d24. Hemodynamic measurements were made, right ventricle hypertrophy was assessed, and tissue samples were collected.

#### Protocol #4 Delayed ISRIB treatment for 8 or 16 day

Rats are given MMC (3 mg/kg) or vehicle (saline) by i.p. on d0. For 16-day treatment, ISRIB (0.25 mg/kg) or vehicle (DMSO) was given 3 times a week between d8 and d24 (total 6 injections) and euthanized on d24. For 8-day treatment, rats were given ISRIB (0.25 mg/kg) or vehicle 3 times a week between d8 and d16 (total 3 injections) and euthanized on d24. Hemodynamic measurements were made, right ventricular hypertrophy was assessed, and tissue samples were collected.

### Isolation of pulmonary artery smooth muscle cells (PASMC)

Primary rat PASMC were isolated according to the standard protocol^85^. The pulmonary artery of anaesthetized WT or Mut rats was removed under the sterile condition and transferred to a culture dish with cold (4°C) DMEM. After removal of the fat tissue around the artery, the artery was longitudinally cut and placed in another cell culture dish containing DMEM. Then, we used a pair of ophthalmic curved tweezers to scrape the intima softly to get rid of ECs. The artery was cut into small tissue blocks. The ophthalmic curved tweezers were used to separate the media from the artery by pressing and pushing the artery with its blunt back side. After half of the media was removed, the same method was used to obtain another half. The media was cut into approximately 1-mm squares and transferred into cell culture plates. The plates were placed in a cell culture chamber for about 4 h to let the small tissue blocks adhere to the plates. DMEM containing 20% FBS was added, and the tissue blocks were incubated in the cell culture chamber without disturbance for the first 5 days. The PASMC began to grow out from the edge of the tissue blocks at about 8 days and became relatively confluent by approximately 16 days. PASMC were identified through morphology (‘hill-and-valley’ pattern) and immunofluorescence detection of α smooth muscle-actin (αSMA). The purity of the PASMC was tested through multiple fluorescent staining with DAPI and an anti-αSMA antibody.

### Monocrotaline (MCT)-induced rat PH model

MCT (#C2401, Sigma-Aldrich) was administered to WT or Mut rats (8-9 weeks old) following the protocol ^86, 87^. In brief, MCT was dissolved in 0.5 N HCl to 200 mg/kg (adjusted pH at 7.4 with 0.5 N NaOH) and then diluted with sterile water to 60 mg/ml. MCT (60 mg/kg) or vehicle (PBS) was subcutaneously injected in the ventral thorax of rats. The animals were maintained at 12-h light–dark cycle at 18–20 °C and 40–50% for 32 days (food and water were provided ad libitum and the animals were checked once per day), followed by the assessment of the phenotypes.

### Schistosomia- and chronic hypoxia (CH)-induced mouse PH models

The levels of proteins were quantified in the whole lung lysates and the plasma samples of preclinical PH models: *Schistosoma mansoni (S. mansoni)*-induced PH model and CH-induced PH model, which are known to recapitulate key features of PAH. For Schistosoma model: *S. mansoni* eggs were obtained from infected mice from the Biomedical Research Institute (Rockville, MD). Mice (C57BL/6) were sensitized intraperitoneally (240 eggs/g) and 14 days later intravenously (175 eggs/g) with *S. mansoni* eggs as described previously^37^. Seven days later, the lung and the plasma samples were harvested for the analysis. Mice that were not exposed to *S. mansoni* eggs were used as control. For CH model: Mice (C57BL/6) were placed in the hypoxia chamber connected to the oxygen feedback sensor ProOx 360 (BioSpherix, Parish, NY) that senses and maintains the oxygen concentration equivalent to 10% FiO2 inside the chamber by infusing nitrogen (N_2_) gas for 21-days. At the conclusion of hypoxia exposure, the lung and plasma samples were collected to perform protein assessment. Mice in room air were used as control.

### Hemodynamic measurement and tissue histology

The animals were anesthetized with an intraperitoneal injection of a ketamine/xylazine cocktail solution (1 ml ketamine (100 mg/ml) + 100 µl xylazine (20 mg/ml); inject 300 µl per 250 g body weight). A tracheal cannula was then inserted, and the animals were ventilated with room air using VentElite rodent ventilator (Harvard Apparatus) set to maintain respiration at 90 breaths/min and tidal volume at 8 ml/kg body weight. The abdominal and thoracic cavity of the rat was opened carefully to avoid any blood loss, and a 2F pressure-volume catheter (SPR-838, Millar AD Instruments, Houston, TX) was used for RVSP measurements. RVSP was measured while a consistently stabilized pressure wave was shown after the transducer was plugged into RV apex. At the end of the experiments, the hearts and lungs were perfused with phosphate-buffered saline (PBS) for blood removal. Fulton index, or the weight ratio of the right ventricle divided by the sum of left ventricle and septum [RV/(LV + S)], was measured and calculated to determine the extent of right ventricular hypertrophy. Lung, liver, and heart tissues were fixed in 10% formalin for 24 h and then further processed for paraffin sectioning. The paraffinized lung tissue sections were used for hematoxylin–eosin (H&E)^88^ and Gomori’s trichrome staining^89^ according to standard protocols. The images were acquired by Olympus BX51 microscope (Olympus), Ts2 microscopes (Nikon), and Leica SPE confocal microscope. Total areas of fibrotic lesion within each section were quantified using a threshold intensity program from ImageJ.

### Assessment of vascular remodeling

To assess pulmonary artery and vein muscularization, rat lung tissue sections (10 µm in thickness) were subjected to conventional H&E staining. The external and internal diameter of a minimum of 50 transversally cut vessels in tissue block ranging from 25 to 80 µm were measured by determining the distance between the lamina elastica externa and lumen in two perpendicular directions as described previously^90^. The vessels were subdivided based on their diameter (microvessels: <50 µm and medium sized vessels: 50-80 µm) and the assessment of muscularization was performed using ImageJ in a blinded fashion by a single researcher to reduce operator variability, who was not aware of the group allocation of the samples being analyzed. The absolute value of the medial thickness was converted to the relative value by setting the medial thickness of vehicle-treated wild type (WT) rats as 1. Additionally, we also assessed the muscularization of pulmonary arteries and veins by the degree of αSMA immunofluorescence staining. The IF signal intensity was quantitated by ImageJ, and the result is presented as a relative signal intensity by setting the value of vehicle-treated WT rats as 1. Images were acquired using an Olympus BX51 microscope, Ts2 microscopes (Nikon), and Leica SPE confocal microscope.

### Casting of pulmonary vasculature with Microfil

The procedure for the casting of pulmonary vessels with Microfil was described previously^87^. Briefly, heparin (1000 UI/kg), as an anticoagulant, was injected intravenously, 10 min before anesthesia. The rats were anesthetized using a ketamine/xylazine cocktail and after removal of the anterior chest wall, a microperfuser tube was inserted and kept in the right ventricle via a needle (25 G) for perfusion with PBS which drained from the left atrium. The lungs were perfused to clear all blood as evidenced by the lung tissue turning white. During the perfusion of pulmonary arteries (PAs), freshly dissolved Microfil polymer mixture (MV compound: MV diluent: MV agent = 5:5:1) was instilled via 25 G needle inserted into the PA from the incision through the RV wall by manual injection. The Microfil mixture was gently infused into the PA under a dissecting microscope until it reached to the terminal branches of PAs and then stopped within 2–3 sec. The lungs were then kept at room temperature for approximately 90 min or overnight at 4℃ while covered with a wet paper towel to avoid desiccation of the lungs. At the end of the experiment, the dissected lungs and hearts were rinsed in PBS for 10–15 min at room temperature. They were then dehydrated in a series of ethanol solutions (50%, 70%, 80%, 95%, and 100%; 2 h each). After dehydration, the lungs were put into a methyl salicylate (Sigma-Aldrich) solution. When the lungs became translucent and the Microfil was clearly visible, they were photographed. The number of branches and junctions in the distal pulmonary vascular networks of five lobes were counted by ImageJ.

### Perfusion of pulmonary vasculature with Evans Blue dye (EB) and the analysis of vascular permeability in vivo

Perfusion of pulmonary vasculature was performed as described previously^91^. Briefly, 2% (weight/volume) EB solution was prepared by mixing EB (#E2129, Sigma) in 0.9% NaCl, followed by filtering a 0.22μm filter. Rats were anesthetized with isoflurane vaporizer set to 5% in 100% oxygen for 5 min. After the incision was made in the midline of the animal and exposed the diaphragm, using forceps, grabbed the sternum and pulled towards the head of the animal, pressed the heart against the diaphragm until easily visible. Inserted the needle through the diaphragm into the LV and injected 2% EB solution (6μl/g) with a 3 ml syringe capped with a 25 G needle. Waiting for ∼5 min to allow the EB to circulate and confirming the successful administration of the EB by observing the snout, paws, and tail turning blue, the animals were euthanized and proceeded to tissue harvest. The harvested tissue was fixed in 4% PFA for O/N at 4°C. After washing in 1X PBS with shaking at R.T. for 3 x 30 min, the tissue was dehydrated with methanol/H_2_O series: 20%, 40%, 60%, 80%, 100%; 1 h each. After further washing with 100% methanol for 1 h, the tissue was incubated for 3 h with shaking in 66% Dichloromethane (DCM, #270997, Sigma) / 33% methanol at R.T., followed by being incubated in 100% DCM 2 x 15 min with shaking. The tissue was then incubated in 100% Dibenzyl ether (DBE, #108014, Sigma) for about until the tissue became translucent and photographed. The permeability of pulmonary vasculature of five lobes was examined by quantitating the intensity of EB stain by ImageJ.

### Human plasma samples

All plasma samples were obtained following informed consent from the UK National Cohort Study of idiopathic and heritable Pulmonary Arterial Hypertension (clinicaltrials.gov NCT01907295; UK REC Ref. 13/EE/0203) following institutional guidelines and following informed consent. Healthy adult controls were recruited for comparison studies. The subsequent whole-blood sample collection was performed under written informed consent of the participants or their parents for use in gene identification studies (UK Research Ethics Committee: 08/H0802/32) and process as described before^92^. Blood samples for research purposes were collected in 6 ml EDTA tubes using standard venepuncture protocols. Plasma samples were subjected to centrifugation and stored at −80 °C before use. Patients diagnosed with IPAH (without *BMPR2* mutation), PAH with BMPR2 mutation, or PVOD, relatives of index cases and unrelated healthy controls were recruited at nine UK centers and followed up by a median of 7.9 years. All cases were diagnosed between March 1994 and November 2016, and diagnostic classification was made according to international guidelines^93^. Clinical, functional, and hemodynamic characteristics at the time of PAH or PVOD diagnosis were prospectively entered into the database. The date of diagnosis corresponded to that of confirmatory right heart catheterization.

### Human lung samples

Human lung tissues from PH patients and control individuals were collected by the Pulmonary Hypertension Breakthrough Initiative (PHBI) in the US. The organization of the PHBI, under the direction of the Cardiovascular Medical Research and Education Fund and supported by R24HL123767, is outlined at http://www.ipahresearch.org/. Lung tissue from patients with PVOD and IPAH was obtained at the time of explant for lung transplantation. Control lung tissue was obtained from unsuccessful organ donors enrolled in the PHBI as control subjects. Clinical data for the PH patients have been published^94^. The present study was approved by both the Colorado Multiple Institutional Review Board (IRB) and the UCSF IRB. The collection of lung tissue was approved by the IRB at all lung transplant sites. For lung tissue processing, the PHBI follows a standardized tissue-processing protocol, as detailed previously.^94^

### Enzyme-linked immunosorbent assay (ELISA)

The amount of VE-Cad and Rad51 protein in the conditioned media (CM) of PMVECs or human and rat plasma samples containing EDTA was quantitated by using ELISA kit (DCADV0, R&D systems for VE-Cad and custom made by MyBiosource for Rad51) according to the manufacturer’s protocol. The relative concentrations of total proteins in the CM or plasma were quantitated by Bradford protein estimation method using NanoDrop 2000c (ThermoScientific) and the sample volume was adjusted by the total protein amount. The CM or plasma samples were diluted 1:50 and 1:20 in PBS, respectively prior to applying to precoated ELISA plates. Plates were washed with PBS-T three times followed by incubation with respective secondary antibody conjugated to streptavidin-horseradish peroxide (HRP) at room temperature (RT) for 2 h. The ELISA was developed in a dark room at RT with a colorimetric substrate comprising tetramethylbenzidine and stabilized hydrogen peroxide mixed in equal volume. The absorbance was measured at 450 nm using SpectraMax M (Molecular Devices LLC).

### Cell culture and MMC treatment

Human primary pulmonary microvascular endothelial cells (PMVECs) were purchased from ScienCell Research Laboratories (#3000, Carlsbad, CA). The culture dish was coated with 0.2% gelatin and the cells were cultured in endothelial growth medium (EGM)-2 complete media (#CC-3162, Lonza Clonetics, Fisher scientific) with EGM-2 SingleQuots supplement kit (CC-4176, Lonza Clonetics, Fisher scientific). All experiments were performed using low-passage cells (passage 3-8). Saline (vehicle) or MMC (150 μM) was added to the culture media (EGM-2 media with supplemented with 0.02% FBS) of PMVECs at 60-70% confluency and treated for 2-14 h as indicated.

### Transfection of siRNAs and plasmids

Small inhibitory RNA (siRNA) for non-targeting control, Rad51, Fbh1, Nedd4-1, Nedd4-2, and Smurf1 (Dharmacon Thermo Scientific) was transfected into PMVECs at 60%-70% confluency. Details of siRNAs are found in Supplementary Table 1. 9.3 μl Lipofectamine RNAiMAX 2000 (#13778-150, Invitrogen) was mixed with 750 μl Opti-Minimum Essential Medium (Opti-MEM purchased from Gibco, Thermo Scientific, Waltham, MA) followed by 5 min incubation at RT. siRNA was mixed with 750 μl of Opti-MEM with final concentration of 100 nM. Furthermore, both the Lipofectamine mixture and siRNA mixture were pooled and incubated for 20 min at RT. Following the incubation, the pooled mixture was added to endothelial cells and put at 37 °C. A 10ml of EGM-2 complete media was added to endothelial cells after 5 h. Finally, the siRNA transfected endothelial cells were harvested after 48 h. Harvested samples were subjected to either immunoblot analysis or quantitative reverse transcriptase PCR (qRT-PCR) to determine the knockdown of genes. For human Fbh1 or Rad51 expression plasmid transfection, PMVECs were seeded in p6-well plates at a concentration of 270,000 cells/well; 24 h later, cells were transiently transfected with 1.5 μg of plasmid using FuGENE 6 transfection reagent (#E2692, Promega), according to the manufacturer’s instructions, for 2.5 h. The cells were washed twice with 1xPhosphate buffered saline (PBS) and allowed to rest at 37°C for 40 h in complete EGM-2 media. Transfected cells were then treated with MMC (150 μM, 0.02% FBS) or vehicle (saline) for the indicated times. As control, original pcDNA3.1(+) was used. Fbh1 and Rad51 expression plasmids were obtained from GeneCopoeia (#EX-E2953-Lv105) and Addgene (CMV-hRad51, #125570), respectively (Supplementary Information).

### Alkaline comet (DNA damage) assay

To assess the extent of DNA damage in human PMVECs, we performed alkaline comet assay using the alkaline comet assay kit (Trevigen Inc., Gaithersburg, MD). The kit manufacturer’s protocol was followed while performing the assay. Briefly, siRAD51 and siFbh1 transfected and vehicle (saline) or MMC (150 μM) for 14 h treated human pulmonary microvascular endothelial cells were washed with ice-cold 1xPBS. Further, cells were de-attached, centrifuged and suspended at 2 × 10^5^ cells/mL in ice-cold 1x PBS. A mixture of 50 μL of suspended cells and 500 μL LMAgarose was spread onto Cometslide^TM^ and sited at 4 °C for 10 min in dark. Next, Cometslide^TM^ was immersed in lysis solution for 60 min at RT. After carefully removal of the lysis solution, slides were immersed in an alkaline unwinding solution for 30 min at RT in the dark. Following the unwinding, the electrophoresis was performed in an alkaline electrophoresis solution for 35 min at 4 °C. The electrophoresis run condition was kept constant as 1V/cm and 300 mA. Slides were carefully immersed in dH_2_O twice after removal of an alkaline electrophoresis buffer and then kept in 70% ethanol for 5 min in dark at r.t. Next, slides were air dried at 37 °C for 15 min and stained with SYBR® Gold for 30 min in the dark. The slides were viewed under a laser scanning confocal microscope (Leica SPE, Buffalo Grove, IL) for nuclei with a comet-like elongated tail.

### Nuclear and cytoplasmic fractionation

PMVECs were washed with 1xPBS twice, scrape off and pelleted by centrifuging at 4,500 g for 5 min. Cells were then swelled by adding 5 volume of lysis buffer (10 mM HEPES, pH 7.9, with 1.5 mM MgCl_2_, 10 mM KCl, 1mM DTT and protease inhibitor, sigma, P8340) and homogenized. After centrifugation at 10,000 g for 15 min, the supernatant was collected as a cytoplasmic fraction. The crude nuclei pellet was resuspended in 2/3 volume extraction buffer (20 mM HEPES, pH 7.9, with 1.5 mM MgCl_2_, 0.42 M NaCl, 0.2 mM EDTA, 25% (v/v) Glycerol, 1mM DTT and protease inhibitors (Sigma P8340) and homogenized with a tissue homogenizer. After centrifuging at 20,000 g for 5 min, the supernatant was collected as a nuclear fraction.

### Preparation of conditioned media (CM)

PMVECs (passage 3-5) cultured in a 10 cm plate with 10 ml particle-free media for 48 h at 70–80% confluency at the time of harvest (3-4×10^6^ cells) were treated with vehicle (saline) or MMC for 14 h. Cells were then washed with 1xPBS three times. The culture media were collected and subjected to centrifugation at 2000×*g* for 20 min at 4°C to remove large cell debris and the supernatant was then filtered through 0.22 mm-diameter filter and syringe to remove small cell debris and large vesicles and used as CM.

### Isolation of Extracellular vesicles and particles (EV)

EV was isolated using a total exosome isolation kit (Invitrogen) ^95, 96^. In brief, 2 ml conditioned media (CM) of PMVECs or 250 μl rat plasma samples were centrifuged at 2000×*g* for 20 min at 4°C to remove cell debris. Plasma samples were further centrifugated at 10,000x*g* for 20 min and the supernatant was transferred to a new tube on ice and mixed with 0.5 volume of 1xPBS. For CM, EV isolation reagent (0.5 volume; ∼1 ml) was added and incubated overnight at 4°C, followed by centrifugation at 10,000X*g* for 1 h. For plasma samples, EV isolation reagent (0.2 volume; ∼50 μl) was added and incubated for 10 min at RT, followed by centrifugation at 10,000X*g* for 5 min. The pellets from CM or plasma samples were resuspended in 120 μl of ice-cold 1xPBS and sequentially centrifuged at 500x*g* for 10 min, at 3,000x*g* for 20 min, and then at 12,000x*g* for 20 min. The pelleted EV were resuspended in 120 μl of ice-cold 1xPBS. The protein concentration of the EV was measured by Bradford assay using NanoDrop 2000c (Thermo Scientific). Proteins derived from EV were mixed with 4X Laemmli buffer (750 mM Tris-HCl pH6.8, 5% SDS, 40% glycerol and 80 mM DTT), heated to 95 °C for 5 min, and subjected to immunoblotting. The purity of EV was determined by the relative levels of CD63 by the immunoblot.

### Nanoparticle tracking analysis (NTA)

The number and size distribution of nanoparticles in the CM or rat plasma samples were measured through NTA, which was performed by a NanoSight NS300 instrument (Malvern Panalytical) equipped with a 488 nm laser and the NTA 3.4 Build 3.4.4 analytic software at the Parnassus Flow Cytometry facility (CoLab) at UCSF. Briefly, samples were diluted in 0.1µm filtered PBS to achieve a concentration of approximately 1–9 × 10^8^ particles/ml and injected into the NanoSight sample chamber using a 1 ml syringe and syringe pump. Three 30 sec videos were captured at camera level 8 and frame rate of 32 per second; and analyzed. The modal diameter and the per cell/animal ratio of nanoparticles were analyzed statistically.

### Immunoblot analysis

We used boiling lysis buffer (10 mM Tris-HCl, 1% SDS, 0.2 mM phenylmethylsulfonyl fluoride) to prepare the whole protein cell lysate. Protease and phosphatase inhibitors (1:100; Sigma-Aldrich, St Louis, MO), were added to the boiling lysis buffer. The boiling lysis buffer was directly added to cells and boiled for 10 min followed by 35 min centrifugation at 12000 rpm. The rat tissue lysates were prepared in the lysis buffer (1% Triton X-100, 150mM NaCl, 50mM Tris-Cl at pH 7.5, 1mM EDTA). The supernatants were collected, and total protein concentration was measured by NanoDrop 2000c (Thermo Scientific). Proteins samples were denatured in SDS-sample buffer for 5 min at 95°C and loaded onto Mini-Protean TGX^TM^ gels (BioRad Laboratories) in equal amounts and subjected to electrophoresis. Nitrocellulose membrane (Genesee Scientific) was used to blot the gels, which were blocked with 5% non-fat milk or 3% BSA in 1x Tris buffered saline with 0.1% tween-20 (1x TBST) for 1 h at RT. The membranes were incubated at 4 °C overnight with a primary antibody.

Chemiluminescence signals were detected using SuperSignal™ West Dura extended duration substrate (ThermoFisher) and imaged using an Odyssey Dlx Imaging System (LI-COR). Antibodies used for immunoblots are found in Supplementary Table 1. The quantity of each protein was normalized to the amount of a loading control protein. Subsequently, its relative quantity was calculated by setting the amount of the protein in the control (vehicle-treated) sample as 1.

### Immunoprecipitation assay

Total cell lysates or CMs of PMVECs treated with saline or MMC (150 nM) for 14 h, rat tissue samples, plasma samples, or EV isolated from CM or plasma samples were lysed in IP buffer (1% Triton X-100, 150mM NaCl, 50mM Tris-Cl at pH 7.5, 1mM EDTA) supplemented with protease inhibitors (1:100 dilution) and phosphatase inhibitor (1:100 dilution). Lysates were nutated for 30 min at 4 °C, followed by centrifugation at 12,000 g for 10 min and supernatant were collected. One tenth of the lysate was saved as an input sample for immunoblot. The lysate was incubated with an anti-Rad51 and anti-IgG (negative control) nutating overnight at 4°C followed by the addition of dynabeads^TM^ Protein A/G and rocking for 4 h at 4 °C. The magnetic beads were precipitated and rinsed thrice with IP buffer for 5 min at 4 °C. The washed elute was boiled at 95°C for 8 min in a sample loading buffer and subjected to immunoblot along with input. For the input samples: initially, the quantity of the indicated protein was normalized to the amount of a loading control protein, such as beta-actin or GAPDH. Subsequently, its relative quantity was calculated by setting the amount of the protein in the vehicle-treated sample as 1. For the IP samples, the amount of the indicated protein in MMC-treated sample was presented with the protein amount in the control (vehicle-treated) sample set as 1.

### RNA-seq analysis

Total RNA was isolated from the endothelial cells using RNeasy Mini Kit (#74104, Qiagen). The quality of RNAs was evaluated with a 2100 Bioanalyzer Instrument (Agilent Technologies). RNA samples with RNA integrity number (RIN) > 8.0 were shipped to Beijing Genome Institute for library preparation and sequencing (Illumina HiSeq 2500). Around 70 million reads were obtained for each pooled sample. To read raw sequence data and set quality checks we used FastQC v0.72. FASTQ files were trimmed with Trimmomatic v0.38.1 to remove low quality reads and any adapter. The reads were mapped with HISAT2 v2.1.0 to the human genome (hg38 / GRCh38). After mapping, all BAM files were used as input for HTSeq-count v0.91 to calculate transcript coverage. DESeq2 (v2.11.40) was used to find deferentially expressed transcripts between samples for each sequencing depth. Deferentially expressed genes (DEG) were kept if the adjusted p value was equal or below 0.05 and if a log2 fold-change of 1 or greater was observed. Integrative Genomics Viewer (2.4.14; Broad Institute) was used for data analysis.

### Chromatin immunoprecipitation (ChIP)

PMVECs treated with vehicle (saline) or MMC (150 nM) for 4 h were crosslinked with 1% formaldehyde for 15 min at RT, followed by quenching with 1M Glycine. Cells were washed with 1xPBS and lysed with lysis buffer (50 mM Tris-Cl pH 8.1, 10 mM EDTA, 1% SDS and protease inhibitor). Genomic DNA was sheared to an average length of 200-500 bp by sonication, followed by clearing lysates by centrifugation at 12,000g for 10 min at 4°C. Incubated the supernatant with protein A/G dynabeads at 4°C for 1 h, diluted the pre-cleared sample to a 1:10 ratio with dilution buffer (20 mM Tris-Cl pH 8.1, 150 mM NaCl, 2 mM EDTA, 1% Triton X-100 and protease inhibitor) and 1/10 volume was kept as input before incubation with non-specific IgG (control), or an anti-ATF4 antibody overnight at 4°C followed by incubation with protein A/G dynabeads. Next, the dynabeads were washed with a buffer I (20 mM Tris-Cl pH 8.1, 150 mM NaCl, 2 mM EDTA, 1% Triton X-100, 0.1% SDS), buffer II (20 mM Tris-Cl pH 8.1, 500 mM NaCl, 2 mM EDTA, 1% Triton X-100, 0.1% SDS), and buffer III (10mM Tris-Cl pH8.1, 250mM LiCl, 1mM EDTA, 1%NP-40, 1% Deoxycholate) at 4°C. The dynabeads were further washed twice with cold TE (10 mM Tris-Cl pH 8.1, 1 mM EDTA) and incubated in 250 μl elution buffer (200 mM NaHCO_3_, 1% SDS) at RT for 15 min twice. The eluates were mixed with 1/25 volume 5M NaCl and incubated at 65 °C for 4 h. 1/50 volume of 0.5 M EDTA, 1/25 volume of Tris-Cl pH 6.5, proteinase K (final 100 μg/ml) were added and incubated at 45°C for 1 h. Precipitated DNA fragments were purified with QIAquick PCR Purification Kit, followed by qRT-PCR analysis. The PCR primer sequences are found in **Supplementary Table 1**.

### Reverse Transcriptase-quantitative Polymerase Chain Reaction (qRT-PCR)

Total RNA was isolated from PMVECs and Rat lung tissue and subjected to cDNA preparation by the reverse transcription reaction using an iScript cDNA Synthesis Kit (#17088890, Bio-Rad). qPCR analysis was performed in triplicate using iQ SYBR Green Supermix (#1708882, Bio-Rad). The relative expression values were determined by normalization to *GAPDH* transcript levels and calculated using the ΔΔCT method. qRT-PCR primer sequences are found in Supplementary Table 1.

### Proximity ligation assay (PLA)

PMVECs were grown on coverslips, followed by the treatment with vehicle (saline) or MMC (150 nM) for 14 h. Cells were washed thrice with 1xPBS, fixed with 4% paraformaldehyde, and permeabilized by 0.15% TBST for 30 min. Cells were then blocked using 5% goat serum in TBST for 1 h, and subjected to PLA with an anti-Rad51 and anti-VE-Cadherin (see Supplementary Table 1) using the Duolink in situ red starter kit (mouse/rabbit) (Sigma-Aldrich) following the manufacturer’s protocol. Images were captured by a confocal microscope (Leica SPE).

### Treatment of PMVECs with ISRIB or C16

ISRIB (200 nM) or C16 (500 nM) was added to the culture media of PMVECs at 15 min before the vehicle (saline) or MMC (150nM) treatment for 14 h. For another experiment, 2 h after the vehicle (saline) or MMC (150nM) treatment, ISRIB (200 nM) or C16 (500 nM) was added to the culture media of PMVECs and incubated for 14 h. After the indicated time the cells were subjected to the Transwell vascular permeability assay.

### Transwell vascular permeability (TVP) assay

PMVECs were seeded in the upper ‘apical’ chamber of 0.4-mm transwell inserts (Corning, MA) that were coated with collagen IV. After culture for 5 days post-confluence, PMVECs were treated with MMC (150nM), or vehicle (saline) for the indicated times. One hour before the sampling time-point, stock concentrations of molecular tracers were added to the upper chamber to bring the final concentration to 0.5 mg/ml. Samples were collected from the upper (luminal) and lower (abluminal) chambers for fluorometry analysis. The molecular tracers (Sigma, MO), such as sodium fluorescein (MW ∼0.4k Da), Evans Blue dye (MW ∼1k Da), and fluorescein isothiocyanate (FITC)-conjugated dextran (MW ∼40k Da). The contribution of the molecular tracers in the bottom chamber was used as a proxy to assess the barrier functionality of PMVECs. The concentration of each tracer was determined using a standard curve for fluorescence intensity (FCI) or permeability coefficient (Ps). The permeability coefficient was calculated as follows: Ps=[A]/t×1/A×V/[L], where [A] is the abluminal concentration, t is time (in seconds), A is the area of the membrane (in cm^2^), V is the volume of the abluminal chamber, and [L] is the luminal concentration. FCI or Ps for each tracer is represented as the mean±SEM from three independent experiments.

### Trans-endothelial electrical resistance (TEER) assay

PMVECs barrier function was determined as described previously^3,4^, by measuring the cell–cell adhesive resistance to electric current using an EVOM Volt/Ohm meter (World Precision Instruments). Briefly, PMVECs were seeded onto transwell pre-coated with collagen IV. After culturing for 5 days post-confluency, the transwells were subjected to total electrical resistance measurement using electrods and recorded TEER are normalized to the baseline. The time when MMC (150nM) or vehicle (saline) was added was set to zero. TEER changes were recorded and submitted for statistical analyses at specified time-points (n=3 independent experiments).

### Immunofluorescence (IF)

PMVECs were grown on coverslips till at 60-70% confluence and transfected with siRNAs or Rad51 expression plasmid followed by vehicle (saline) and MMC (150nM), permeabilized, blocked, and subjected to primary antibody incubation. Anesthetized rats were flushed with 1X PBS and perfused with 4% (w/v) paraformaldehyde and organs like lung, heart, and liver were post-fixed in 4% paraformaldehyde, transferred to 1X PBS after 24 h, and subjected to paraffin embedding. Additionally, the right bronchus of flushed lungs was sutured, and the left lung inflated with 1% low melt agarose for fixation and paraffin embedding, and the right lung split for snap freezing for protein and RNA studies. Immunofluorescence was performed using antibodies listed in the Supplementary information. For SMA and EC staining, sections were subjected to deparaffinization, antigen retrieval, and permeabilization followed by blocking and primary antibody incubation overnight at 4°C. A solution of Alexa Fluor secondary antibodies (Invitrogen) was applied for 2 hours at RT. IF images were acquired using a confocal microscope (Leica SPE) or Eclipse Ts2 Inverted LED phase contrast microscope (Nikon) and analyzed using ImageJ. Antibodies used are found in the Supplementary Information.

### In vivo 5-ethynyl-2′-deoxyuridine (EdU) labeling and detection

To determine cellular proliferation, EdU (50 mg/kg body weight) was administered intraperitoneally to rat by i.p. injection 24 h before sacrifice following the previously published protocol^97^. After the EdU incorporation, lung was isolated, fixed in 4% PFA, and paraffin embedded sections were mounted on slides. Lung sections were permeabilized with 0.5% Triton X-100 in PBS and EdU labeling was detected by using Click-iT™ EdU Cell Proliferation Kit (#C10337; Invitrogen) according to the manufacturer’s protocol. The lung sections were co-stained with an endothelial cell marker CD31 (sc-376764; Santa Cruz Biotechnology) and signals were detected by Nikon Ts2 fluorescence microscope.

### Annexin V staining

To determine apoptosis, rat lung FFPE sections was stained with Annexin V (#11060-1-AP; Proteintech), co-stained with endothelial cell marker CD31 (sc-376764; Santa Cruz Biotechnology), and immunofluorescence signal were detected by Nikon Ts2 fluorescence microscope according to the manufacturer’s protocol.

### Glutathione-S-transferase (GST)-fusion protein pull-down assay

Recombinant proteins, which contain the entire cytoplasmic region of mouse VE-Cad (Cyto; aa 621-789), Cyto region except deleted in aa 780-784 (Δ5), proximal region (Prox; aa 621-689), middle region (Mid; aa 661-730), and distal region (Dist; aa 715-789), all fused to GST^7^, were used for mapping the Rad51 interaction site. GST-pull-down assay was performed as described previously^8^. Briefly, *Escherichia coli* BL21 (GE Healthcare) was transformed with each GST-fusion protein construct and the expression of GST-fusion protein was induced by 0.1 mM isopropyl-β-D-thiogalactoside (IPTG) at 37°C for 4 h. After lysing bacteria, GST-fusion protein was isolated by affinity chromatography on glutathione-Sepharose 4B beads (Cytiva). PMVECs lysates (50 μg) were incubated with 30 μl of a 50% slurry of GST-fusion protein conjugated agarose beads for 2 h at 4°C under constant agitation. After washing the beads for five times with buffer (10 mM Tris pH 8, 1 mM EDTA pH 8, 150 mM NaCl, 5 mM DTT, 1% Triton X-100, 1X protease inhibitor cocktail), proteins associated with the GST-fusion protein were eluted by boiling for 5 min in SDS sample buffer, subjected to SDS–PAGE, and analyzed by immunoblot with an anti-Rad51 antibody (Supplementary Information).

### In vivo puromycin incorporation assay

Wild type rats were injected once with either a vehicle (saline) or MMC (3 mg/kg). Twenty-four days after the injection, the rats were perfused with 100 µg/ml puromycin in 1XPBS via intracardiac injection. Ten minutes following the perfusion, the lungs were harvested, and total lung lysates were subjected to SDS-PAGE. Immunoblot was conducted using an anti-puromycin antibody.

### Statistical analysis

All numerical data are presented as mean ± standard error of the mean (SEM). Statistical analysis was performed using Microsoft Excel and GraphPad Prism 10 (La Jolla, CA). Datasets with two groups were subjected Student’s *t*-test, unpaired, equal variance whereas comparison among three or more than three groups was done by ANOVA followed by Tukey’s post hoc corrections. Analysis of variance was applied to experiments with multiple parameters, one- or two-way as appropriate. And, where required, significance was analyzed using a post hoc Tukey test and indicated as P-values.

### Data availability

Sequencing data presented in this study have been deposited in the NCBI Gene Expression Omnibus (GEO) database (https://www.ncbi.nlm.nih.gov/geo/) under the accession code: GSE229404.

## Authors Contributions

The authors confirm contribution to the paper as follows. *Study conception and design*: Amit Prabhakar and Akiko Hata. *Execution of experiments*: Amit Prabhakar, Rahul Kumar, Meetu Wadhwa, Carlos Lizama Valenzuela, Jingkun Zhang, Prajakta Ghatpande, Ziwen Zhao, Bhushan Kharbikar. *Data analysis*: Amit Prabhakar, Rahul Kumar, Meetu Wadhwa, Carlos Lizama Valenzuela, and Akiko Hata. *Interpretation of results*: Amit Prabhakar, Rahul Kumar, Meetu Wadhwa, Carlos Lizama Valenzuela, Stefan Gräf, Carmen Treacy, Nicholas Morrell, Brian Graham, Giorgio Lagna, and Akiko Hata. *Primary draft manuscript preparation*: Amit Prabhakar and Akiko Hata. All authors reviewed the results and approved the final version of the manuscript.

## Acknowledgements

We thank Dr. Rubin M. Tuder, Ms. Aneta Gandjeva (Univ. of Colorado Anschutz Medical Campus), and the Pulmonary Hypertension Breakthrough Initiative (PHBI) for providing human lung samples. We also thank the UK National Cohort of Idiopathic and Heritable PAH (UKNCPAH) for providing human plasma samples. We also thank Drs. Xuan Jiang (UCSF and San Yat-Sen Univ.), Sanna Vattulainen-Collanus (Blueprint Genetics), Tejal A. Desai (UCSF and Brown Univ.), Ms. Ananyaa Arvind (UCB), Simren Gupta (UCSB), Yuhang Yao (Tshinghua Univ.) and members of the Atabai lab for technical advice and support. We thank Dr. Klaus Ebnet (Univ. of Münster) for a kind gift of GST-VE-Cad fusion constructs and Drs. Kamran Atabai, Biao Wang and Guo Huang (UCSF) for sharing reagents. Grant funding was provided by National Heart, Lung, and Blood Institute (NHLBI; R01HL132058, R01HL153915, and R01HL164581) and Tobacco-Related Disease Research Program (28IR-0047) to A.H.; American Heart Association 19CDA34730030, Cardiovascular Medical Research Fund (CMREF) and United Therapeutics Jenesis Innovative Research Award to R.K.; by NHLBI (R01HL135872 and P01HL152961) to B.B.G. N.W.M. is supported by a Programme Grant from the British Heart Foundation (RG/19/3/34265) and Personal Chair Award (CH/09/001/25945). The UKNCPAH is supported by the British Heart Foundation (SP/12/12/29836), the BHF Cambridge Centre of Cardiovascular Research Excellence, and the UK Medical Research Council (MR/K020919/1).

---

**Supplementary Fig. S1.**
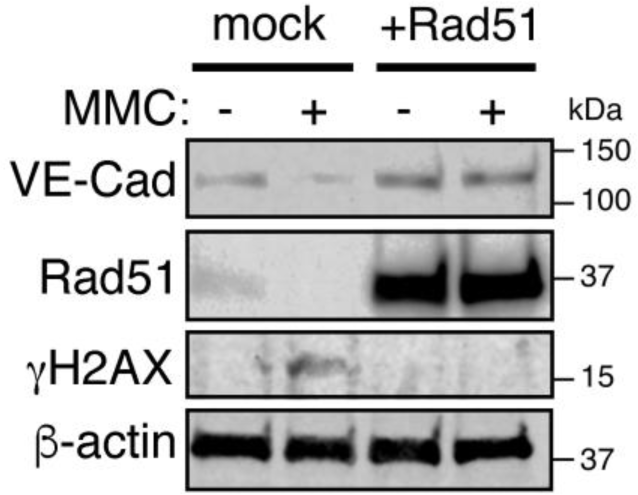
Overexpression of Rad51 prevents MMC-mediated DNA damage in PMVECs. PMVECs (1×10^6^ cells) were transfected with empty vector (mock) or Rad51 expression plasmid (+Rad51), followed by vehicle or MMC treatment for 14 h. Total cell lysates were subjected to immunoblot of VE-Cad, Rad51, γH2AX, and β-actin (control).

**Supplementary Fig. S2.**
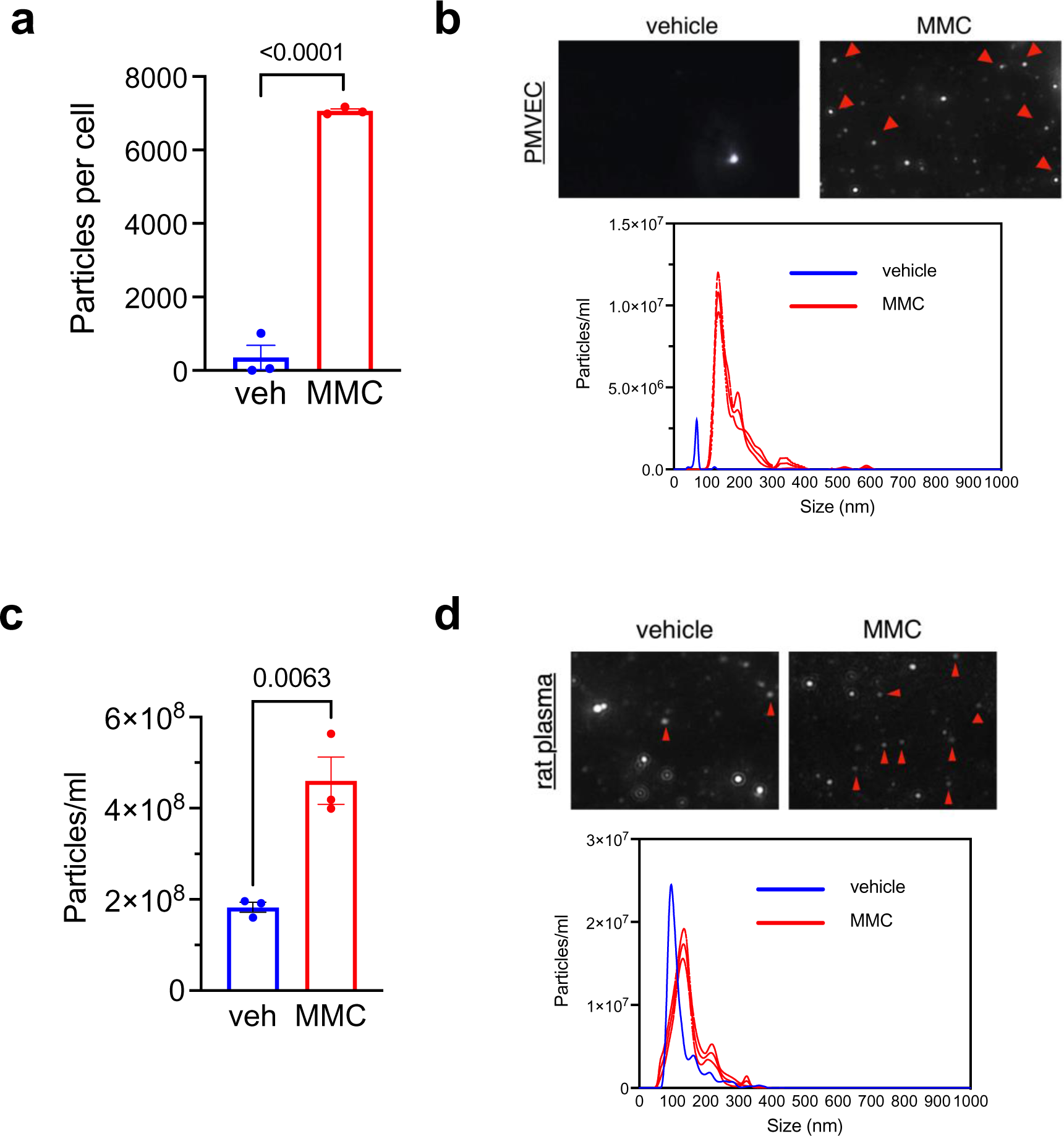
Nanoparticle tracking analysis (NTA) of EV released into the conditioned media and the rat plasma. **a.** The number of EV particles (100-200 nm in diameter) released from vehicle- or MMC-treated PMVECs (100,000 cells) was quantitated by NanoSight and shown as a particle number per cell (mean±SEM). n=3. **b.** Representative image of EV particles released from PMVECs after stimulation with vehicle or MMC for 14 h (top) and size distribution (bottom) was examined by the NTA (NanoSight). Blue=EV from vehicle-treated cells. Red=EV from MMC-treated cells. n=3. Red arrowheads indicate EV. **c.** The number of EV particles (140-200 nm in diameter) was quantitated by NanoSight and shown as a particle number per plasma (ml) (mean±SEM). n=3 independent samples. **d** Representative image of EV released into the plasma of rats administered with vehicle or MMC (top) and size distribution (bottom) was examined on day 16 by the NTA (NanoSight). n=3 independent samples. Red arrowheads indicate EV.

**Supplementary Fig. S3.**
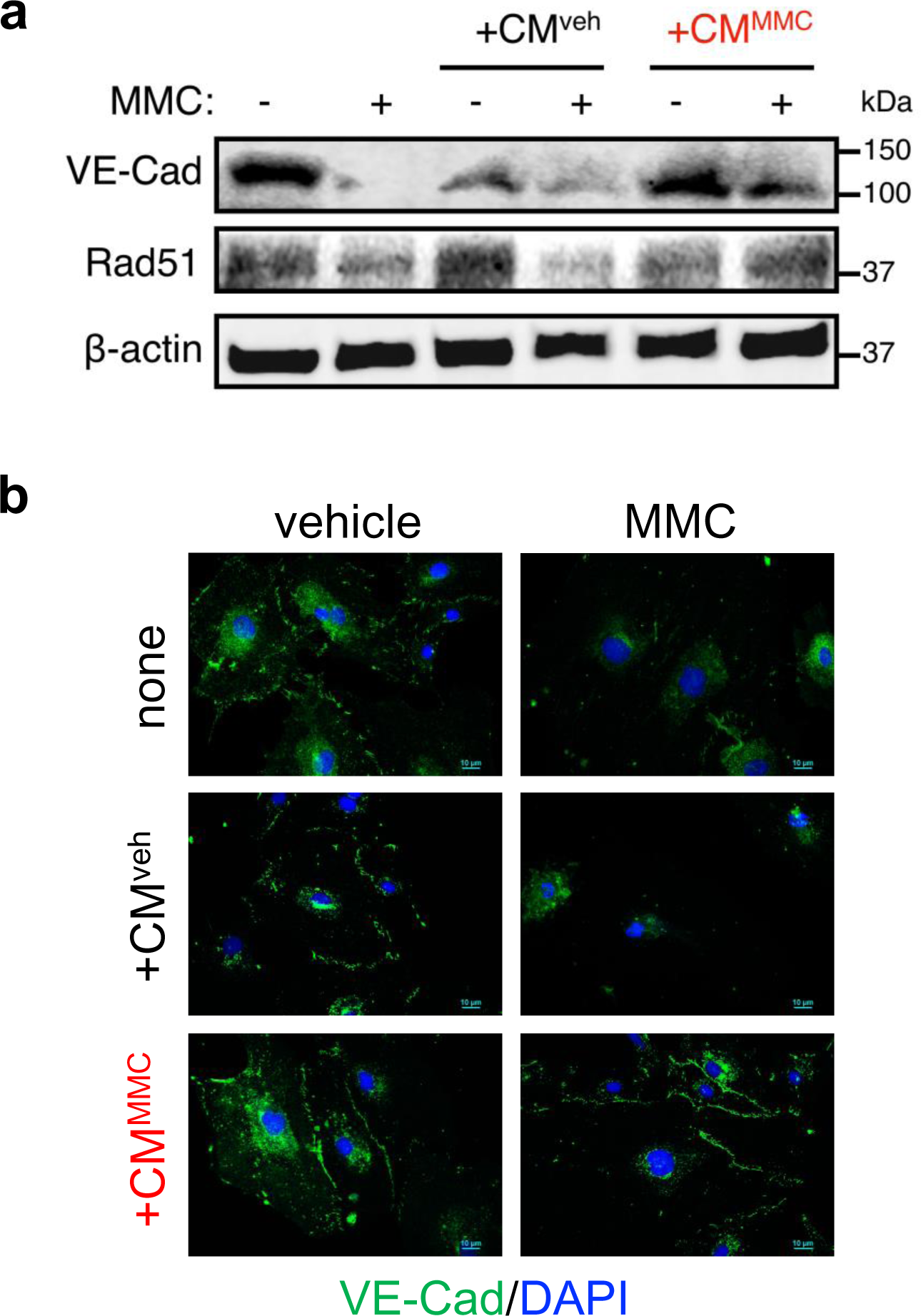
EV from the MMC-stimulated PMVECs can transfer their cargo-VE-Cad and Rad51-to the MMC-damaged recipient PMVECs. **a.** The levels of intracellular VE-Cad, Rad51, and β-actin (loading control) protein in the MMC-treated PMVECs (recipient cells; 1×10^6^ cells) after the incubation with the CM from PMVECs (1×10^6^ cells) treated with vehicle (CM^veh^) or MMC (CM^MMC^) for 14 h were examined by immunoblot analysis. **b.** IF staining of PMVECs treated with CM^veh^ or CM^MMC^ (5×10^3^ cells) with an anti-VE-Cad antibody (green). Cell nuclei were stained with DAPI (blue). Scale bar=10 μm.

**Supplementary Fig. S4.**
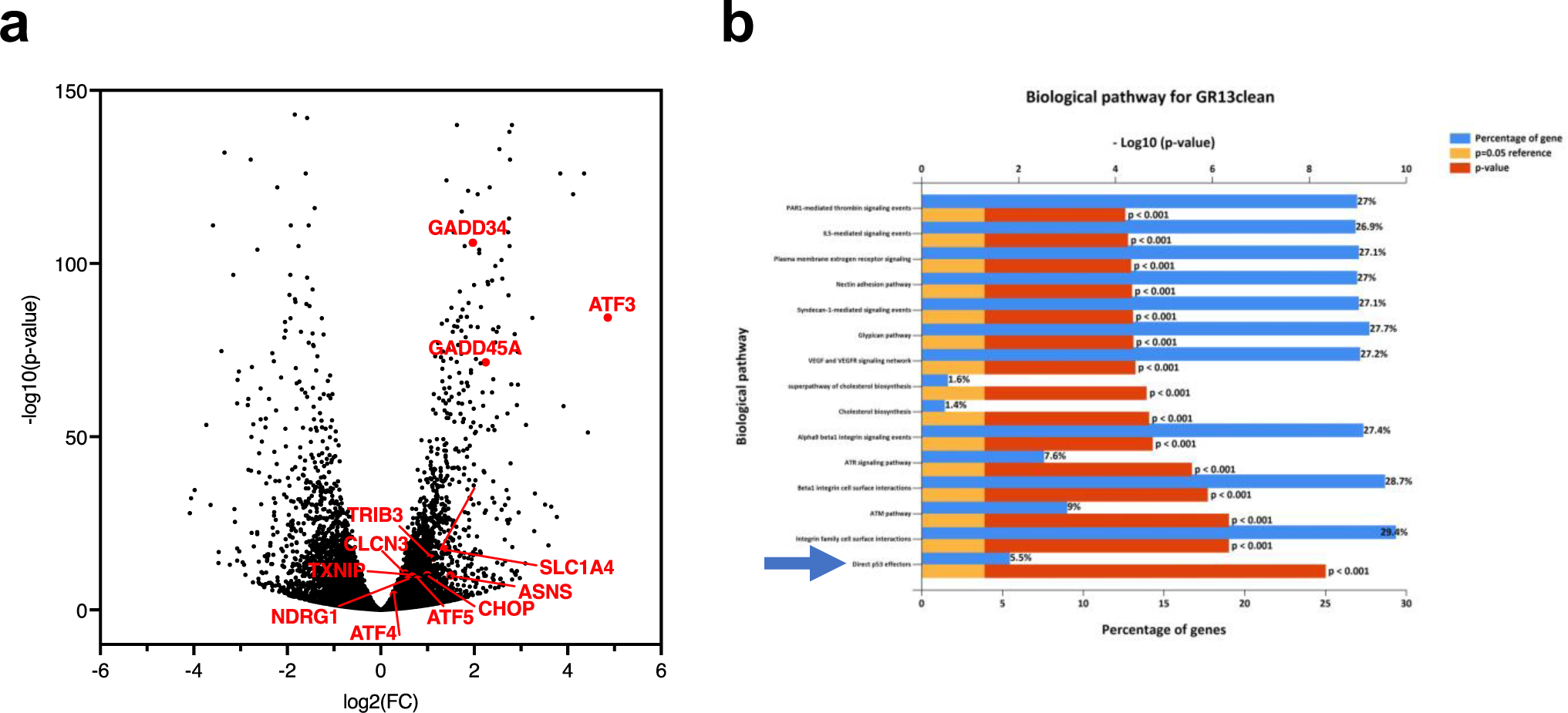
Transcriptional activation of ATF4 target genes upon MMC treatment in PMVECs. **a.** RNA-seq data (MMC 0 h vs 4 h) is plotted in volcano map. X-axis and Y-axis indicate p-value (-Log_10_) and fold change (FC) (Log_2_). Annotated red dots indicate known ATF4 targets. n=3 independent RNA samples per condition **b.** The result of the pathway analysis of differentially expressed genes in PMVECs treated with MMC for 0 h vs 14 h are shown. p53 pathway is indicated with blue arrow.

**Supplementary Fig. S5.**
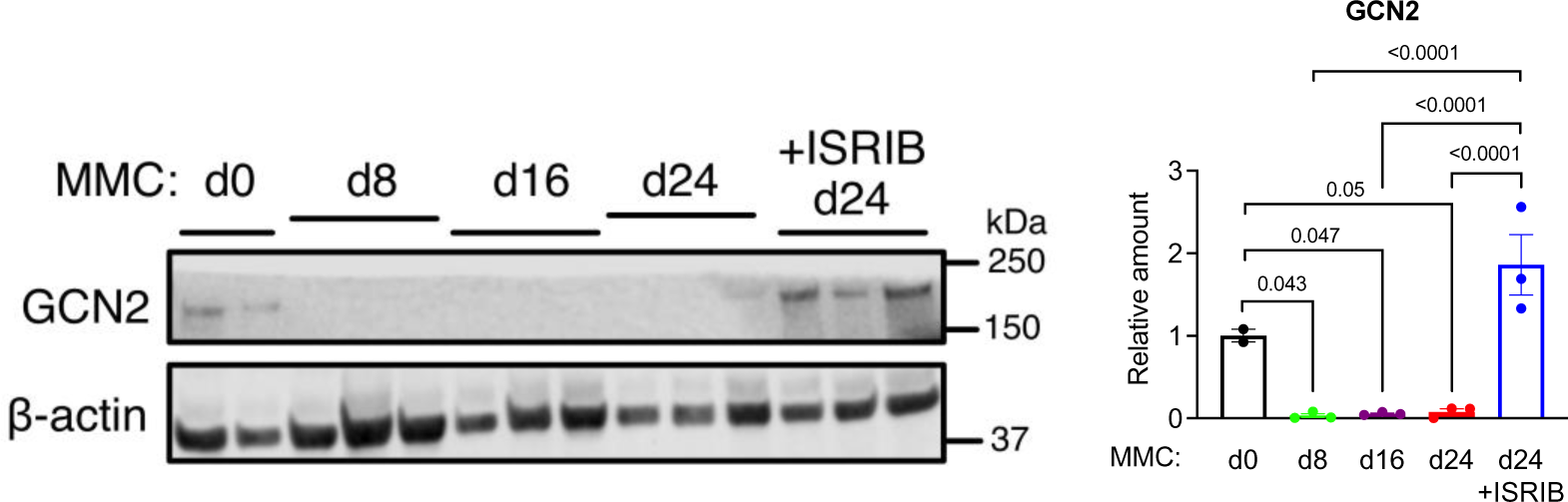
A time-course changes in GCN2 protein amount after MMC treatment. Lung lysates of WT rats 0, 8, 16, or 24 days after MMC administration with or without ISRIB (24 days) were subjected to immunoblot of GCN2 and β-actin (loading control) (left). The amount of GCN2 relative to β-actin is shown as mean±SEM (right). n=2 or 3 independent samples per condition.

**Supplementary Fig. S6.**
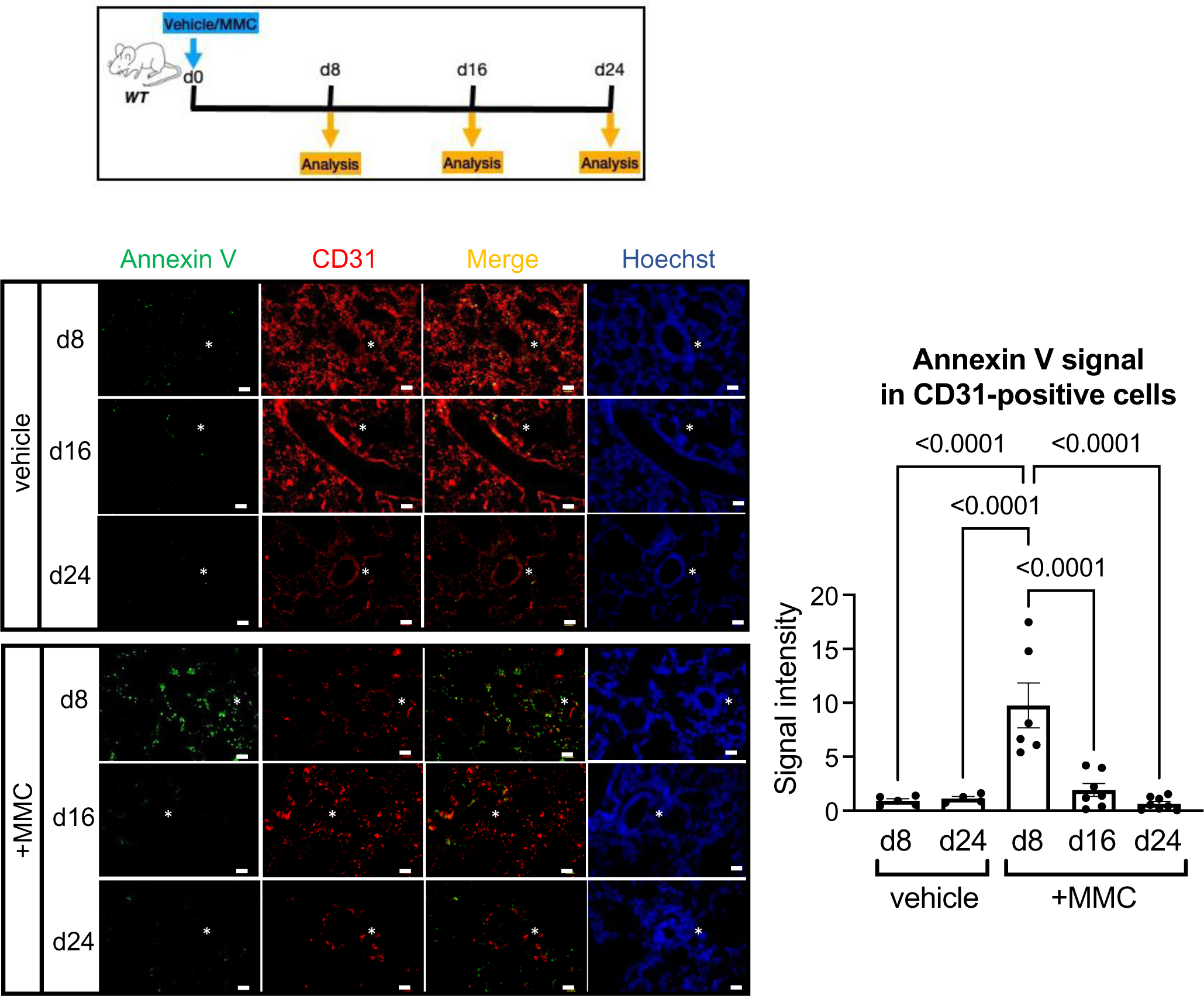
Apoptotic cells are detected in the pulmonary vascular endothelium in WT rats 8 days after MMC treatment. Schematic description of the experimental procedure (top left). Lung samples of WT rats treated with vehicle or MMC were harvested on day 8, 16, and 24. IF staining of the apoptosis marker Annexin V (green) and CD31(red) for vascular endothelial cells (bottom left). Hoechst (blue) is showing nuclei. The images of Annexin V and CD31 stain were merged and shown in the “Merge” row (left). The intensity of Annexin V signal in the CD31-positive endothelial layer was quantitated by ImageJ and shown as mean±SEM (right). n=4∼7 independent samples per condition. White asterisks indicate pulmonary arteries. Scale bar= 10 μm

**Supplementary Fig. S7.**
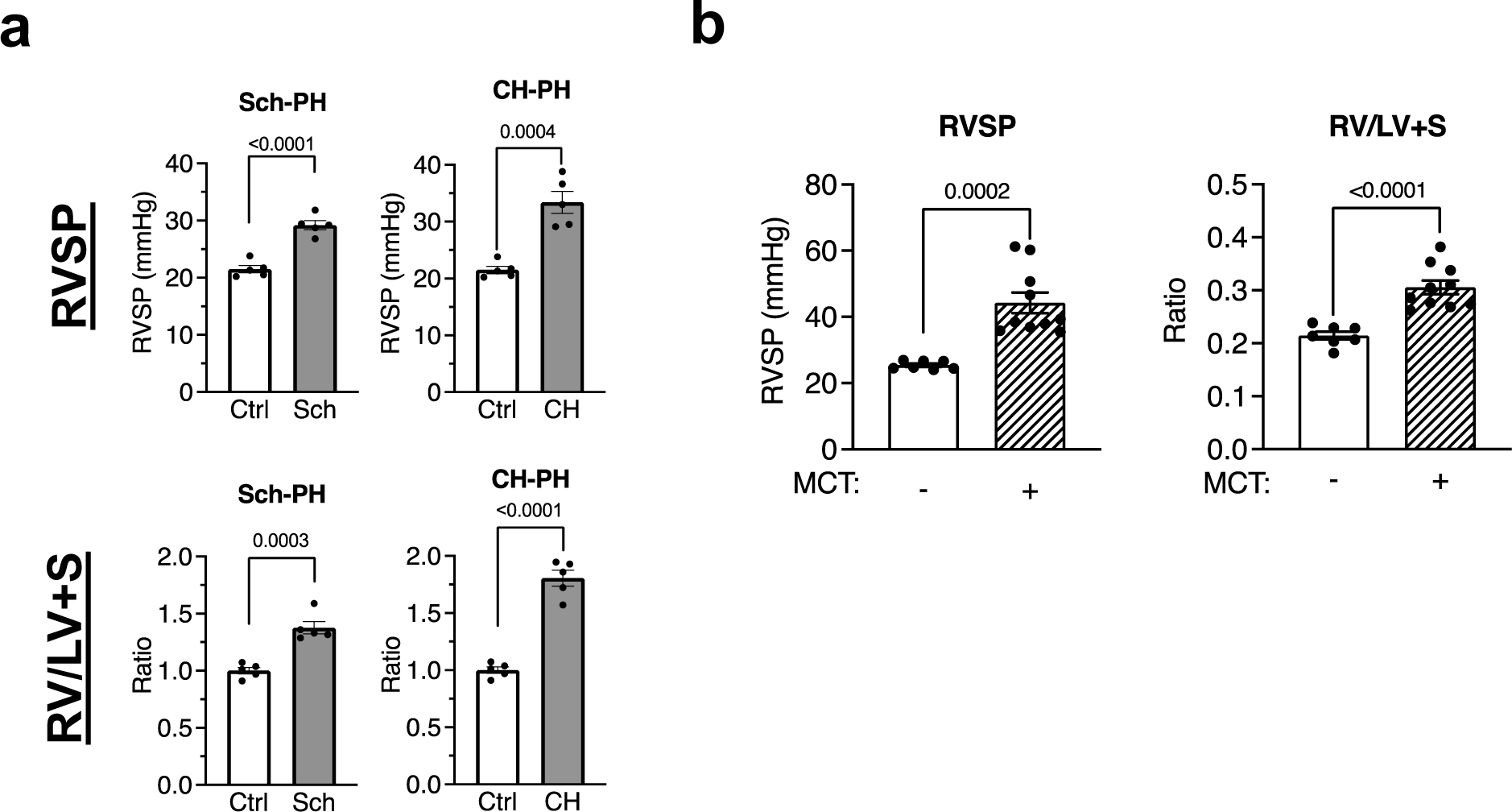
PH phenotypes in Schistosomia-induced PH (Sch-PH) and chronic hypoxia-induced PH (CH-PH), and Monocrotaline (MCT)-induced rats (MCT-PH) **a.** Right ventricular (RV) systolic pressure (RVSP, top) and RV to left ventricular (LV)+septum weight ratio (RV/LV+S, bottom) were measured in control, Sch-PH, and CH-PH mice and shown as mean±SEM. n=5 independent samples per group. **b.** The RVSP (left) and RV/LV+S ratio (right) in vehicle or MCT administered rats were measured and shown as mean±SEM. n=7∼10 independent samples per group.

**Supplementary Fig. S8.**
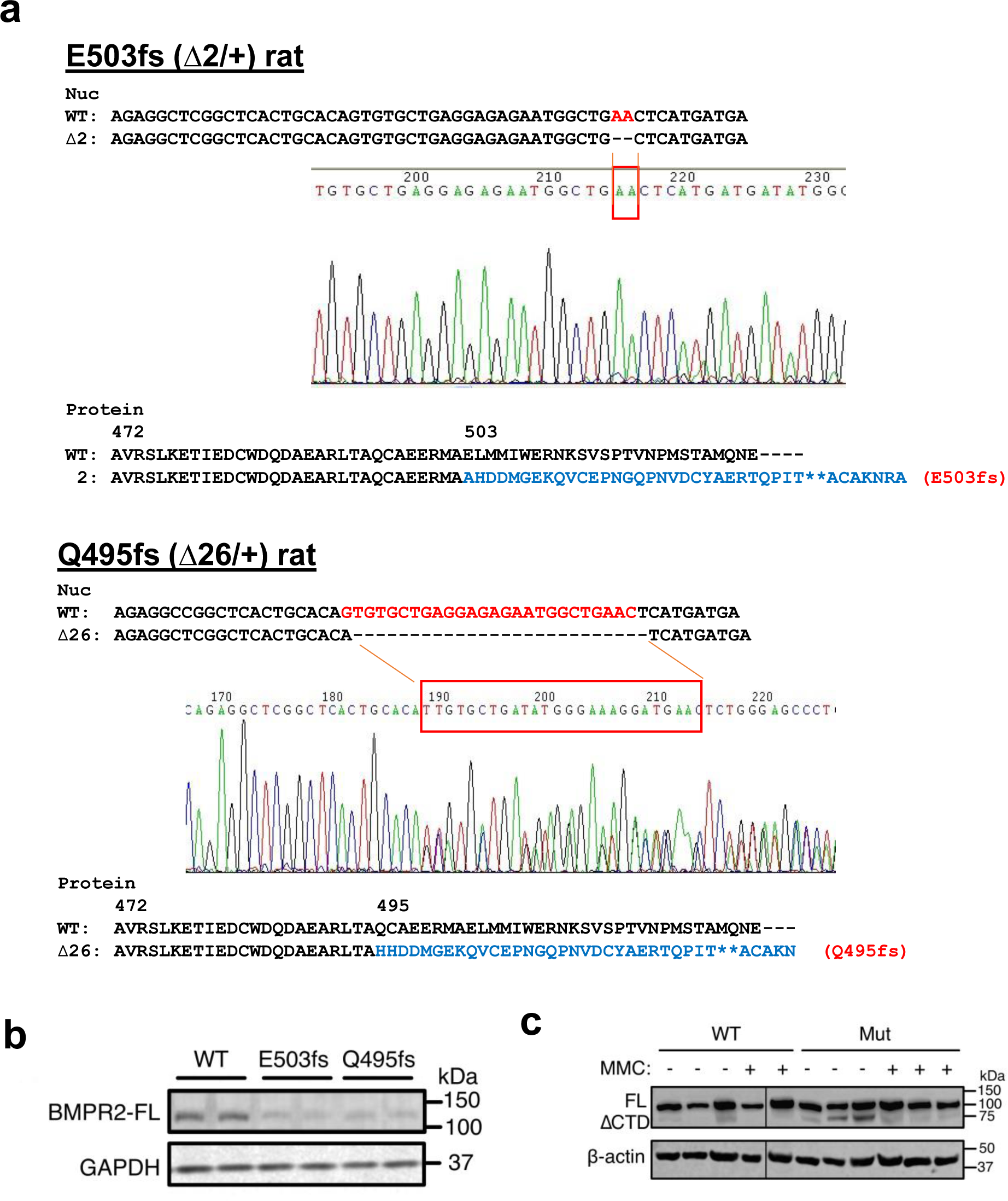
Generation of heterozygous *BMPR2* mutant rats (Q495fs and E503fs) **a.** The top 2 panels show BMPR2 gene sequencing result of E503fs (Δ2/+) rat and Q495fs (Δ26/+) rat, where the bases shown in red or in the red rectangle are mutations introduced. Blue letters (bottom) show protein sequence after the mutation. Asterisks indicate stop codons. **b.** Immunoblot using the antibody against BMPR2-CTD confirms the reduction of a full-length BMPR2 (BMPR2-FL) in the lung of Mut rats in comparison with WT rats. n=2 independent samples per condition. **c.** Immunoblot using the antibody against BMPR2 kinase domain detects BMPR2-FL in the liver of WT rats and BMPR2-FL and -ΔCTD in Mut rats. n=2∼3 independent samples per condition.

**Supplementary Fig. S9.**
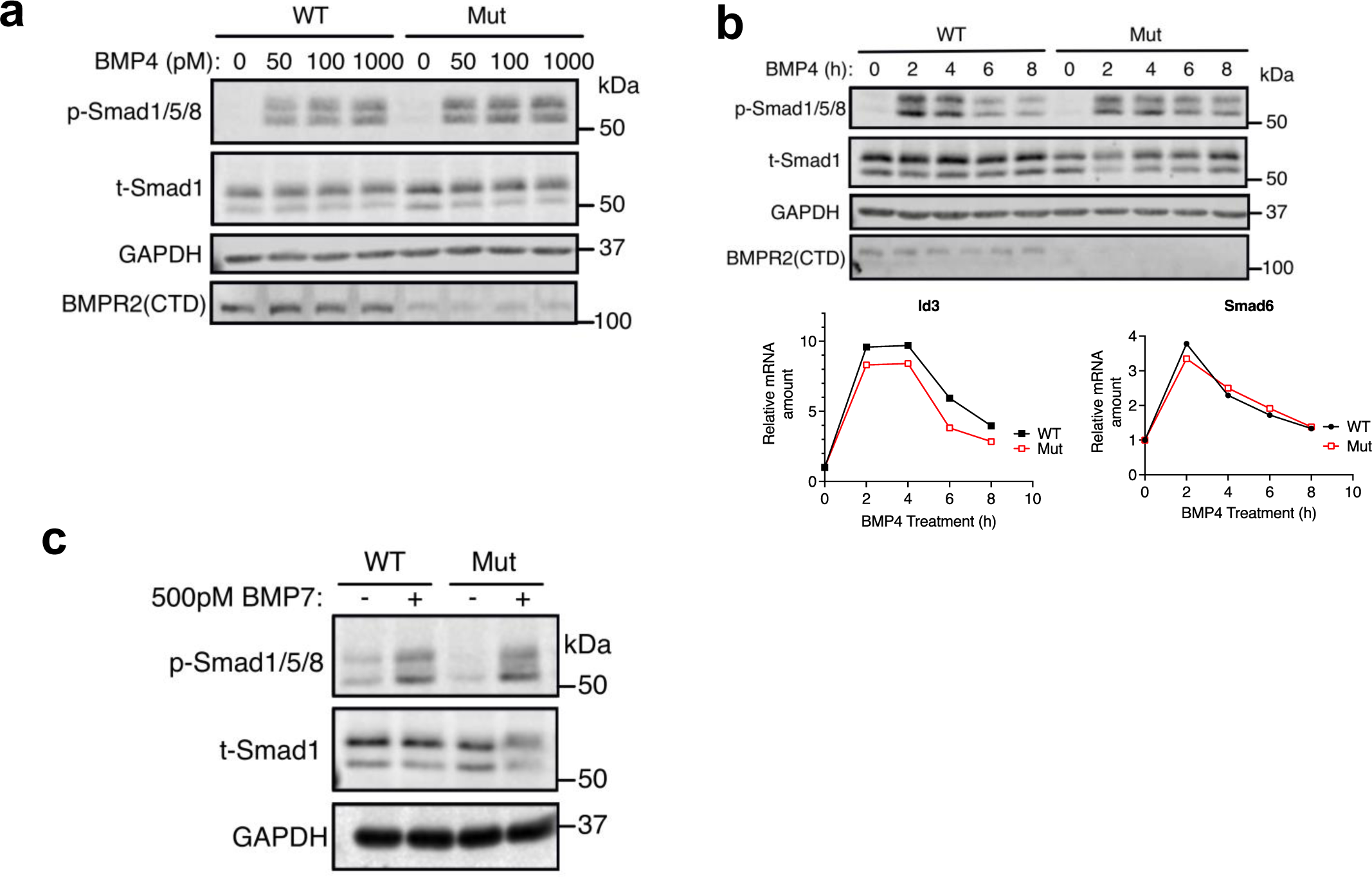
Vascular smooth muscle cells derived from WT and BMPR2-Mut (E503fs) rats respond to BMP4 and phosphorylate Smad1/5/8 in a similar manner. **a.** WT or Mut cells were treated with increasing concentrations of BMP4 and total cell lysates from 1×10^6^ cells were subjected to immunoblot with anti-phospho-Smad1/5/8 (p-Smad1/5/8), anti-total Smad1 (t-Smad1), anti-GAPDH (loading control), and anti-BMPR2(CTD) antibody. **b.** WT or Mut cells (1×10^6^ cells) were treated with 500 pM of BMP4 for indicated time (h) and total cell lysates were subjected to immunoblot with anti-p-Smad1/5/8, anti-t-Smad1, anti-GAPDH (loading control), and anti-BMPR2(CTD) antibody. as mean±SEM. Total RNAs isolated from the same cells were subjected to qRT-PCR of BMP-Smad1/5/8 target genes, Id3 and Smad6. Levels of Id3 and Smad6 mRNAs relative to GAPDH are shown by mean±SEM, n=3. **c.** WT or Mut cells (1×10^6^ cells) were treated with or without 500 pM BMP4 and total cell lysates were subjected to immunoblot with anti-p-Smad1/5/8, anti-t-Smad1, and anti-GAPDH (loading control) antibody.

**Supplementary Fig. S10.**
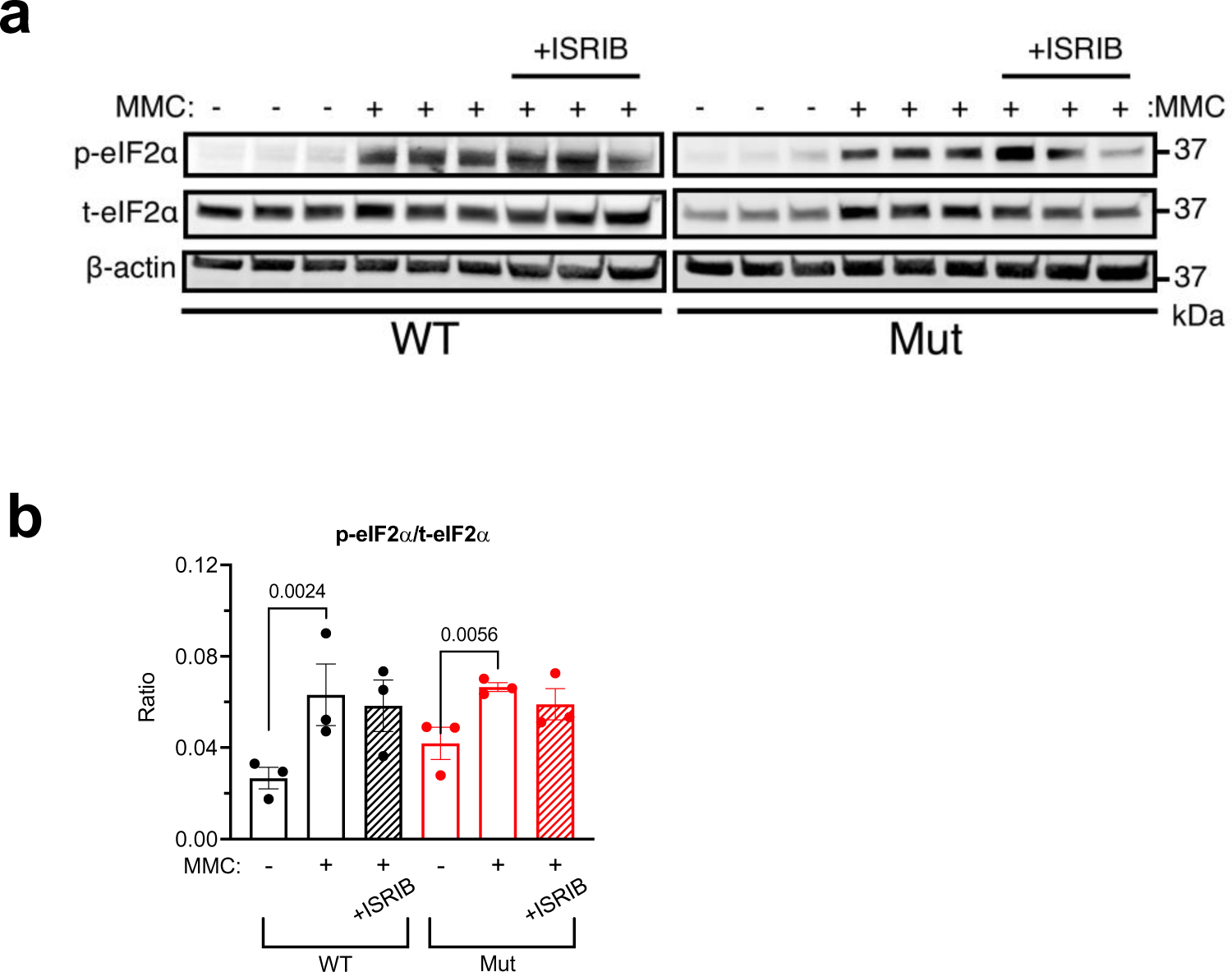
ISRIB does not affect phosphorylation status of eIF2α. **a.** Lung lysates from vehicle-, MMC-, or MMC+ISRIB-treated WT and Mut rats were subjected to immunoblot analysis of p-eIF2α, t-eIF2α, and β-actin (loading control) in triplicate. n=3 independent samples per condition. **b.** p-eIF2α/t-eIF2α ratio is shown as mean±SEM. n=3 independent samples per condition.

**Supplementary Fig. S11.**
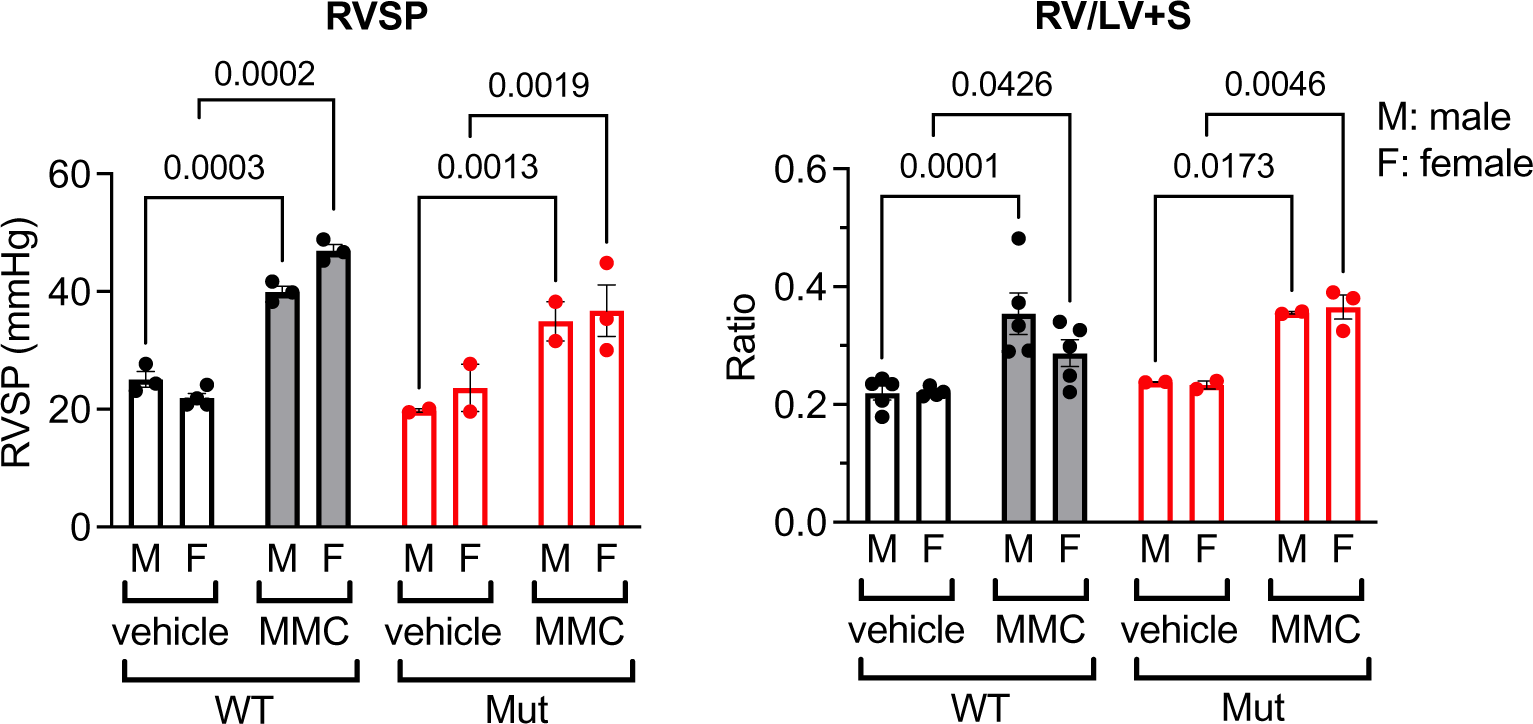
PH phenotypes induced by MMC in males and females are indistinguishable. RVSP (left) and RV/LV+S ratio (right) in female (orange) and male (blue) WT and Mut rats measured 24 days after the administration of vehicle of MMC and shown as mean±SEM. M and F stand male and female. n=3∼5 independent samples per condition.

**Supplementary Fig. S12.**
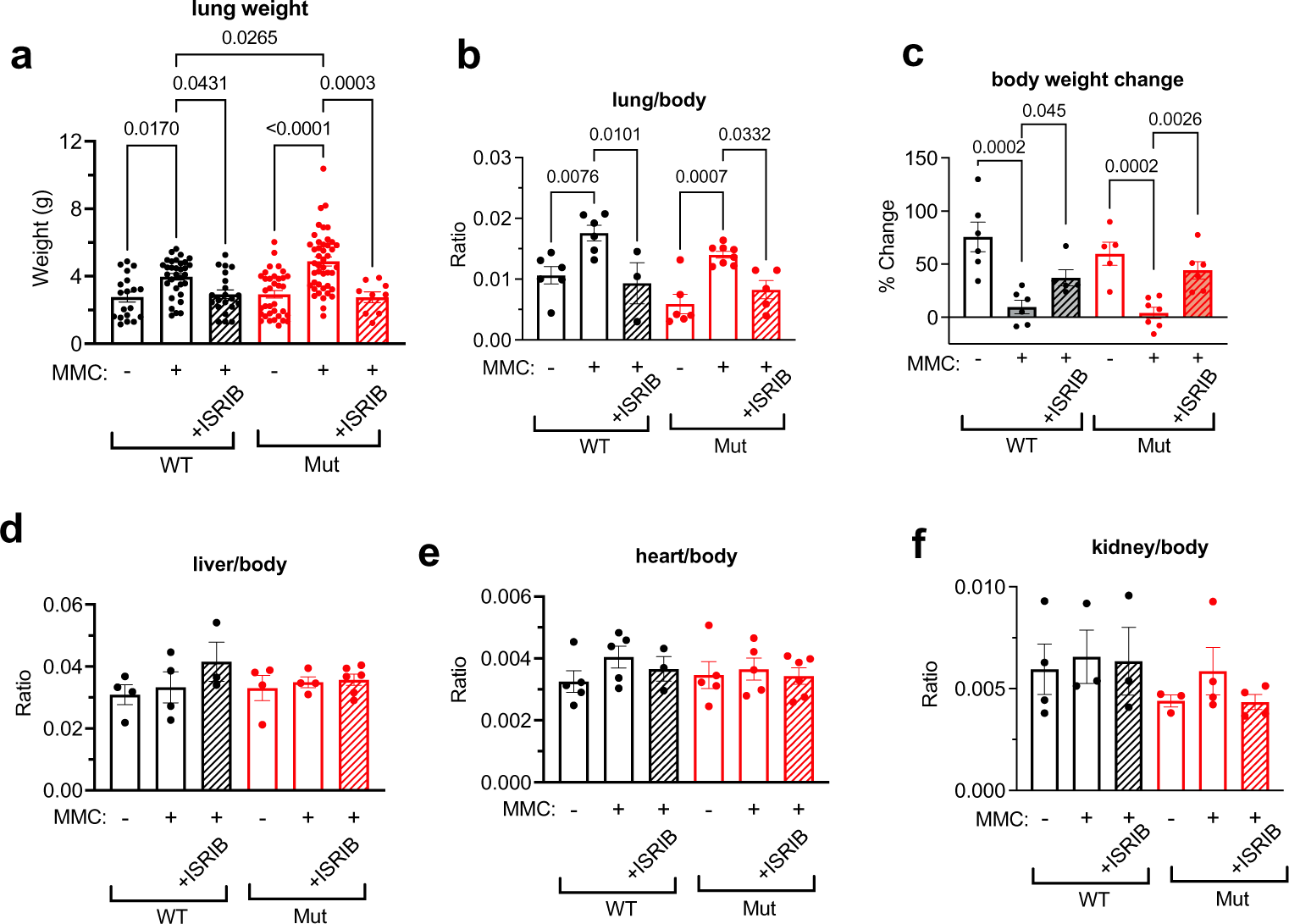
Physiological measurements of WT and Mut rats subjected to 24-days treatment of ISRIB. **a.** lung weight, **b.** lung weight/body weight ratio, **c.** % body weight change, **d.** liver weight/body weight ratio, **e**. heart weight/body weight ratio, and **f.** kidney weight/body weight ratio of WT or Mut rats subjected ISRIB (24-days treatment) are shown as mean±SEM. n=4∼44 independent samples per group

**Supplementary Fig. S13.**
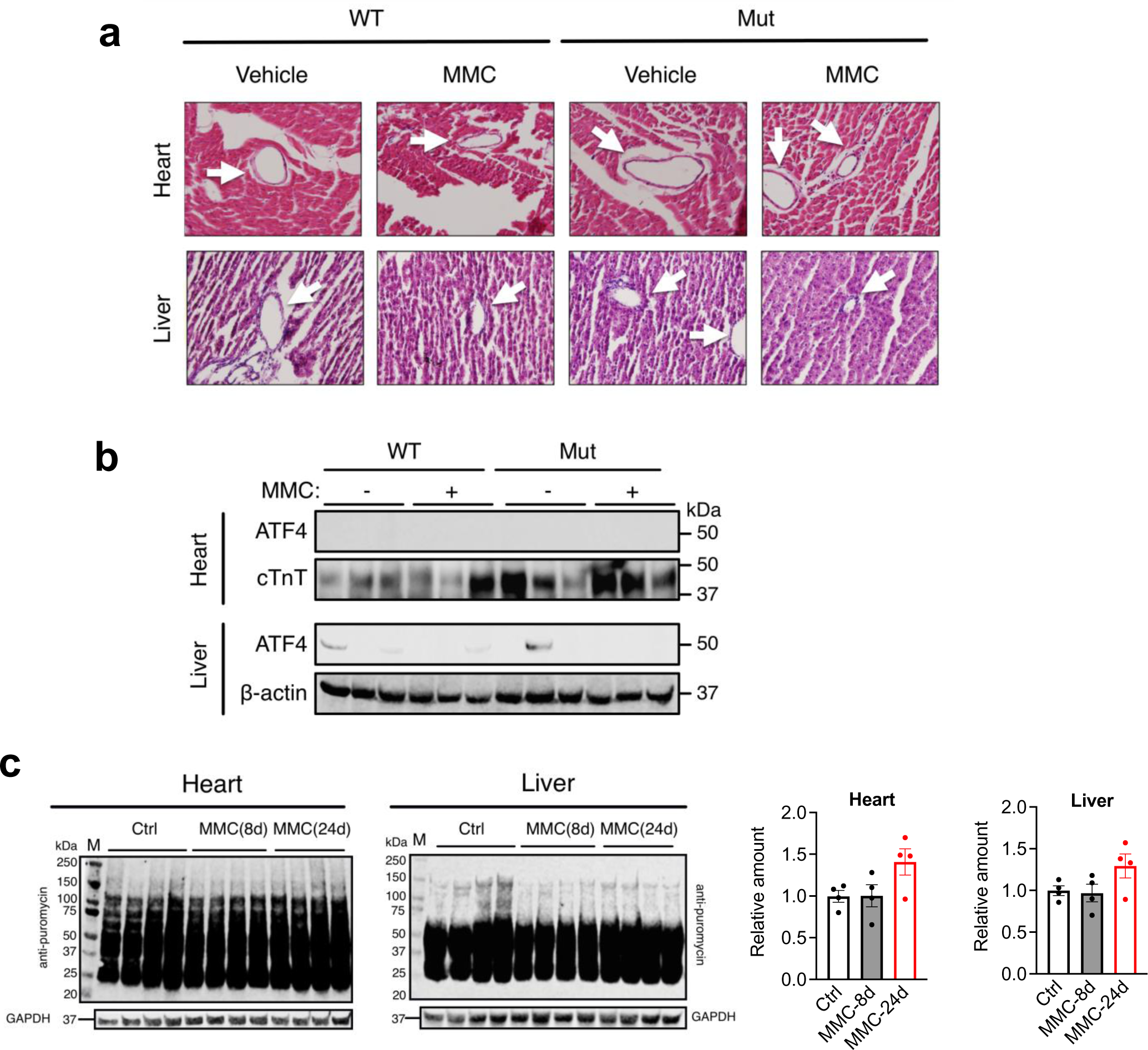
No vascular remodeling and no ISR activation in the heart and liver upon MMC administration in rats. **a.** Heart and lung were harvested from from WT rats 24 days after the injection of vehicle (saline) or MMC (3 mg/kg), followed by H&E staining. Arrows indicate vessels. **b.** Immunoblot analysis of ATF4, cardiac troponin T (cTnT; loading control for heart) and β-actin (loading control for liver) using heart and liver lysates harvested from WT and Mut rats 24 days after the treatment with vehicle of MMC. n=3 independent samples per group. **c.** Heart (left) and liver (right) lysates from WT rats treated with vehicle (24d) or MMC (8d or 24d) were subjected to anti-puromycin immunoblot and anti-GAPDH blot (loading control) (left). The relative levels of puromycin-labeled proteins normalized by GAPDH were quantitated and shown as mean±SEM. n=4 independent samples.

**Supplementary Fig. S14.**
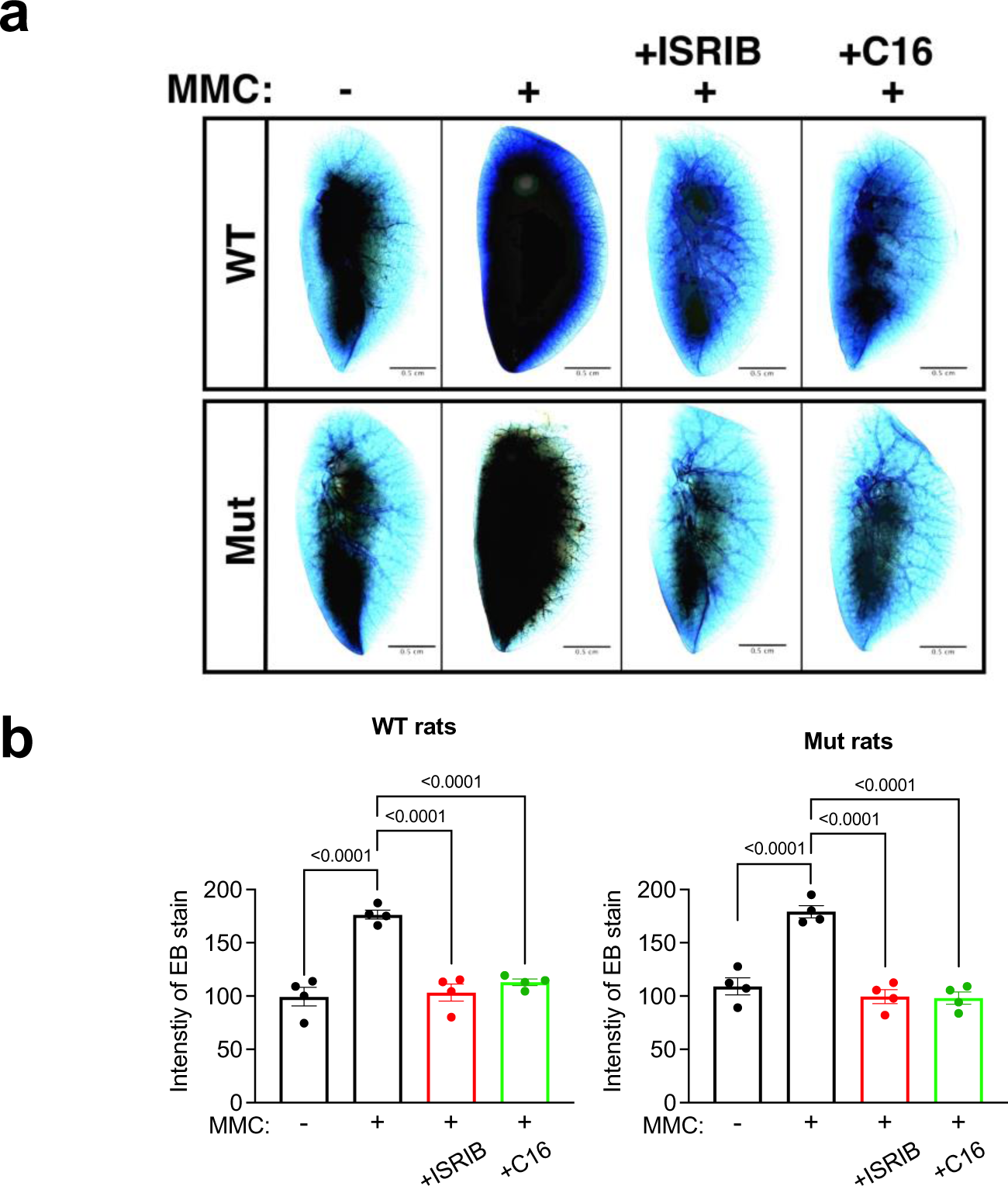
Examining the pulmonary vascular permeability in vivo. **a.** The permeability of pulmonary vasculature was assessed by injecting Evans Blue dye in WT and Mut rats administered with vehicle or MMC with or without ISRIB or C16. The lung was harvested on day 24 and the lung image was taken after the lung became translucent. Scale bar=0.5 cm. **b.** The relative intensity of EB staining of the lung in WT (left) and Mut rats (right) was quantitated by ImageJ and shown as mean±SEM. n=4 independent samples.

**Supplementary Fig. S15.**
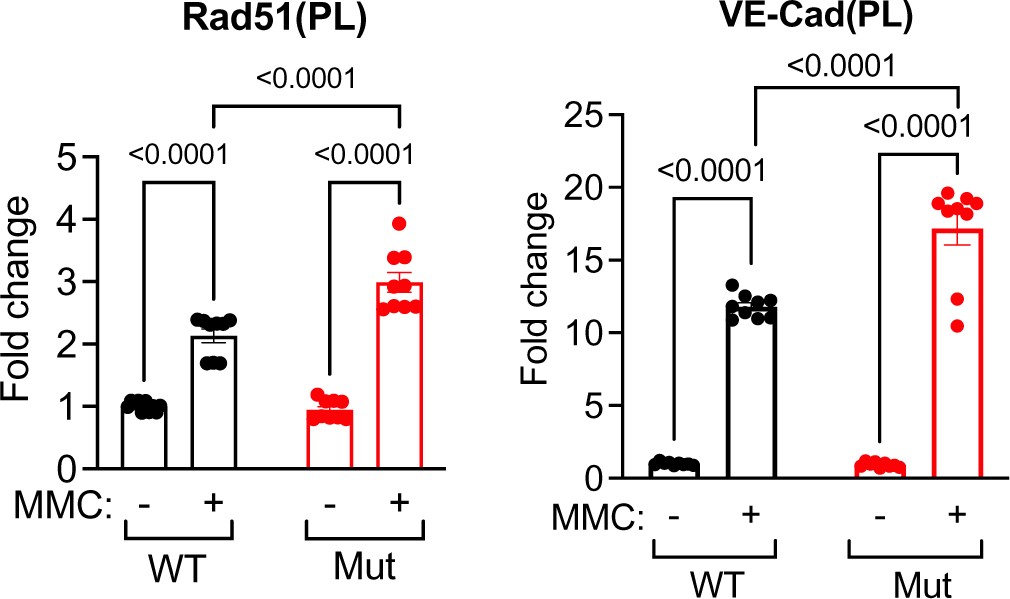
The amount of Rad51 and VE-Cad in the plasma of WT and Mut rats after MMC treatment. The amount of VE-Cad and Rad51 in the plasma of WT and Mut rats after MMC treatment (24d) were measured by ELISA and plotted as mean±SEM. n=9 independent samples per group.

**Supplementary Fig. S16.**
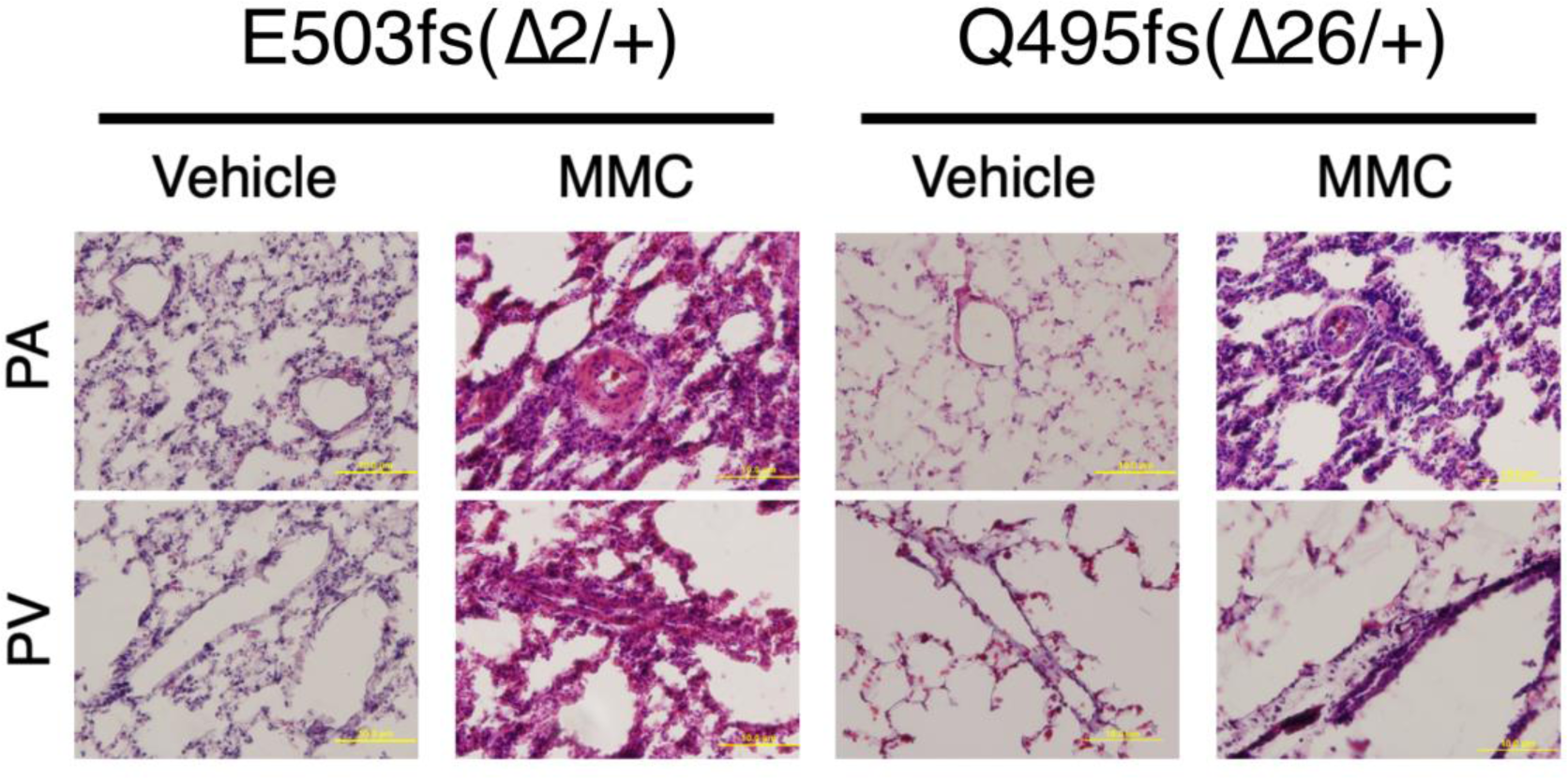
The vascular morphology of BMPR2 Mut lines E503fs (Δ2/+) rat and Q495fs (Δ26/+) rat after MMC treatment are indistinguishable. Representative image of pulmonary vasculature (H&E staining) exhibits similar phenotypes in E503fs (Δ2/+) and Q495fs)Δ26/+) rats 24 days after vehicle or MMC administration. Scale bar=10μm

**Supplementary Fig. S17.**
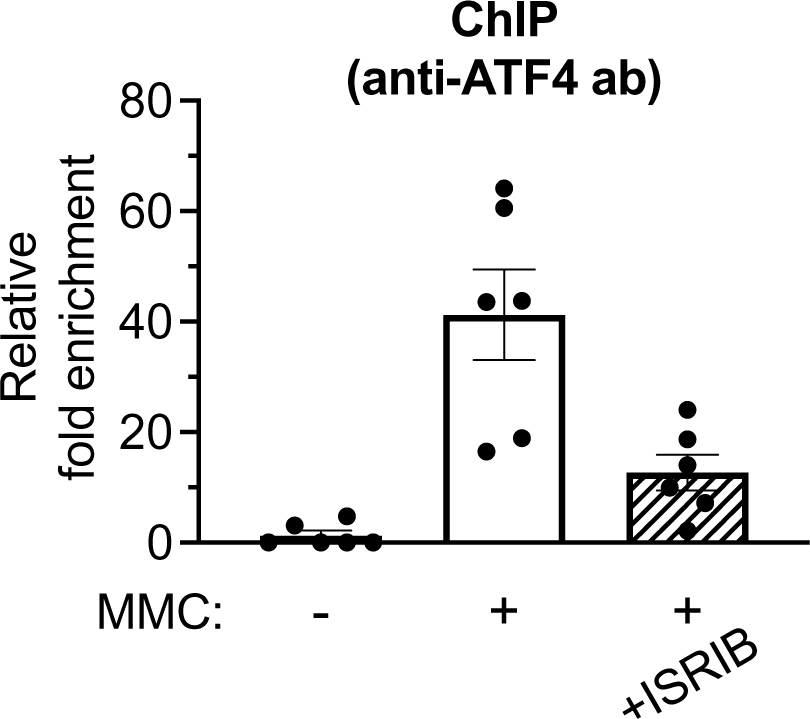
Recruitment of ATF4 to intron1 of the PKR gene upon MMC treatment is diminished by ISRIB. ChIP assay was performed in PMVECs (1×10^6^ cells) treated with vehicle, MMC or MMC+ISRIB for 4 h with anti-ATF4 antibody, followed by PCR amplification of intron 1 of the PKR gene that contains three ATF4 consensus binding motifs. The result is shown as fold enrichment over the input sample as mean±SEM. n=6 independent samples.

**Supplementary Fig. S18.**
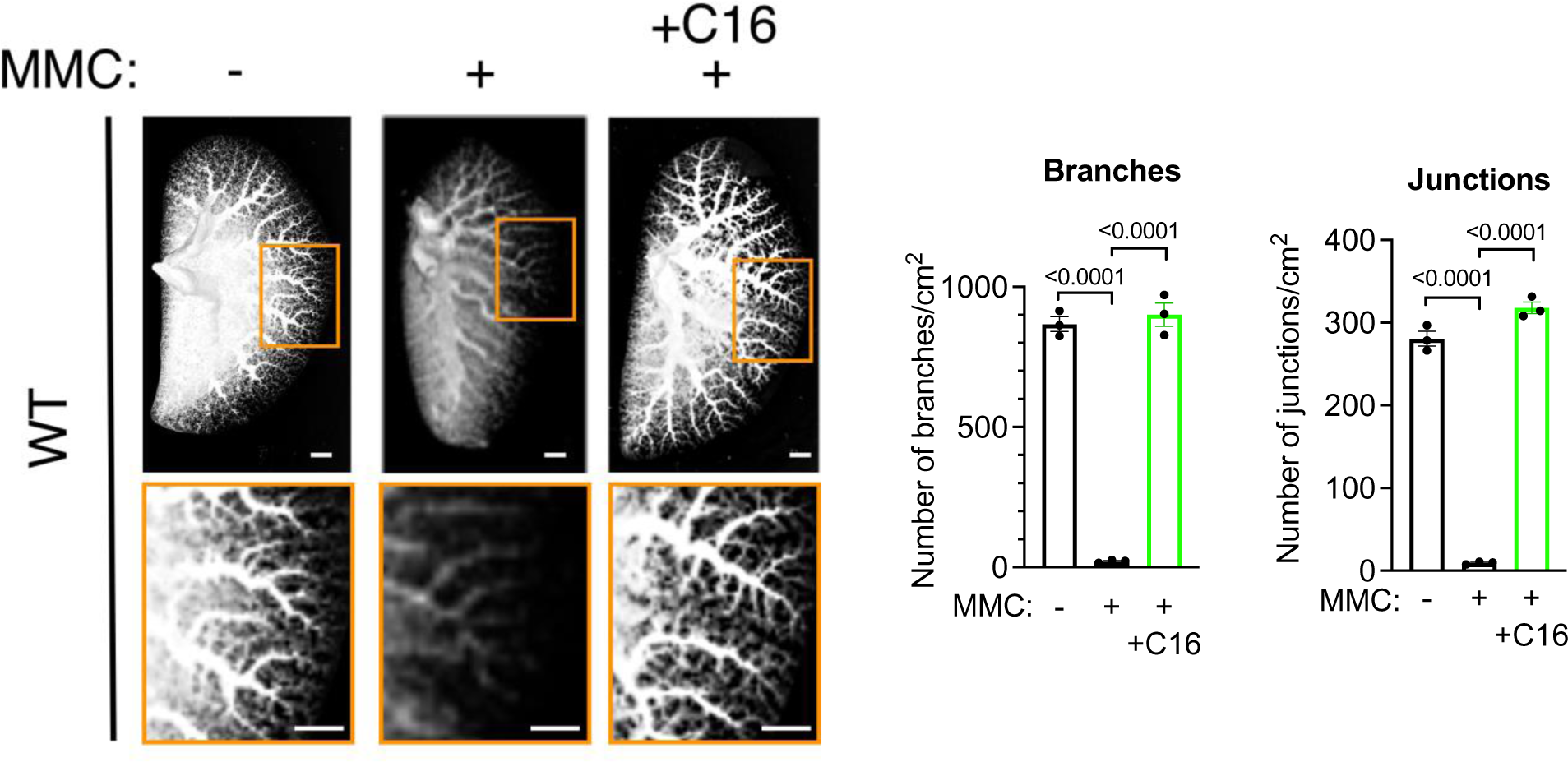
Administration of C16 prevents reduction of the distal pulmonary vessel density mediated by MMC. The pulmonary vessels in WT rats treated with vehicle or MMC with or without C16 were casted with Microfil. After harvesting the lung, the image of the lobe was taken and presented after converting it to the black and white image. Distal pulmonary vessels in the area indicated by orange box (top) are magnified and shown below (left). Scale bar=0.5 cm. A number of branches and junctions (per cm^2^) of distal pulmonary vessels was counted and shown as mean±SEM (right). n=3 independent samples per group.

**Supplementary Fig. S19.**
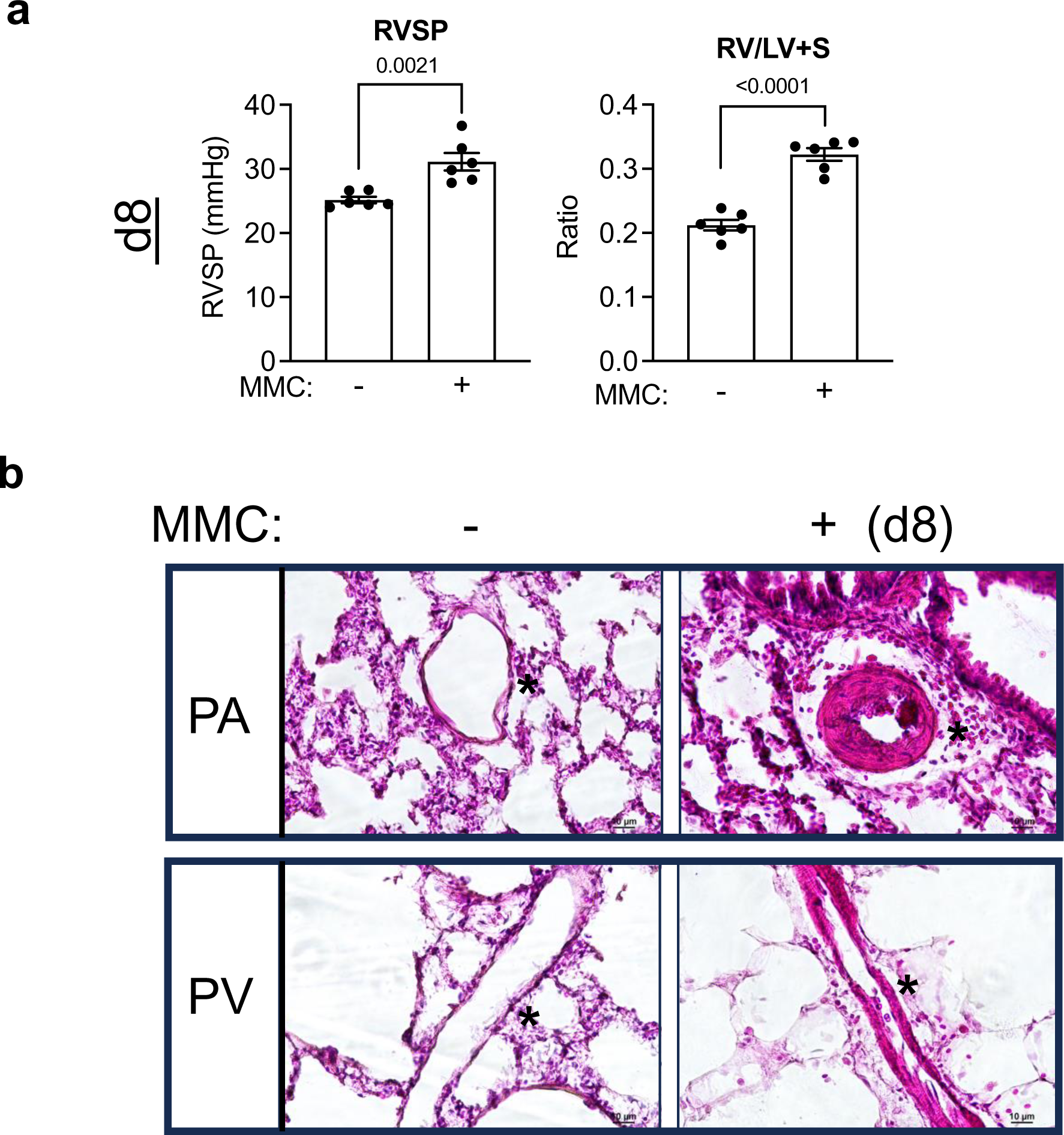
PVRD phenotypes are evident as early as 8 days after the administration of MMC. **a.** RVSP and RV/LV+S ration in WT rats was measured on 8 days after the administration of vehicle or MMC and shown as mean±SEM. n=6 independent samples. **b.** To examine the morphology of the pulmonary arteries (PAs) and veins (PVs), lungs were harvested from WT rats 8 days after the administration of vehicle or MMC and subjected to H&E staining. Asterisk indicates the location of vessels. Scale bar=10μm.

**Supplementary Fig. S20.**
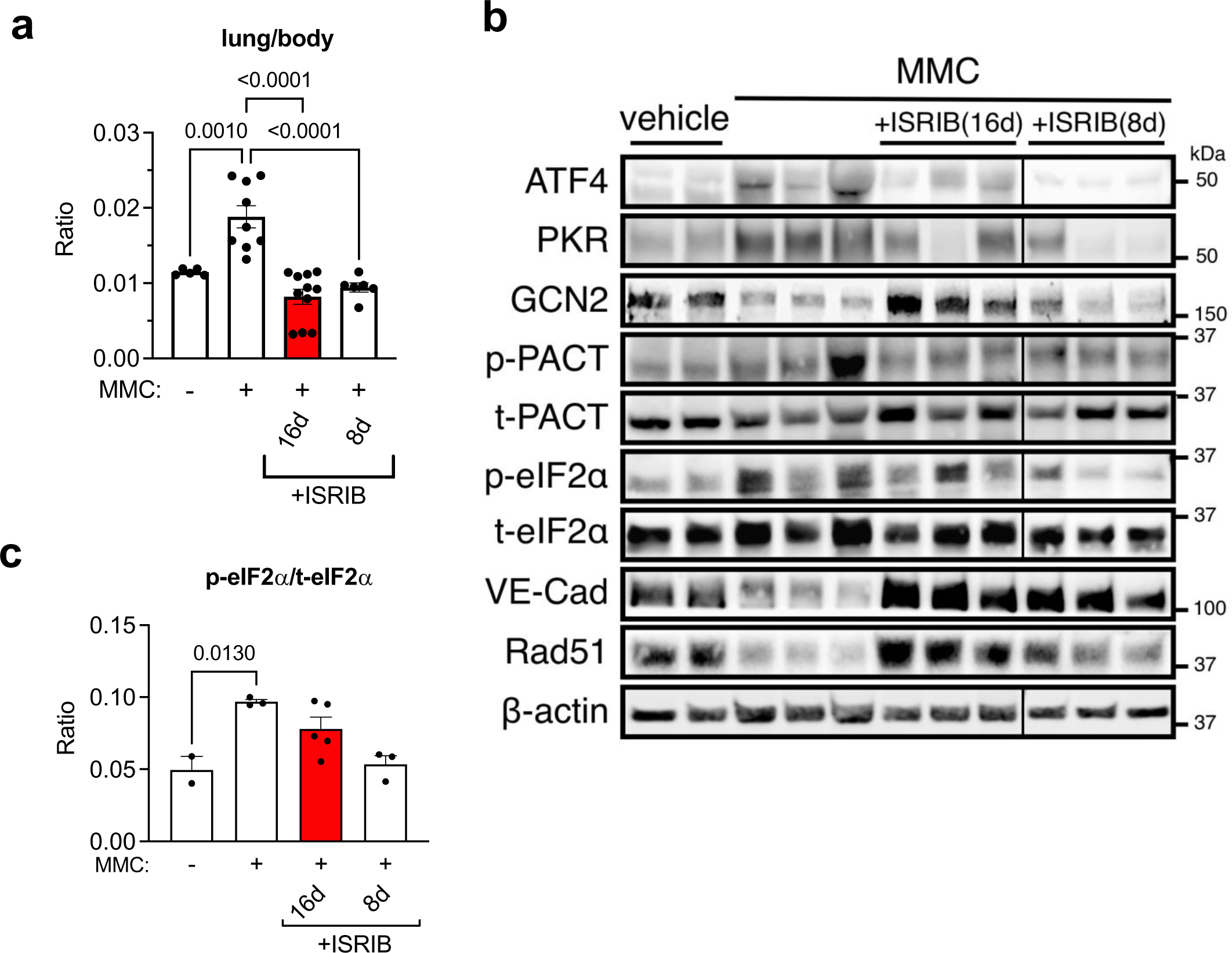
Delayed treatment with ISRIB attenuates ISR activation and reverses MMC-induced vascular phenotypes. The lung weight/body weight ratio **a.** immunofluorescence staining VE-Cad, Rad51, and DAPI. n=3∼11 independent samples per condition. **b.** WT rats administered with vehicle or MMC with or without 16-day (16d) or 8-day (8d) treatment with ISRIB was harvested on day 24, and lung lysates were subjected with immunoblot of indicated proteins. β- actin is loading control. Quantitation of these blots are shown in Fig. 8e. **c.** Immunoblots of lung lysates of vehicle- or MMC-treated WT rats with or without 16-day or 8-day treatment with ISRIB with antibodies against p-eIF2α and t-eIF2α and p-eIF2α/t-eIF2α ratio was calculate and shown as mean±SEM. n=2∼3 independent samples per condition.

**Supplementary Fig. S21.**
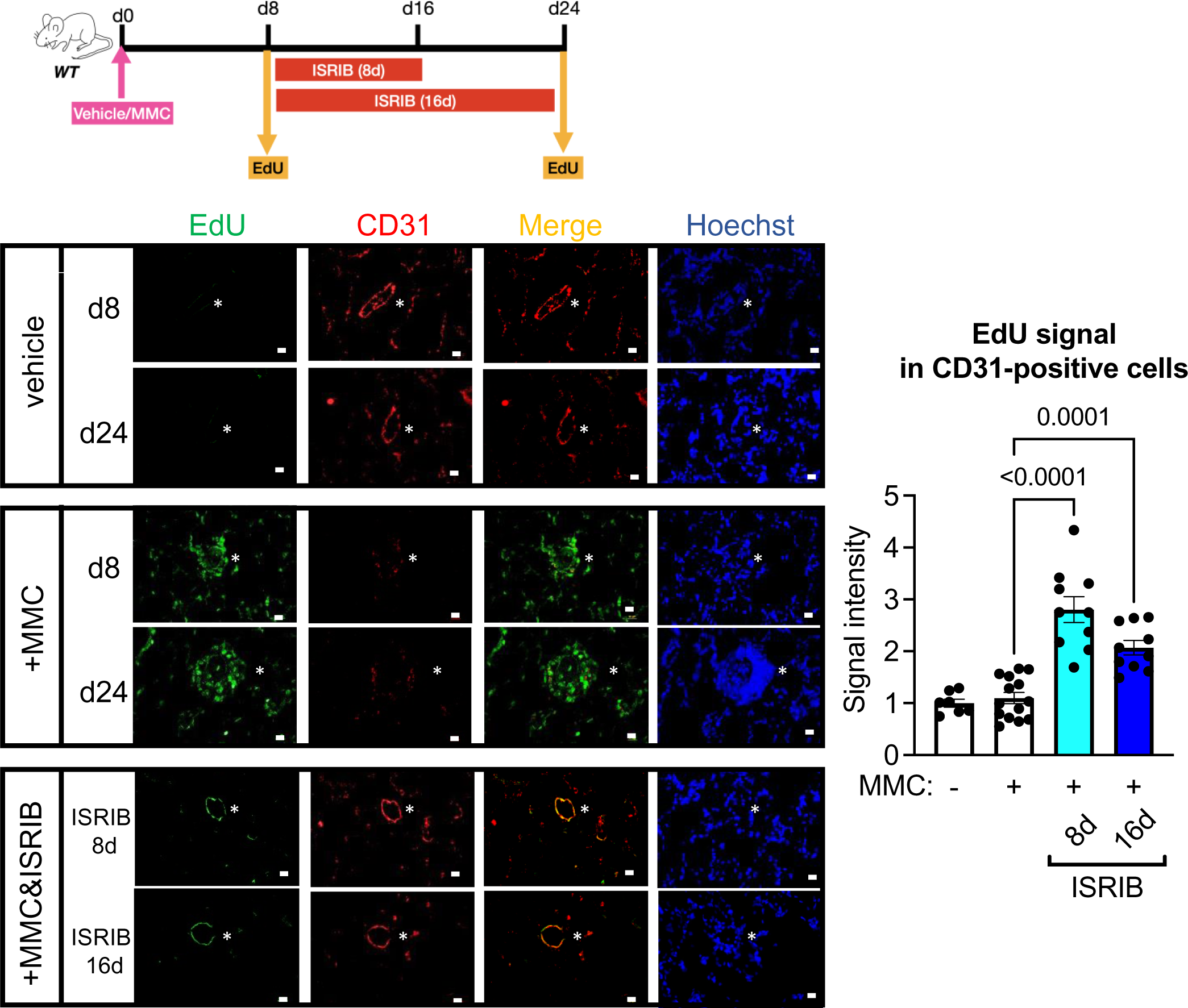
ISRIB promotes the proliferation on vascular endothelial cells in the MMC-treated rats. Scheme of delayed ISRIB treatment and EdU staining (16d and 8d treatment) (top left). WT rats were administered with vehicle or MMC on day 0 and treated with ISRIB starting on day 8 for 8 days (8d) or 16 days (16d). The EdU staining (green) was performed on day 8 (d8) or day 24 (d24). The lung section was co-stained with the endothelial marker CD31 (red) and Hoechst (blue; for nuclei staining). The images of EdU and CD31 stain were merged and shown in the “Merge” row (bottom left). The intensity of EdU signal in the CD31-positive vascular endothelial layer was quantitated by ImageJ and shown as mean±SEM (right). n=7∼14 independent samples per condition. White asterisks indicate pulmonary arteries. Scale bar= 10 μm

**Supplementary Fig. S22.**
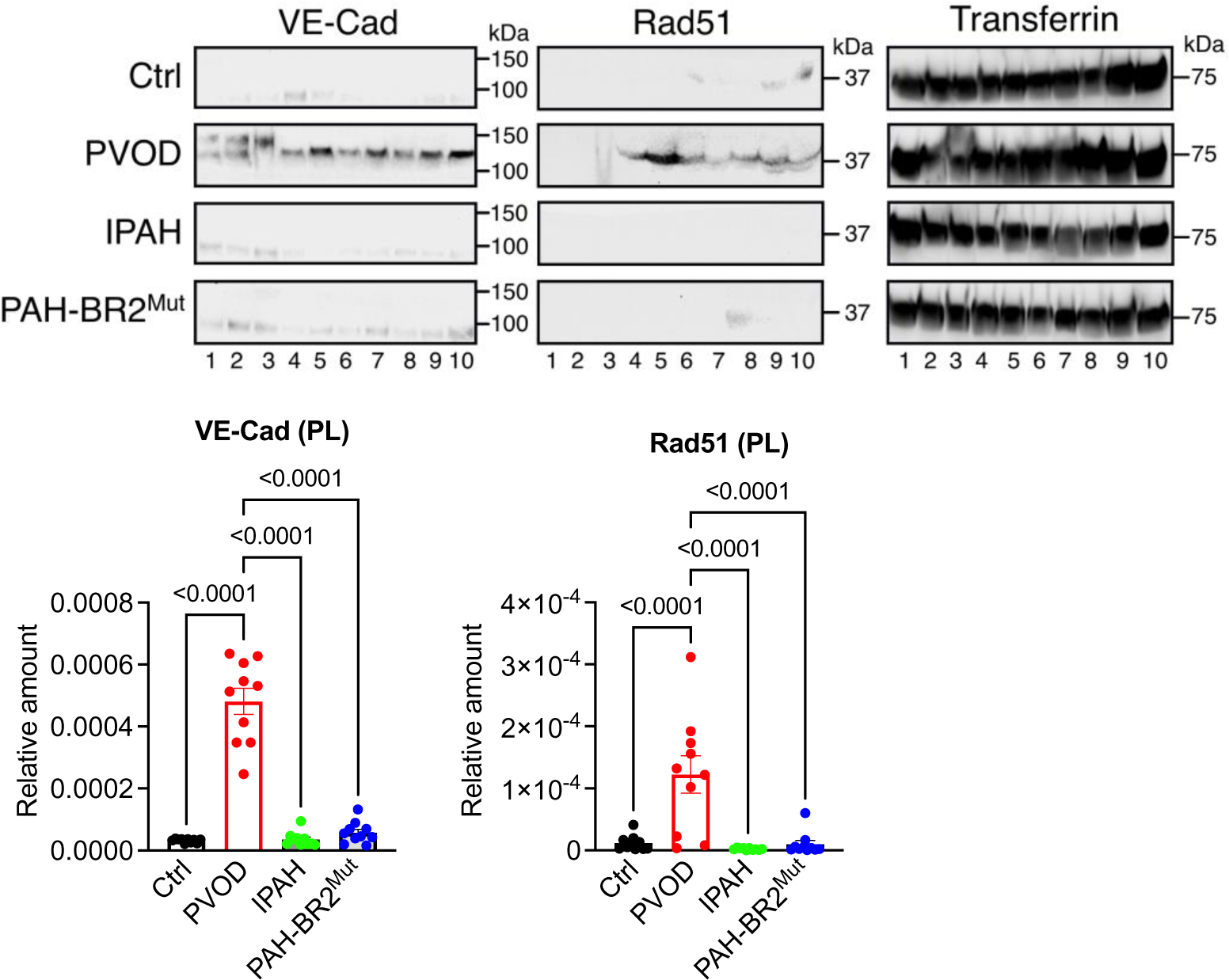
Release of VE-Cad and Rad51 into the plasma in patients with PVOD. Human patients with PVOD, IPAH, and PAH with BMPR2 mutations (PAH-BR2^Mut^) and non-PAH control individuals (n=10 per group) were subjected to immunoblot with anti-VE-Cad, anti-Rad51, and anti-Transferrin (loading control) antibody (top). #1-10 indicate 10 samples from individuals. The relative amounts of VE-Cad and Rad51 normalized by transferrin are quantitated and shown as mean±SEM (bottom). n=10 independent samples per condition.

**Supplementary Fig. S23.**
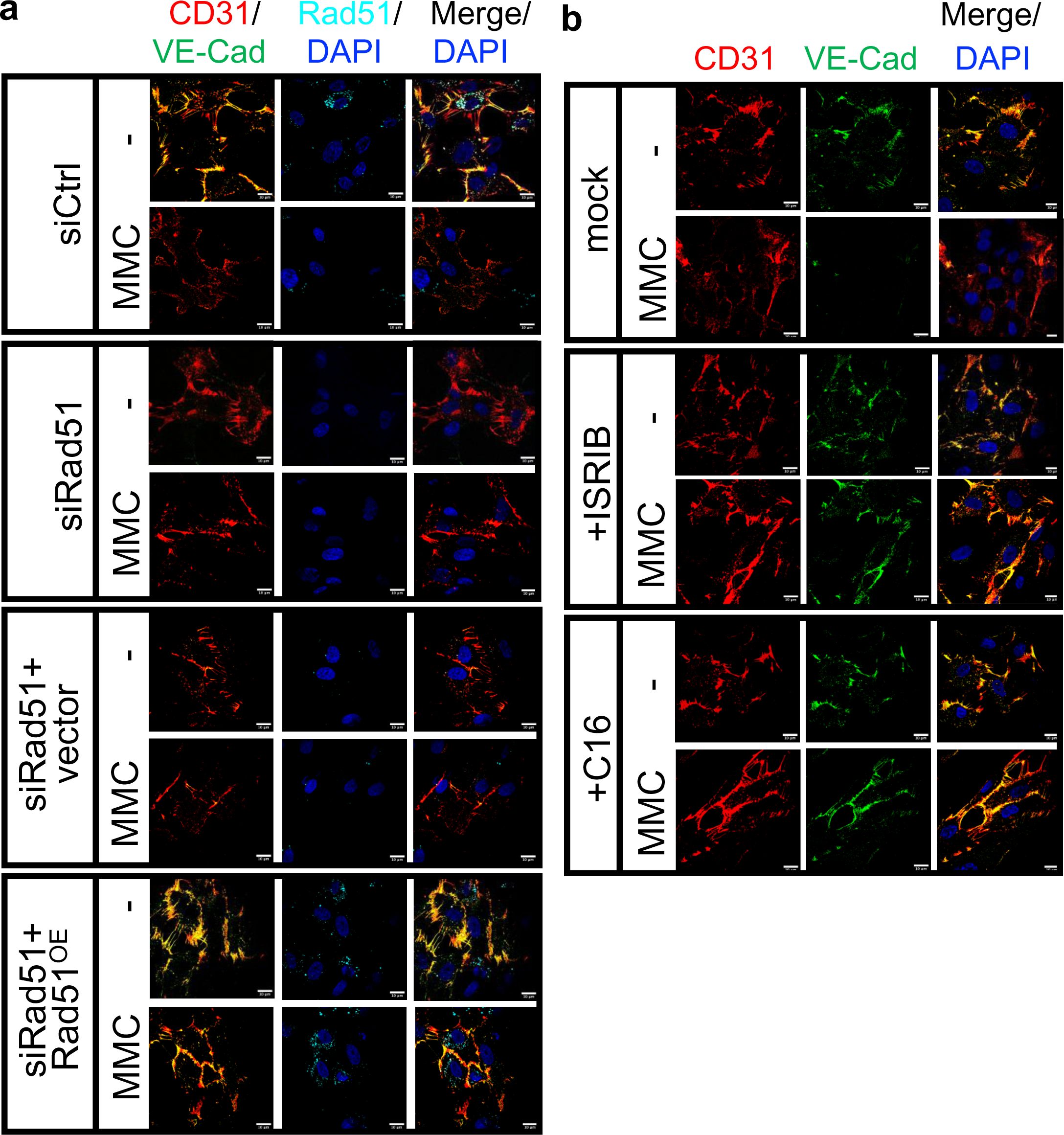
The morphology of PMVECs in the presence or absence of Rad51. **a.** PMVECs (5×10^3^ cells) were treated with vehicle or MMC for 14 h after the transfection of Control siRNA (siCtrl) or siRNA against Rad51 (siRad51) with Rad51 overexpression (Rad51^OE^) or the transfection of empty vector (vector) as control. Twenty-four hour after the transfection, cells were subjected to the immunofluorescence staining of CD31 (red), VE-Cad (green) and Rad51 (cyan). DAPI stain (blue) for nuclei. Scale bar=10μm **b.** PMVECs (5×10^3^ cells) were treated with vehicle or MMC for 14 h with vehicle, ISRIB, or C16, and subjected to the immunofluorescence staining of CD31 (red) and VE-Cad (green). DAPI stain (blue) for nuclei. Scale bar=10μm

**Supplementary Fig. S24.**
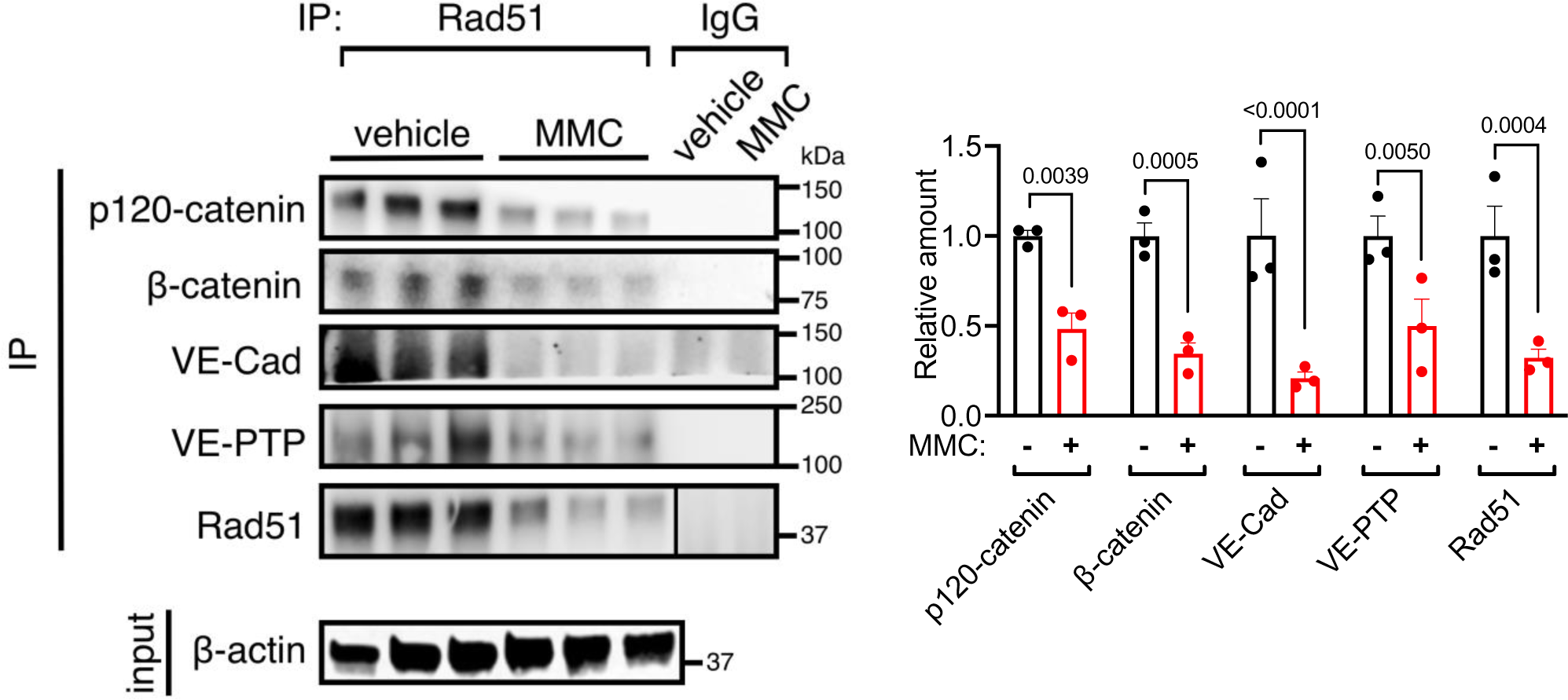
β-catenin, VE-PTP, and p120-catnin associates with VE-Cadherin that is in the complex with Rad51. PMVECs were treated with vehicle or MMC for 14 h prior to the immunoprecipitation (IP) with an anti-Rad51 antibody, followed by immunoblot analysis of p120-catenin, β-catenin, VE-Cad, VE-PTP, and Rad51 in triplicates (left). The total cell lysates without IP were subjected to immunoblot with β-actin (loading control). The amounts of p120-catenin, β-catenin, VE-Cad, VE-PTP, and Rad51 relative to β-actin were quantitated and shown as mean±SEM (right). n=3 independent samples.

